# Pathogenic mutations of human phosphorylation sites affect protein-protein interactions

**DOI:** 10.1101/2023.08.01.551433

**Authors:** Trendelina Rrustemi, Katrina Meyer, Yvette Roske, Bora Uyar, Altuna Akalin, Koshi Imami, Yasushi Ishihama, Oliver Daumke, Matthias Selbach

## Abstract

Despite their lack of a defined 3D structure, intrinsically disordered regions (IDRs) of proteins play important biological roles. Many IDRs contain short linear motifs (SLiMs) that mediate protein-protein interactions (PPIs), which can be regulated by post-translational modifications like phosphorylation. 20% of pathogenic missense mutations are found in IDRs, and understanding how such mutations affect PPIs is essential for unraveling disease mechanisms. Here, we employed peptide-based interaction proteomics to investigate 36 disease-causing mutations affecting phosphorylation sites. Our results unveiled significant differences in interactomes between phosphorylated and non-phosphorylated peptides, often due to disrupted phosphorylation-dependent SLiMs. We focused on a mutation of a serine phosphorylation site in the transcription factor GATAD1, which causes dilated cardiomyopathy. We found that this phosphorylation site mediates interaction with 14-3-3 family proteins. Follow-up experiments revealed the structural basis of this interaction and suggest that 14-3-3 binding affects GATAD1 nucleocytoplasmic transport by masking a nuclear localisation signal. Our results demonstrate that pathogenic mutations of human phosphorylation sites can significantly impact protein-protein interactions, offering fresh insights into potential molecular mechanisms underlying pathogenesis.

## Introduction

Understanding protein function in health and disease is a key challenge in the post-genomic era (Eisenberg *et al*, 2000). “Omics” techniques provide a wealth of data, but mechanistic understanding is often lagging behind. For example, although sequencing technologies have identified numerous single amino acid variants (SAVs), their functional implications remain mostly unknown, even when they have been linked to disease (Backwell & Marsh, 2022; Lek *et al*, 2016; Wright *et al*, 2015). Proteins are also modified by posttranslational modifications (PTMs) such as phosphorylation, and proteomics can now routinely identify tens of thousands of phosphorylation sites (Riley & Coon, 2016; Bekker-Jensen *et al*, 2020a; Kitata *et al*, 2021; Skowronek *et al*, 2022). However, most sites have no known kinase or biological function (Needham *et al*, 2019). Hence, while genomic and proteomic technologies provide abundant information about SAVs and PTMs, respectively, how these changes affect protein function remains largely unexplored.

The classical sequence-structure-function paradigm posits that amino acid sequences determine protein structure and therefore protein function. Accordingly, the impact of SAVs and PTMs on protein function is often investigated from a structural angle. However, around 40% of the proteome consists of intrinsically disordered regions (IDRs) that have low sequence complexity (Uversky, 2019), and over 20% of known disease-causing and cancer driver mutations affect amino acid residues in IDRs (Vacic *et al*, 2012; Mészáros *et al*, 2021). Amino acids in IDRs are also frequently modified by PTMs (Darling & Uversky, 2018; Bah & Forman-Kay, 2016; Bludau *et al*, 2022). Due to the lacking structure-function relationship, understanding how SAVs and PTMs in IDRs affect protein function is especially challenging.

It is now well established that IDRs are critically involved in virtually every cellular process (Wright & Dyson, 2015). They achieve this in a number of different ways such as the induction of structural changes in adjacent structured regions, transitioning from disorder-to-order, and/or by mediating protein-protein interactions (Dunker *et al*, 2005; Dyson & Wright, 2005; Bugge *et al*, 2020). Protein-protein interactions in IDRs are mediated by so-called short linear motifs (SLiMs) – sequence stretches shorter than ten amino acids with simple specificity determinants that are recognized by cognate domains in interacting proteins (Tompa *et al*, 2014; Davey *et al*, 2023). Importantly, many SLiM-mediated interactions are dynamically regulated by PTMs, and the dynamic interplay between specific PTMs (e.g. tyrosine phosphorylation) and recruitment of protein “readers” with cognate domains (e.g. SH2 domains) plays a pivotal role in cell signaling (Seet *et al*, 2006). In fact, more than 20% of validated SLiMs in the Eukaryotic Linear Motif (ELM) database are post-translationally modified, with phosphorylation being the most common modification type (Davey *et al*, 2012).

A number of different experimental approaches to map the SLiM-based interactome have been established (Davey *et al*, 2023). One such approach is peptide-based interaction proteomics (reviewed by (Meyer & Selbach, 2020)). It employs synthetic peptides corresponding to IDRs of interest that are used to pull-down interacting proteins from complex protein lysates. In combination with the parallel synthesis of peptides on cellulose membranes (SPOT synthesis) (Frank, 1992), the throughput of peptide-based interaction proteomics can be greatly increased (Hernandez & Dittmar, 2021; Meyer & Selbach, 2020; Ramberger *et al*, 2021b). This “Protein Interaction Screen on Peptide Matrix” (PRISMA) method has been applied in a number of recent studies (Ramberger *et al*, 2021a; Kassa *et al*, 2023; Meyer *et al*, 2018). For example, we used PRISMA to study how pathogenic mutations in IDRs affect protein-protein interactions (PPIs). This revealed that a pathogenic point mutation in the glucose transporter GLUT1 causes GLUT1 deficiency syndrome by creating a SLiM that recruits adaptor proteins and mediates GLUT1 endocytosis (Meyer *et al*, 2018). A key advantage of peptide pulldown approaches is that the peptides can be synthesized in modified forms, enabling direct assessment of the impact of PTMs on PPIs (Schulze & Mann, 2004; Lundby *et al*, 2019; Vermeulen *et al*, 2010; Selbach *et al*, 2009; Ramberger *et al*, 2021b, 2021a).

Disease-associated SAVs in IDRs are enriched at interaction interfaces, supporting the view that many mutations in IDRs cause disease by affecting PPIs (Wong *et al*, 2020). However, if and how these perturbed interactions also involve PTMs is not well understood. Since both disease-causing SAVs and PTMs can affect PPIs, we reasoned that studying SAVs that affect known phosphorylation would be particularly interesting. Here, we investigated the interplay of disease-associated SAVs and PTMs on protein-protein interactions. Specifically, we selected SAVs affecting known phosphorylation sites and asked if the SAV and/or the phosphorylation state of the site changes protein-protein interactions. To this end, we used PRISMA to directly compare the interactome of the wild-type, mutated and phosphorylated site.

## Results

### A peptide-based screen for interactions affected by phosphorylation and disease-causing SAVs

To assess how disease-causing SAVs of protein phosphorylation sites affect protein-protein interactions, we first selected pathogenic missense mutations of known serine, threonine or tyrosine phosphorylation sites. To this end, we used a dataset from PhosphositePlus that maps posttranslational modification sites to disease-associated genetic variants (Hornbeck *et al*, 2015). At the time, this dataset included 33,359 entries, including 12,658 disease-causing mutations from various databases (COSMIC, Uniprot Humsavar, TCGA, cBio) and seven types of posttranslational modifications. After filtering for mutations in intrinsically disordered regions, we obtained 1,965 mutations, of which 126 directly affected the phosphorylated amino acid. We then selected mutations that had no other annotated modifications within seven amino acids of the target residue. This filtering resulted in a final set of 38 disease candidates (Figure 1A).

**Figure 1.**
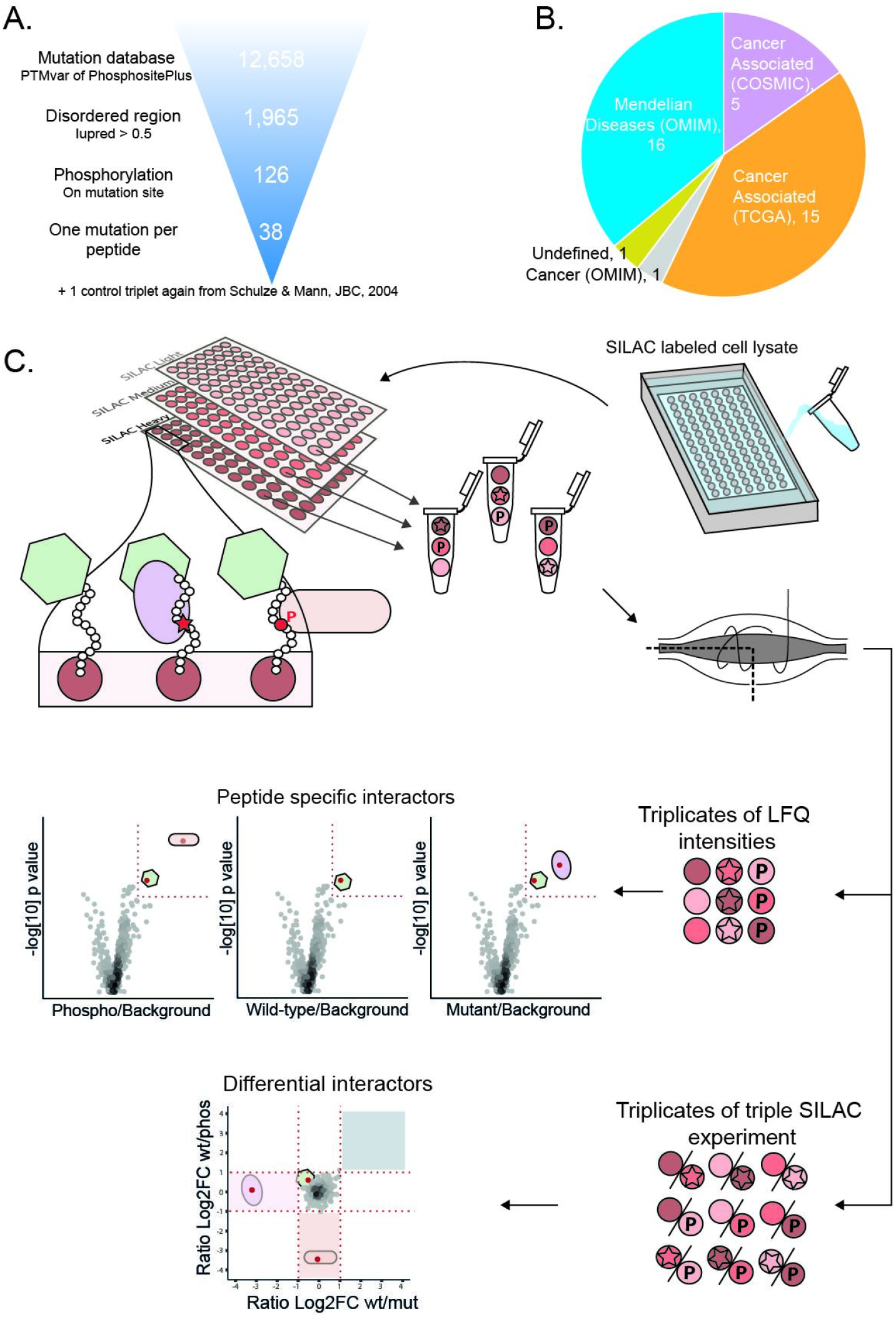
Candidate selection and experimental design. A) Selection scheme for peptide candidates from the PTMvar database of PhosphositePlus. B) Out of the 38 peptide candidates included in the screen, 20 are associated with cancer, 17 cause Mendelian diseases and for 1 candidate the disease is undefined. C) A scheme illustrating the experimental design and data analysis of the PRISMA screen. Three peptide states (empty: wild-type non-phosphorylated, P: wild-type phosphorylated, star: mutated)

To experimentally investigate how mutation and/or phosphorylation of these sites affects protein-protein interactions, we employed a peptide-based interaction screen (Meyer & Selbach, 2020). In particular, we adapted the “Protein interaction screen on peptide matrix” (PRISMA) set-up, where peptides are synthesized on a cellulose matrix that is used to pull-down interaction partners from protein extracts directly (Dittmar *et al*, 2019; Meyer *et al*, 2018; Ramberger *et al*, 2021b). To evaluate the impact of both the disease-causing mutation and phosphorylation, we designed an experiment that allowed us to compare interactions of all three peptide states (wild-type non-phosphorylated, wild-type phosphorylated, and mutated) directly with each other (Figure 1C). All 38 peptides were synthesized on cellulose membranes via SPOT synthesis (Frank, 1992) as 15-mers with the phosphorylation site in the central position.

We also included the three forms of a well-characterized EGFR-derived phosphopeptide as positive control (Schulze & Mann, 2004). Thus, the cellulose membranes contained 3 x 39 = 117 different peptide spots. For quantification, we employed stable isotope labeling with amino acids in cell culture (SILAC) (Mann, 2006). We used lysates of unlabelled (light or L), medium heavy (M), or heavy (H) SILAC-labeled HEK-293 cells. Each of the three differently SILAC-labeled cell lysates was incubated with a different copy of the cellulose membrane to pull-down specific interaction partners. After washing, peptide spots with their bound proteins were excised and combined with the two other peptide states from the other two membranes. We always combined the pull-downs of the wild-type non-phosphorylated, wild-type phosphorylated, and mutated peptides from three membranes into one sample. In this way, SILAC-based quantification allows us to directly evaluate differences in binding partners across the three peptide states.

We analyzed all 117 combined samples by high resolution shotgun proteomics. Two of the 38 mutations were mixed-up and were therefore excluded. In each of the remaining 111 samples, we identified between 300 and 1,000 proteins (Figure S1A), which results in a total number of about 70,000 putative protein-peptide interactions. The correlation of label-free quantification (LFQ) values between replicates was considerably higher than the correlation between different peptide pulldowns, demonstrating good reproducibility (Figure S1B). To further validate the data we grouped the ∼70,000 putative protein-peptide interactions into two categories: those that can be explained by a matching SLiM-domain pair and those that cannot. We observed higher protein LFQ values, i.e. enrichment, in pulldowns when the peptide exhibited a SLiM matching a protein domain, thus demonstrating that our data provides valuable insights into SLiM-dependent interactions (Figure S1C).

### Quantification enables detection of specific interactions

Quantification is an efficient means to differentiate specific interaction partners from non-specific contaminants (Richards *et al*, 2021; Meyer & Selbach, 2015; Vermeulen *et al*, 2008; Smits & Vermeulen, 2016). Following our previously published strategy (Meyer *et al*, 2018), we used two consecutive quantitative filters to identify proteins that exhibit both specific interactions with a particular peptide and are influenced by its phosphorylation and/or mutation state. First, we used label-free quantification (LFQ) to identify proteins that interact specifically with a peptide compared to all other peptides in the screen. For the EGFR control peptide, this LFQ filter identified 2, 2 and 12 specific interaction partners of the wildtype, muted and phosphorylated state, respectively (Figure 2A). Second, we employed SILAC-based quantification to compare interactions across these three peptide states. We present the SILAC ratios of wildtype versus mutant and wildtype versus phosphorylated states as a scatter plot (Figure 2B). As expected for this tyrosine-phosphorylated peptide, a number of SH2-domain containing proteins are LFQ-specific interactors of the phosphorylated form (Figure 2A, right panel), and the SILAC data confirms their phosphorylation-dependent interaction (Figure 2B). The autophosphorylated EGFR peptide has been shown to bind to the GRB2 protein (Batzer *et al*, 2023; Schulze & Mann, 2004; Lundby *et al*, 2019). In our analysis, as depicted in the scatter plot, it is evident that GRB2 exhibits the highest phos/wt and phos/mut ratio >20, in all three replicates. MS1 spectra for a GRB2-derived peptide across label swaps are shown as an example (Figure 2C). In addition to GRB2, we identified several other SH2 domain-containing proteins as binders, including STAT3 and PLCG1, consistent with previous data (Lundby *et al*, 2019).

**Figure 2.**
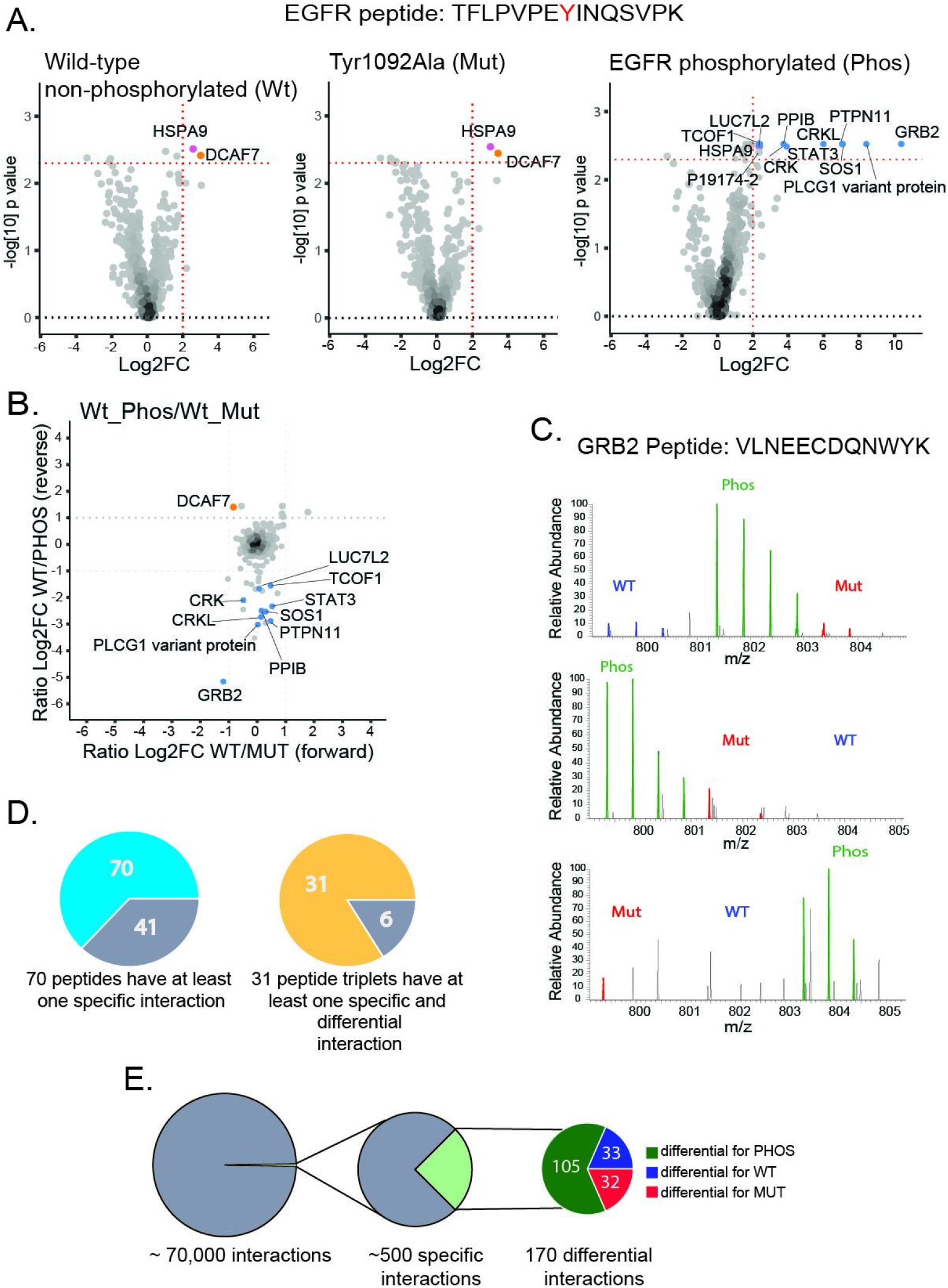
Quantification enables detection of specific interactions. A) Volcano plots (based on LFQ quantification) show that only the phosphorylated form of the EGFR control peptide specifically interacts with SH2 domain-containing proteins. Proteins passing the LFQ significance cut off are highlighted in different colors (blue: phosphopeptide specific, orange: wild-type and mutated peptide specific, purple: specific for all three peptide forms). B) SILAC ratios scatter plots of differential interactions (wt/mut versus wt/phos) show that several SH2 domain-containing proteins bind preferentially to the phosphorylated EGFR control peptide. C) Mass spectra of an exemplary GRB2 peptide show specific GRB2 binding to the phosphorylated EGFR peptide in all three replicates. D) 70 out of all 111 peptide pulldowns had at least one specific interactor (left). 31 out of 37 peptide triplets had at least one specific and differential binder (right). E) The two quantitative filters drastically reduce the number of interactions with the LFQ filtering revealing around 500 specific interactors and then SILAC filtering revealing 170 specific and also differential interactors.

Having validated our quantitative filters for the positive control, we applied this strategy to the entire dataset (Figure S3). LFQ-based filtering reduced the ∼70,000 interactions to approximately 500 (Figure 2E and S2A). 70 of the 111 peptides had at least one specific interaction partner and 31 out of 37 peptide triplets had at least one specific and differential binding partner (Figure 2D). Among all LFQ-specific interactors, SILAC-based filtering identified 170 interactions to be differential between the three peptide states. Thus, our filtering approach dramatically reduces the number of interactions. Among the specific and differential interactors, 105 preferentially interact with the phosphorylated peptide, 33 with the non-phosphorylated wild-type and 32 with the mutated peptide (Figure 2E and S2B).

### A network of interactions affected by phosphorylation and/or pathogenic single amino acid variants

We present all 170 specific and differential interactions with the 31 peptides in the form of an interaction network (Figure 3A). To illustrate which interactions can be explained by protein domains binding to a peptide SLiM, we extracted all annotated SLiMs from the peptide sequences and highlighted the proteins in the network that contain matching domains (Table S3). For example, the network contains three tyrosine phosphorylated peptides (including the EGFR control) that have SH2-domain binding SLiMs, and we identified six SH2 domain-containing proteins as specific and differential interactors (Figure 3B). Interactors other than those containing SH2 domains likely bind indirectly to these tyrosine phosphorylated peptides. For instance, the interaction between the EGFR peptide and SOS1 is likely mediated through GRB2 (Chardin *et al*, 1993). We also observed eleven peptides interacting with PIN1 (Peptidyl-prolyl cis/trans isomerase NIMA-interacting 1) (Figure 3C). All of these peptides contain an [pS/T]P motif known to interact with the WW domain of PIN1 (Lee *et al*, 2011). Finally, we were intrigued by the observation that MMTAG2, ARL6IP4 and PC4 interact with many phosphopeptides (9, 7 and 6, respectively). While these proteins do not contain annotated domains known to mediate phosphorylation-dependent binding, they carry regions with compositional bias for basic amino acids and have high predicted isoelectric points (Figure 3D). These proteins are therefore positively charged in the neutral pH range we used in the pulldown. Since phosphate groups are negatively charged, the observed phosphorylation-dependent interaction could reflect electrostatic effects. In summary, our interaction network contains a number of phosphorylation-dependent interactions that can be explained by SLiM-domain pairs or other specific protein features. Nevertheless, a majority of the identified interactions are novel, offering significant potential for elucidating the impact of SAVs and/or phosphorylation on protein function.

**Fig. 3:**
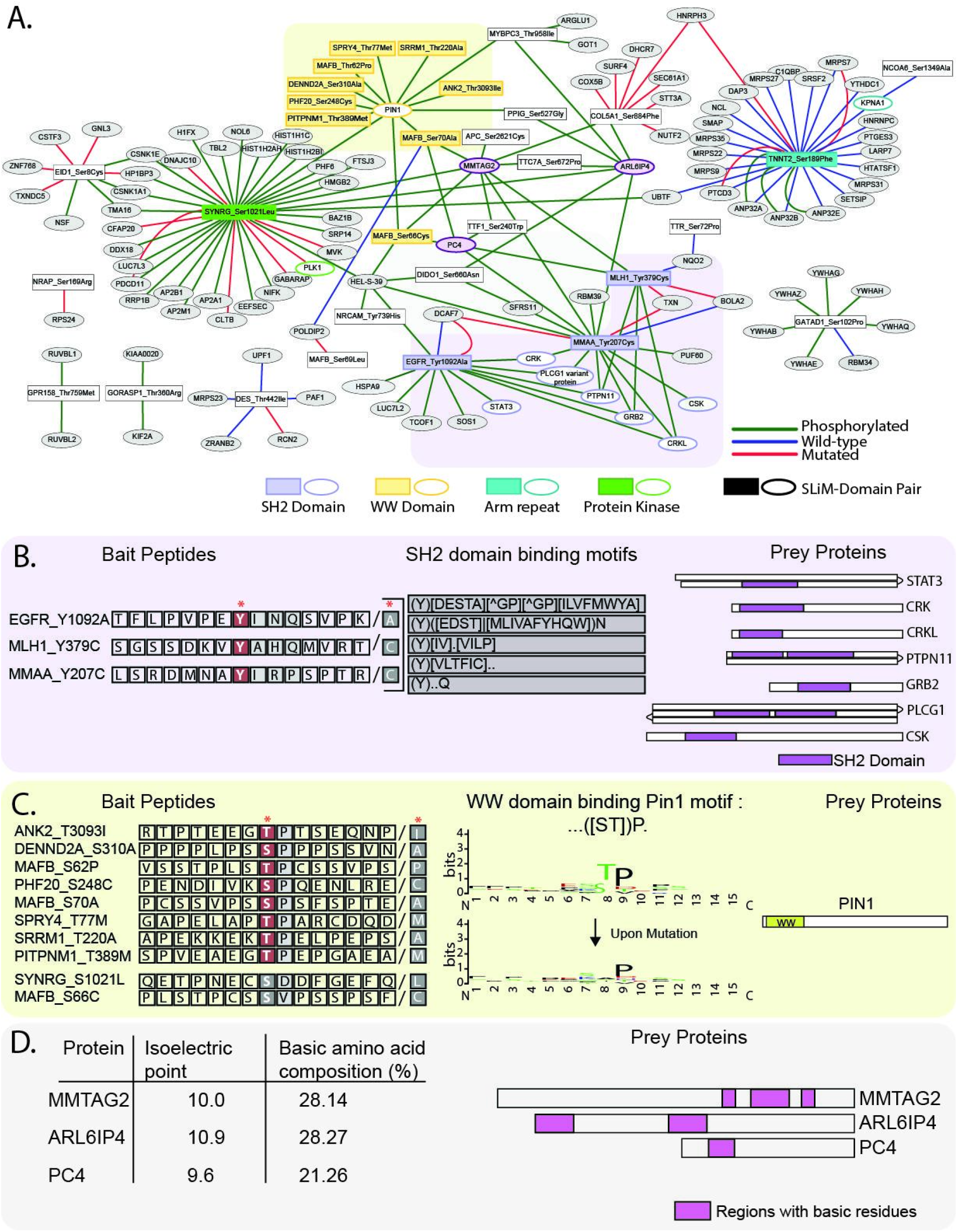
Network and SLiM analysis. A) A Peptide-Protein network that illustrates all interactions that passed both quantitative filters (LFQ and SILAC). Rectangles represent bait peptides and ovals interacting proteins. Edges indicate preferential binding. Highlighted regions and colored nodes indicate interactions that can be explained by annotated SLiM-Domain pairs. B) Interactions explainable by SH2 domains. Peptide sequences and corresponding SH2 domain binding SLiMs (left and middle) are displayed, along with a graphical representation of the SH2 domain-containing proteins (right). C) Peptides binding to PIN1 protein. 8/10 peptides contain the ([ST])P motif that binds to the WW domain of PIN1 in a phosphorylation-dependent manner. The sequence logo demonstrates the SLiM that is lost upon mutation. D) Peptides interacting with MMTAG2, ARL6IP4 and PC4. These proteins have a compositional bias towards positively charged amino acids (right), explaining preferential binding to the more negatively charged phosphorylated peptides.

### A phosphorylation site in the transcription factor GATAD1 interacts with 14-3-3 proteins

To further investigate the potential function of novel interactions, we directed our attention to the binding of multiple members of the 14-3-3 family of proteins to a GATAD1-derived phosphopeptide (Figure 4A and S3). 14-3-3 family proteins are important regulatory molecules involved in a staggering number of cellular processes that interact with target proteins in a phosphorylation-dependent manner (Pennington *et al*, 2018; Ballone *et al*, 2018; Fu *et al*, 2000). GATAD1 is a transcription factor affecting proliferation and cell cycle via controlling AKT signaling (Sun *et al*, 2018). The GATAD1 S102P mutation we investigated in the screen was described to cause dilated cardiomyopathy (DCM) in an autosomal recessive manner in a consanguineous family (Theis *et al*, 2011). Experiments in zebrafish provided additional evidence for the pathogenicity of this mutation (Yang *et al*, 2016). However, neither the pathogenic mechanism nor the function of this GATAD1 phosphorylation site are currently known. Intriguingly, although GATAD1 is expressed in many tissues, the only evidence for phosphorylation of this site comes from murine heart tissue, indicative of a heart-specific function (Lundby *et al*, 2013; Hornbeck *et al*, 2015).

**Figure 4:**
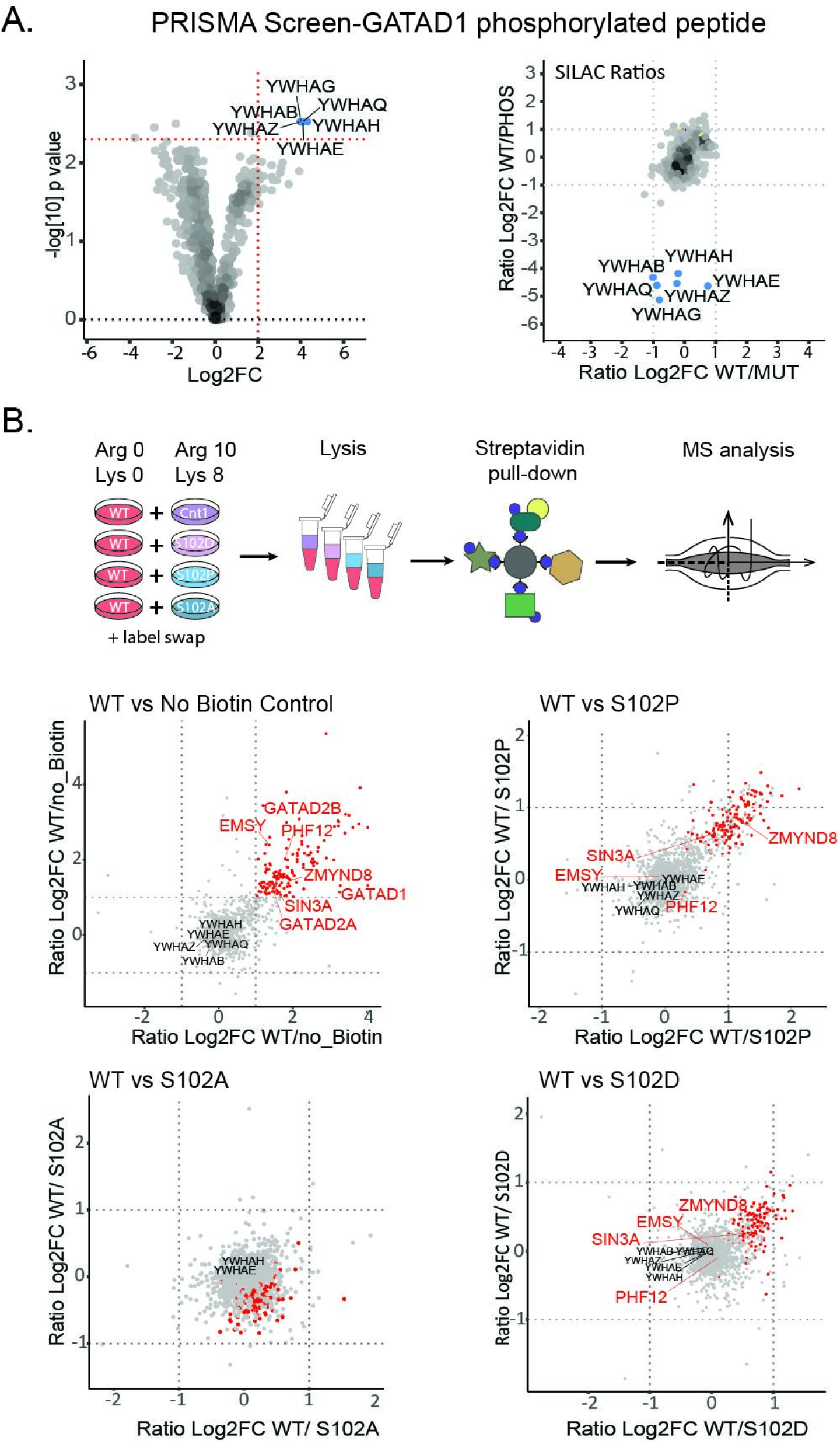
Interaction of 14-3-3 family proteins with GATAD1 phosphorylated on serine 102. A) The phosphorylated GATAD1 peptide specifically interacts with the multiple 14-3-3 protein family members, despite lacking an annotated 14-3-3 binding motif. Both the Volcano plot (left) and the SILAC scatter plot (right) show 14-3-3 proteins as specific and differential binders of the phosphorylated GATAD1 peptide (blue: LFQ specific binders to the phosphorylated peptide). B) BioID results for full-length GATAD1. Top: Experimental design. Scatter plots display protein SILAC ratios of two biological replicates (label swap) for different experiments. Top left: WT GATAD1 versus the “no biotin” control identifies several well-established GATAD1 interactors (red and labeled) and 130 additional proteins (red), indicating that the BioID data reflects GATAD1 biology. In the other three scatter plots WT GATAD1 is compared to different mutants. The S102P and S102D mutant show similar but overall somewhat reduced interactions while the S102A mutant is essentially identical to the wild-type. 14-3-3 family proteins were detected (YWHAB, YWHAE, YWHAH, YWHAQ and YWHAZ) but do not show any specific interaction with any of the GATAD1 proteins.

To further characterize the interactome of WT and mutated GATAD1 we first used BioID – a proximity-dependent biotinylation approach that systematically captures the “neighborhood” of proteins in living cells (Gingras *et al*, 2019; Roux *et al*, 2012; Rees *et al*, 2015). To this end, we generated stable inducible Flp-In^TM^-293 cells expressing GATAD1 fused to the promiscuous biotin ligase BirA*. Immunofluorescence confirmed that the BirA*-GATAD1 fusion protein localizes to the nucleus (Figure S4), as expected. Hence, the tag does not appear to affect GATAD1 localization. BioID identified 135 proteins as specific GATAD1 interactors (Figure 4B). This included several well-known GATAD1 binders such as the transcriptional regulators EMSY, PHF12 and SIN3A. Together, these proteins form the EMSY complex, which binds to H3K4me3-marked, active promoters (Vermeulen *et al*, 2010; Varier *et al*, 2016). Additionally, ZMYND8, which is known to form a complex with KDM5A and GATAD1 in response to DNA damage, was also identified as an interactor (Gong *et al*, 2017). More globally, gene ontology enrichment analysis of these 135 proteins revealed their involvement in transcriptional regulation, histone modification, and cell cycle (Zhang *et al*, 2019), consistent with the known biology of GATAD1.

We did not detect 14-3-3 family proteins as significant GATAD1 interactors in the BioID data. One possible explanation for this could be that GATAD1 is not phosphorylated on S102 in HEK cells. Indeed, even though GATAD1 is widely expressed and multiple phosphorylation sites (T34, T52, S55, S194, S235, Y248) have been identified in different cell lines (HeLa, KG1, K562, MKN-45), the only evidence for S102 phosphorylation comes from murine heart tissue (Lundby *et al*, 2013; Hornbeck *et al*, 2015). Therefore, to mimic phosphorylation, we created a stable cell line expressing BirA*-GATAD1 with a S102D mutation. This mutation introduces a negative charge and is thus considered to be “phosphomimetic” (Thorsness & Koshland, 1987; Pearlman *et al*, 2011). However, the BioID results for this mutant were overall very similar to the wildtype protein, and we did not observe increased interaction of 14-3-3 proteins (Figure 4B).

It should be noted that phosphomimetic mutations do not recapitulate all features of a phosphorylated residue (Dephoure *et al*, 2013; Pérez-Mejías *et al*, 2020). While phosphomimetic mutations have been successfully employed to imitate phosphorylation in many cases (Vieira-Vieira *et al*, 2022; Thorsness & Koshland, 1987; Koyano *et al*, 2014; Imami *et al*, 2018), a number of examples have shown that they are not universally an effective substitute for a phosphorylated amino acid (Somale *et al*, 2020; Kozeleková *et al*, 2022; Chino *et al*, 2022). The observed variations in behavior can be attributed to the contrasting biochemical properties of phosphorylated serine/threonine residues and aspartic/glutamic acid. In fact, the ability to work with phosphorylated peptides instead of phosphomimetic mutants is a key advantage of the peptide pulldown approach over genetic screens such as yeast two-hybrid and phage display (Kliche *et al*, 2023; Grossmann *et al*, 2015).

To unambiguously test whether or not GATAD1 phosphorylated on S102 binds to 14-3-3 family proteins we turned to isothermal titration calorimetry (ITC). To this end, we expressed human recombinant 14-3-3 epsilon (YWHAE) as a GST fusion protein and assessed its interaction with various peptides (Figure 5A). We found that GATAD1 pS102 interacted with 14-3-3ε with a binding constant of about 3 µM (Figure 5A). This interaction strictly depends on the phosphorylation of S102: Neither the non-phosphorylated wild-type peptide nor the peptide with the disease-causing S102P mutation showed detectable binding (Figure 5A). Importantly, the ITC data also showed that the phosphomimetic S102D and S102E mutant peptides do not interact with 14-3-3ε (Figure 5A). Hence, phosphomimetic mutations of this phosphorylation site do not recapitulate the phosphorylated state, explaining the BioID data.

**Fig. 5:**
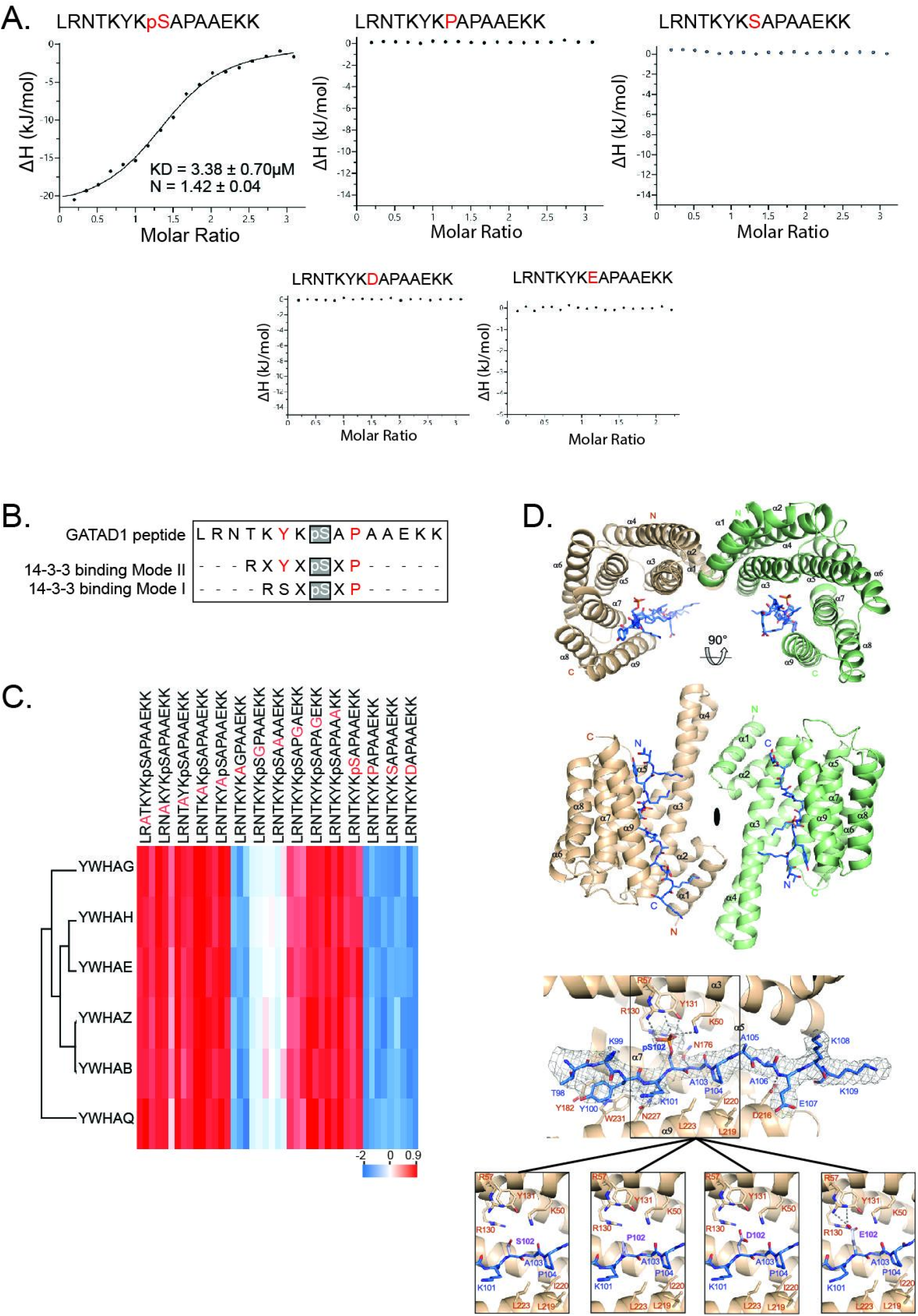
Structural analysis of the GATAD1 14-3-3 interaction. A) Isothermal titration calorimetry (ITC) validates the interaction of the GATAD1 phosphopeptide to 14-3-3 epsilon (YWHAE) with a K_d_ of 3.4 μM. In contrast, the wild-type non-phosphorylated, S102P and phosphomimetic (S102D and S102E) mutant peptides do not interact with 14-3-3 epsilon. B) The GATAD1 peptide displays similarities to 14-3-3 binding motifs but does not fully match. C) Heatmap of 14-3-3 proteins quantified in a PRISMA screen with GATAD1-derived peptides modified by alanine scanning. Results highlight the importance of proline at the +2 and glycine at the +1 position. D) Crystal structure of the 14-3-3 epsilon in complex with a phosphorylated GATAD1 peptide as determined by xray crystallography. Left: Side and top view of the 14-3-3 homodimer shown as cartoon structure with the two monomers colored in green and wheat. The bound peptides are shown in stick representation in blue color. The two-fold rotation axis between the monomers is indicated with an ellipsoid in the center of the homodimer. A detailed view into the peptide binding groove with hydrogen bonds as gray dashed lines and electron densities around each peptide as a gray mesh is shown. In bottom images the GATAD1 phosphoserine 102 was modeled as serine, proline, aspartic acid and glutamic acid, respectively, and highlighted in light blue.

### Structural analysis of the GATAD1 14-3-3 interaction

The discrepancy between the phosphorylated GATAD1 and the phosphomimetic mutants renders cell biological experiments involving these mutants inconsequential. Therefore, we focused on a more detailed structural analysis of the interaction with the phosphorylated peptide instead.

First, we sought to better characterize the GATAD1 14-3-3 binding motif. The Cantley lab initially determined two 14-3-3 recognition motifs (RSXpSXP and RXF/YXpSXP) using degenerate peptide libraries (Yaffe *et al*, 1997). Although neither of these motifs exactly matches the GATAD1 phosphorylation site, some sequence similarities are apparent (Fig. 5B). To experimentally determine which amino acids surrounding GATAD1-pS102 are important for 14-3-3 binding we used alanine scanning (Cunningham & Wells, 1989). To this end, we individually replaced each amino acid surrounding the phosphorylation site by alanine in a PRISMA screen (Figure 5C). Among the total number of 1,367 proteins detected in triplicate experiments, 121 were found to differ significantly between the 15 peptides, including six 14-3-3 protein family members (14-3-3 beta/alpha, gamma, epsilon, zeta, eta and theta). Comparing the abundance of the 14-3-3 proteins across peptide pulldowns yields a number of important insights (Figure 5C). First, all six 14-3-3 family proteins show essentially identical binding preferences. This is interesting since – despite the relatively high sequence homology – not all ligands show equal affinity for different 14-3-3 isoforms (Gardino *et al*, 2006). Second, all alanine-substituted phosphopeptides pulled down more 14-3-3 proteins than any of the non-phosphorylated peptides. Hence, the interaction strictly depends on phosphorylation, corroborating the ITC results. Third, we identified proline in position +2 and alanine in position +1 (substituted by glycine) as being important for binding while replacing none of the other amino acids had a significant effect. The relevance of proline at the +2 position is in good agreement with both 14-3-3 binding motifs. Also, while studies have shown that there is no strong preference for any specific amino acids in the +1 position, specific amino acids are not tolerated at this position, including glycine (Yaffe *et al*, 1997). This explains our observation that changing the alanine at position +1 to glycine disrupts binding.

To further investigate the different binding behavior of the tested GATAD1 peptides, we used X-ray crystallography to obtain the structure of 14-3-3ε in complex with the GATAD1 phosphopeptide at 3.1 Å resolution (see Supplemental Table S5 for the complete data statistics). The two 14-3-3ε proteins in the asymmetric unit (ASU) of the crystals formed a two-fold symmetric homodimer, which is typical for the 14-3-3 protein family. Each monomer consists of nine α-helices and harbors one phosphorylated GATAD1 peptide in a binding groove formed by helices α3, α5, α7 and α9 (Fig. 5D). Superimposing the 14-3-3ε structure with the 14-3-3ζ structure bound to phosphorylated polyomavirus middle-T antigen (mT) (Yaffe *et al*, 1997) revealed a high overall structural similarity with a room mean square deviation (RMSD) of 0.87 for 421 superimposed Cα atoms.

Residues 98-107 of the GATAD1 peptide were resolved in the electron density in both 14-3-3ε molecules in the ASU, while residues 108-109 were only visible in one of them (Fig. 5D). 14-3-3ε binds the central phospho-serine 102 of the GATAD1 peptide via the triad Arg57, Arg130, and Tyr131, which is highly conserved in the 14-3-3 family. In addition, Lys50 of 14-3-3ε directly interacts with the phosphate group of the peptide (Fig. 5D). GATAD1 Pro104 packs into a hydrophobic cage formed by 14-3-3ε Leu219, Ile220, and Leu223, whereas Tyr100 of the peptide shows a T-shaped π stacking against Tyr182 and Trp231 of 14-3-3ε. Additional hydrogen bonds between the main chain of the peptide and 14-3-3 residues Asn176, Asn227, and Asp216 (only in one monomer of the ASU) contribute to the peptide orientation (Fig. 5D, bottom).

Our structure explains the deficits of the non-phosphorylated GATAD1 peptide and the pathogenic S102P and the phosphomimetic S102D peptide variants to interact with 14-3-3ε. The non-phosphorylated serine, as well as the proline chain (in the S102P variant) and the aspartic acid side chains (in the S102D peptide variant) are too short to reach into the positively charged triad patch in 14-3-3ε for proper binding (Fig. 5D, bottom left). A glutamic acid side chain in the S102E mutant, on the other hand, appears to be able to reach into the binding triad; however, it is not able to form a hydrogen bond with Tyr131 nor Lys50 of 14-3-3ε (Fig. 5D, bottom right). In summary, our structural analyses reveal the sequence and structural requirements for GATAD1 pS102 14-3-3ε interaction, explain why binding strictly depends on phosphorylation and why it cannot be mimicked by phosphomimetic mutations.

### The 14-3-3 binding site in GATAD1 is a new nuclear localization signal

The absence of a cellular model mimicking GATAD1 phosphorylation complicates the analysis of the functional consequences of the interaction with 14-3-3 family proteins. To shed more light onto possible cellular mechanisms we took a closer look at the 121 proteins found to interact differentially to the 15 peptides in the alanine scanning experiment (Figure 6A). Intriguingly, we observed a cluster of 72 proteins that specifically interacted with all non-phosphorylated GATAD1 peptides. Enrichment analysis of KEGG pathways and Reactome Gene Sets identified nucleocytoplasmic transport as the most significantly enriched term (Figure 6A), with several carrier proteins facilitating nuclear localisation signal (NLS)-based nuclear import, including IPO4, IPO5, IPO7, TNPO1, TNPO3, and RANBP2. This intriguing observation suggests that this region of GATAD1 may function as a nuclear localization signal (NLS), potentially playing a role in nuclear transport. This is surprising, considering previous reports indicating that GATAD1 lacks an NLS and instead is believed to be imported via a piggyback mechanism alongside its binding partner HDAC1/2 (Knight, 2017; Vermeulen *et al*, 2010). If the 14-3-3 protein binding region of GATAD1 was indeed an NLS, recruitment of 14-3-3 to phosphorylated GATAD1 would be expected to block nuclear import. This is reminiscent of the function of 14-3-3 proteins in multiple prior studies that have demonstrated that 14-3-3 proteins can bind to phosphorylation sites in proximity to NLSs (Kumagai & Dunphy, 1999; McKinsey *et al*, 2001; Jung *et al*, 2020). This hinders recruitment of importins and thus blocks nuclear import.

**Figure 6.**
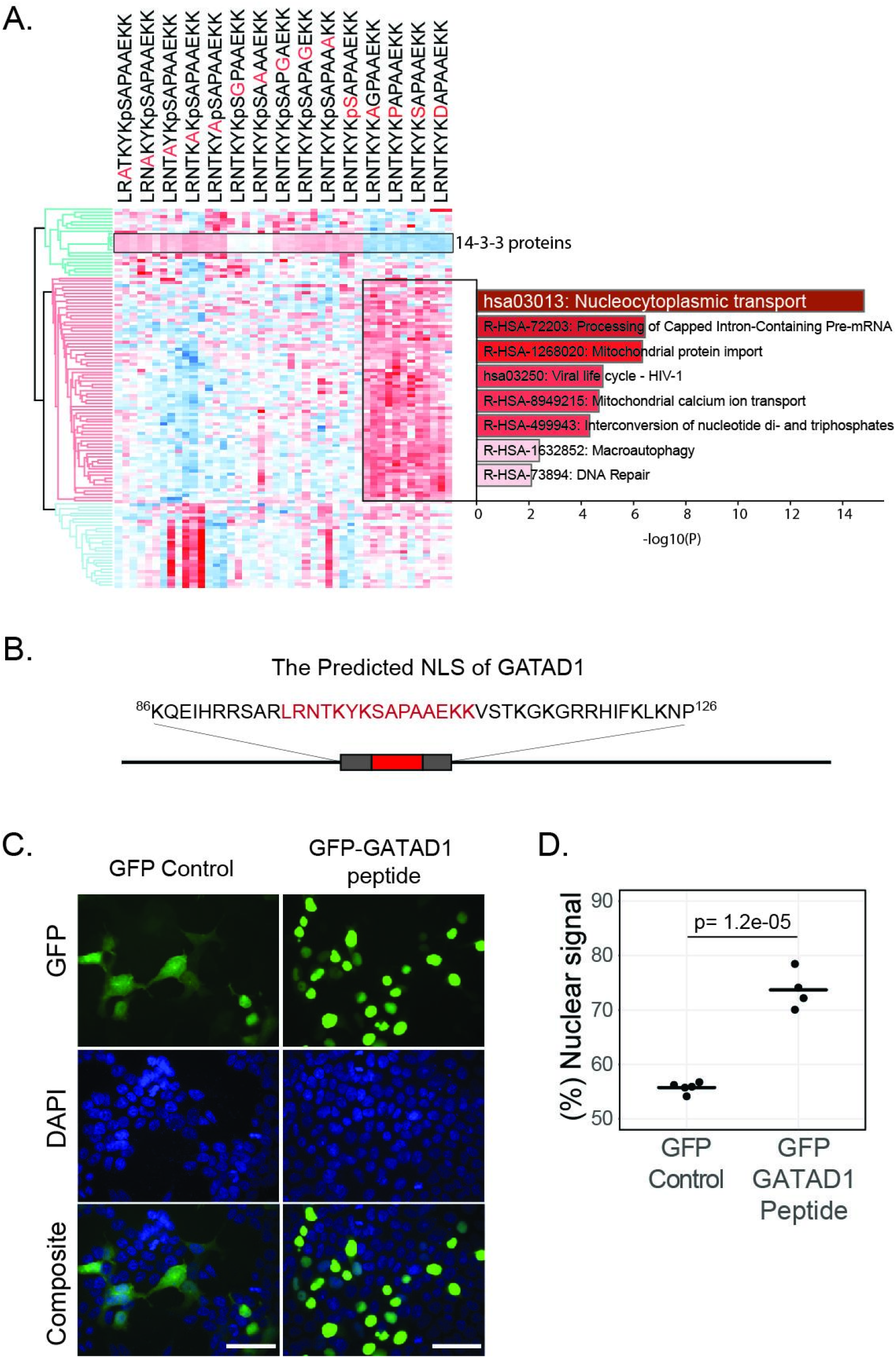
The 14-3-3 binding region of GATAD1 harbors a nuclear localisation signal (NLS). A) Heatmap of ANOVA significant proteins identified in the alanine scanning experiment. The largest protein cluster comprises 72 proteins that exhibit specific binding to the non-phosphorylated peptide forms. Enrichment analysis of KEGG pathway and Reactome Gene Sets of these proteins identified nucleocytoplasmic transport as the most significantly enriched term. Each peptide pull-down was performed in triplicates and all three replicates are represented in the heatmap. The cluster of 14-3-3 family proteins is also highlighted (see Fig. 5C for more details). B) The NLS of GATAD1 as predicted by NLStradamus. The GATAD1 peptide studied in this paper is highlighted in red. C and D) Immunofluorescence of GFP and GFP-fused to the putative GATAD1 NLS shows predominant nuclear localisation of the fusion protein. IF images were quantified using imageJ with at least with the minimum of 70 cells quantified for a condition. Scale bar = 50µm.

To further investigate this possibility, we first used the Hidden Markov Model-based tool NLStradamus to predict possible NLSs in GATAD1 (Nguyen Ba *et al*, 2009). Using the default settings (2 state HMM static, prediction cutoff 0.6), this algorithm indeed identified a region containing the 14-3-3 binding peptide as a potential NLS (Figure 6B). Next, to assess if this region is a functional NLS, we created a plasmid encoding a GFP-fusion protein and transiently transfected it into HEK-293 cells. As expected, the GFP-only control showed diffuse fluorescence throughout cells. In contrast, fusing GFP to the potential GATAD1 NLS resulted in predominantly nuclear localisation (Figure 6B and C). We conclude that this region is sufficient to drive nuclear localisation, indicating that binding of 14-3-3 proteins to this region could indeed regulate GATAD1 trafficking.

## Discussion

Understanding the functional consequences of mutations and posttranslational modifications in health and disease remains a major challenge, especially for intrinsically disordered regions. Here, we employed peptide-based interaction proteomics to investigate how known disease-causing SAVs of known serine, threonine or tyrosine phosphorylation sites affect protein-protein interactions. We identify many interactions affected by the mutation and/or the phosphorylation state (Figure 3). We further show that a phosphorylation site in the transcription factor GATAD1 that is mutated in a family of patients with dilated cardiomyopathy binds 14-3-3 proteins (Figure 4). Finally, we show that the 14-3-3 binding region of GATAD1 is a functional nuclear localisation signal, suggesting that binding of 14-3-3 proteins to phosphorylated GATAD1 could affect its subcellular localisation (Figure 6).

The 14-3-3 binding site of GATAD1 does not match a known 14-3-3 binding motif as defined in the ELM database (Kumar *et al*, 2020). We therefore extensively validated the interaction using isothermal titration calorimetry (ITC), alanine scanning and x-ray crystallography (Figure 5). The results obtained confirm the interaction and demonstrate that despite the lack of an exact match, the interaction is overall consistent with known 14-3-3 binding patterns. This highlights the previous observation that 14-3-3 family proteins can bind to a wide range of target sequences that sometimes deviate from the canonical motifs (Pennington *et al*, 2018; Ballone *et al*, 2018). More generally, our observation highlights the challenges associated with predicting SLiM-dependent interactions and the importance of experimental approaches (Davey *et al*, 2012; Tompa *et al*, 2014; Davey *et al*, 2023).

A key advantage of the PRISMA method used here is its ability to directly compare interaction partners of phosphorylated and non-phosphorylated wild-type and mutated peptides to each other. The ability to directly utilize phosphorylated peptides is a key advantage over phosphomimetic approaches since it avoids limitations associated with charged amino acids as surrogates for phosphorylation. In fact, we show that 14-3-3 binding to phosphorylated GATAD1 cannot be mimicked by amino acid substitution, providing another example for the problems associated with phosphomimetics (Somale *et al*, 2020; Kozeleková *et al*, 2022; Chino *et al*, 2022; Pérez-Mejías *et al*, 2020; Dephoure *et al*, 2013).

The inability of phosphomimetic mutations to resemble the phosphorylated state and the lack of GATAD1 phosphorylation in HEK-293 cells leads to a key limitation of this study: Due to the lack of a suitable model system, we cannot fully investigate the cellular consequences of the GATAD1 14-3-3 interaction. Moreover, we do not know which kinase(s) phosphorylate GATAD1 under which conditions. It is intriguing that phosphorylation of S102 has so far only been observed in the heart *in vivo* (Lundby *et al*, 2013). Together with the finding that the GATAD1_S102P mutation causes dilated cardiomyopathy (Theis *et al*, 2011), this is strongly indicative of a heart-specific function of this phosphorylation site. Additionally, our alanine scanning experiment showed that non-phosphorylated forms of the GATAD1 peptide interact with importin carrier proteins, suggesting the mutated region plays a role in nucleocytoplasmic transport of GATAD1. Indeed, we validated that this GATAD1 region harbors a functional NLS (Figure 6). These observations could help explain the perturbed subcellular distribution pattern of GATAD1 in cardiomyocytes of patients affected by this mutation (Theis *et al*, 2011). Importantly, however, the BioID (Figure 4) and alanine scanning data (Figure 6) indicate that the mutation itself does affect nuclear translocation. Building upon existing data for the role of 14-3-3 proteins in nucleocytoplasmic transport (Kumagai & Dunphy, 1999; McKinsey *et al*, 2001; Grozinger & Schreiber, 2000; Jung *et al*, 2020), we propose that the binding of 14-3-3 to GATAD1 masks the NLS and impairs the protein’s nuclear localization. Nevertheless, due to the lack of a cellular model system, we are not able to experimentally validate this hypothesis.

Overall, our study highlights the potential of the PRISMA screen for elucidating the functional consequences of mutations and PTMs, advancing our knowledge of protein regulation and interaction networks. In the future, it will be interesting to also investigate mutations adjacent to phosphorylation sites, not just changes of the phosphorylated residue itself. Such mutations can also modify SLiMs and thereby change PPIs. For example, the lung cancer associated P1019L mutation in EGFR has already been shown to switch the binding specificity of the adjacent phosphorylation site pY1016 (Lundby *et al*, 2019). Combining PRISMA with ultrahigh throughput proteomics enables the analysis of a much larger number of sites (Bekker-Jensen *et al*, 2020b; Messner *et al*, 2021; Tomioka *et al*).

## Material and Methods

### Cell Culture

HEK-293, HEK-293T, and Flp-In^TM^-293 T-REx cells were cultured in DMEM medium supplemented with 10% fetal calf serum (FCS) from Pan-Biotech. The cells were incubated at 37°C with 5% CO2.

For SILAC labeling, SILAC DMEM from Life Technologies was used. The SILAC DMEM was supplemented with 10% dialyzed FCS from Pan-Biotech, glutamine (Glutamax, Life Technologies), and non-essential amino acids. Different SILAC formulations were employed: Arg0 and Lys0 for “light” labeling, Arg6 and Lys4 or only Lys4 for “medium-heavy” labeling, or Arg10 and Lys8 (Sigma-Aldrich) for “heavy” labeling. To ensure complete incorporation of SILAC amino acids, cells were passaged and grown for at least 2 weeks or approximately 8 doublings.

### Peptide-Protein Interaction Screen

#### Selection of peptide candidates

Pathogenic mutations affecting phosphorylation sites were selected from the PTMVar dataset in PhosphositePlus (Hornbeck et al., 2015). The PTMVar dataset provides comprehensive information on Post-Translational Modifications (phosphorylation, ubiquitylation, acetylation, methylation, and succinylation) that overlap with genetic variants associated with diseases and genetic polymorphisms (PMID: 20668451).

For our analysis, we specifically focused on entries in which mutations were causative for disease (VAR_TYPE = Disease), resulted in changes to phosphorylation sites (MOD_TYPE = Phosphorylation), and directly altered the phosphorylated amino acid (VAR_POSITION = 0). We further filtered the entries to include only peptides originating from disordered regions and with no additional reported PTMs on the peptide (peptides with more than one lowercase letter in the MOD-SITE_SEQ column were excluded). Disorder was predicted using IUPred (Dosztányi *et al*, 2005), using ‘SHORT’ profile and considering a neighborhood of 25 amino acids. Regions with an IUPred score higher than 0.5 are considered disordered. In cases where the residue of interest mutated into multiple amino acids, we retained only one mutant form. Specifically, we excluded amino acid substitutions that could still be phosphorylated (Ser, Thr, Tyr). Following these criteria, we selected a total of 36 variant peptides for further analysis (see Supplemental Table S1). As a positive control we included an EGFR peptide known to contain an SH2 domain binding motif (Schulze & Mann, 2004).

#### Experimental Setup

A total of 117 peptides, consisting of 15 amino acids each, were synthesized in situ on a cellulose membrane using SPOT synthesis techniques (Frank, 1992) provided by JPT Peptide Technologies in Berlin, Germany. Among these peptides, 39 were wild-type non-phosphorylated, 39 were mutated, and 39 were phosphorylated. Whenever possible, the mutation site was positioned at position 7, which is in the center of the immobilized peptides, with their C-termini serving as the point of immobilization.

For the experimental procedure, three membranes were initially incubated with a lysis buffer containing HEPES (50 mM, pH 7.9), NaCl (150 mM), EGTA (1 mM), MgCl2 (1 mM), glycerol (20%), NP-40 (1%), SDS (0.1%), and sodium deoxycholate (0.5%). The membranes were incubated with the lysis buffer until completely moistened. To minimize non-specific binding, the membranes were then treated with yeast t-RNA at a concentration of 1 mg/ml for 10 minutes at 4°C, following two 5 minute washes with the lysis buffer. Subsequently, the membranes were incubated with HEK-293 cell lysate (5 ml at a concentration of 8 mg/ml) labeled as heavy, medium, or light, and this incubation step was carried out for 2 hours at 4°C. Afterward, the membranes were washed twice for 5 minutes with a washing buffer containing HEPES (50 mM, pH 7.9), NaCl (150 mM), EGTA (1 mM), and MgCl2 (1 mM). Finally, the membranes were air dried to complete the procedure.

#### Sample preparation for mass spectrometric analysis

Following the drying of the membranes, the peptide spots were carefully excised using a 2 mm diameter ear punch (Carl Roth). To facilitate further analysis, SILAC triplets were combined in a single 96-well plate, with each well containing 30 µl of denaturation buffer composed of 6 M urea (Sigma-Aldrich), 2 M thiourea (Sigma-Aldrich), and 10 mM HEPES at pH 8. The samples underwent reduction by adding 10 mM DTT (Sigma-Aldrich) and incubating for 30 minutes at room temperature (RT). Subsequently, an alkylation step was performed by adding 10 µl of 50 mM iodoacetamide (IAA) (Sigma-Aldrich) to each well and incubating for an additional 30 minutes at RT in the dark.

To initiate digestion, 0.5 µg of LysC enzyme was added to each well, and the samples were incubated for 1.5 hours at RT. Subsequently, the digestion was continued with 0.5 µg of trypsin (Promega) overnight at RT. The digestion process was halted the next day by acidifying the samples with 10 µl of 10% trifluoroacetic acid (TFA). Finally, the samples were desalted using the standard StageTip method (Rappsilber *et al*, 2003).

#### LC/MS method

The elution of samples from StageTips was carried out using buffer B, consisting of 0.5% acetic acid and 80% acetonitrile. Subsequently, the eluted samples were dried using a speedvac (Eppendorf) and resuspended in buffer A, which contained 0.5% acetic acid, prior to analysis. For nano-scale reversed-phase liquid chromatography coupled with tandem mass spectrometry (nanoLC/MS/MS), an Orbitrap Fusion mass spectrometer (Thermo Fisher Scientific) was utilized. The mass spectrometer was connected to a Thermo Ultimate 3000 RSLCnano pump equipped with a self-pulled analytical column measuring 150 mm in length and 100 μm in internal diameter. The analytical column was packed with ReproSil-Pur C18-AQ material (3 μm; Dr. Maisch GmbH). The mobile phases consisted of buffer A (0.5% acetic acid) and buffer B (0.5% acetic acid and 80% acetonitrile) (Battellino *et al*, 2023).

During the analysis, peptides were eluted from the analytical column at a flow rate of 500 nl/min, employing a gradient as follows: 5 to 10% B over 5 minutes, 10 to 40% B over 60 minutes, 40 to 99% B over 5 minutes, and maintaining 99% B for 5 minutes. The samples were measured in the orbitrap fusion mass spectrometer using the following methods: A full scan was performed with a resolution of 120,000 over an *m/z* range of 300-1500. The maximum injection time was set to 50 ms, and the AGC target was set at 400,000. Following the full scan, 20 MS/MS scans were conducted using isolation mode with a quadrupole, an isolation window of 1.6, HCD activation type, 30% collision energy, IonTrap detector type, and a maximum injection time of 35 ms.

#### Data Analysis

The analysis of the raw files was performed using MaxQuant version 1.6.2.6a (Cox & Mann, 2008), with default settings except for enabling the “match between run” and “requantify” options. Variable modifications were set to oxidation of methionines and acetylation of N-terminal residues, while fixed modifications were set to Carbamidomethylation. *In silico* digestion of proteins in the reference database, uniprot_human_20181012_canonical_isoform.fasta, and peptide identification were carried out using trypsin/P, allowing a maximum of 2 missed cleavages. The data were analyzed in both SILAC mode, with Lys4 as the medium-heavy label and Lys8, Arg10 as the heavy labels, as well as in label-free quantification (LFQ) mode. A false discovery rate of 1% was set at both the peptide-spectrum matches (PSM) and protein levels, and the assessment was performed by searching a decoy database generated by reversing the reference database.

The resulting MaxQuant files were further analyzed using R (R version 4.2.1 and Rstudio version 2022.07.1), including all statistical analysis and generation of figures. Figures were modified using Adobe Illustrator CS6. Initially, the ProteinGroups table was filtered to exclude proteins identified only by site, proteins from the reverse database, and potential contaminants. Peptide candidates identified based on their tryptic sites were also filtered out. LFQ values of the remaining proteins were log2 transformed and filtered to retain only those with at least one valid value within the three replicates. Missing values were imputed by randomly drawing values from a log distribution calculated as 0.25 times the standard deviation of the measured log-transformed values, down-shifted by 1.8 standard deviations. These LFQ values were used to determine peptide-specific interactors. The nonparametric Wilcoxon test was employed to compare the median LFQ values of the proteins identified in triplicates of a peptide with the median LFQ values of the background proteins. The background comprised proteins identified in all other pull-downs, excluding the corresponding variant peptide. Due to the charge effect, plenty of phosphorylated peptide variants interacted with the same proteins. We saw it fit to keep these proteins in the analysis. Therefore, all phospho-pull-downs were excluded from the background. The resulting p-values and fold changes were plotted as volcano plots. The significance cut-off was established using the control EGFR_pTyr1092 peptide, previously characterized as SH2 domain binding. A cut-off was set to capture all SH2 domain proteins identified with the phosphorylated peptide variant, requiring a log2 fold change greater than 2 and a p-value smaller than 0.005.

The SILAC ratios were then utilized to determine the differential interactors. The SILAC ratios were logarithmized, normalized by subtracting the median SILAC ratio of each experiment from all individual SILAC ratios in that experiment, and subjected to label swap. This resulted in triplicates of triple SILAC experiments (wt/mut, wt/phos, and phos/mut ratios). The SILAC data were filtered to retain only those with at least two valid values out of the three replicates, and the medians of these ratios were plotted on a scatter plot with wt/mut on the x-axis and wt/phos on the y-axis. Differential proteins were identified as those with wt/mut, phos/mut, and/or wt/phos ratio log2 fold change greater than one or smaller than minus one. The specific and differential proteins obtained are summarized in table S2 and were used to create the peptide-protein network using Cytoscape v.3.9.1 (Shannon *et al*, 2003).

### Proximity labelling (BioID) of GATAD1 variants

#### Generation of stable HEK-293 cell lines expressing mutant GATAD1 variants

The GATAD1 gene variants were ordered in a pTwist ENTR Kozak vector from Twist Bioscience. The genes were then transferred to a pDEST_pcDNA5_BirA_FLAG_Nterm vector (Couzens *et al*, 2013) using the Gateway Clonase II system (Thermo Fisher Scientific). The resulting vectors, pEXPR_GATAD1-WT_N-term_BirA_FLAG, pEXPR_GATAD1-S102P_N-term_BirA_FLAG, pEXPR_GATAD1-S102D_N-term_BirA_FLAG, and pEXPR_GATAD1-S102A_N-term_BirA_FLAG, were used to generate tetracycline-inducible stable Flp-In^TM^-293 T-REx cells (Invitrogen). A 6-well plate was co-transfected with 0.5 µg of the pEXPR plasmid and 1 µg of the pOG44 Flp-recombinase expression vector (Thermo Fisher Scientific) using 4.5 µg of PEI (Polysciences) as a transfection reagent. The following day, the cells were trypsinized and transferred to a 10 cm dish containing DMEM supplemented with 10% FBS and 200 µg/ml hygromycin B (Invivogen) for selection. The cells were cultured for approximately 18 days with media exchange every 3 days until visible colonies were observed. After around 18 days, the colonies were trypsinized and transferred to another 10 cm dish for further characterization and experiments.

#### Interactome analysis of GATAD1 variants using BioID

Stable Flp-In^TM^-293 cell lines expressing GATAD1 variants were cultured using SILAC light (Lys0, Arg0) and SILAC heavy (Lys8, Arg10) media, as explained previously. After full labeling, cells were split into 15 cm dishes and were treated with 1 μg/mL tetracycline for 24 hours to induce the expression of GATAD1. After the induction period, cells were incubated with 50 mM biotin overnight for proximity biotinylation. As a control, wild-type GATAD1 cells, without biotin addition, were used. Samples were multiplexed, as shown in Figure 4B with both forward and reverse label swap. Cells were then lysed in lysis buffer (50 mM Tris-HCl pH 7.5, 150 mM NaCl, 1% Triton X-100, 1 mM EDTA, 1 mM EGTA, 0.1% SDS) with freshly added protease inhibitors and 1% sodium deoxycholate. To digest the excess DNA, 1 μl of Benzonase was added, and the samples were incubated for 20 minutes at 37 °C. Biotinylated proteins were enriched for 3 hours at 4°C using streptavidin-sepharose beads (GE Cat# 17-5113-01). Beads were pre-washed twice with 0.01% BSA and twice with lysis buffer. After enrichment, the beads were washed one time with lysis buffer, two times with washing buffer (50 mM HEPES-KOH pH 8.0, 100 mM KCl, 10% glycerol, 2 mM EDTA, 0.1% NP-40), and six times with 50 mM ammonium bicarbonate. Beads were resuspended in ammonium bicarbonate, and an on-bead protein digest with 1 μg of trypsin followed overnight. The next day, the digested proteins were transferred to a fresh tube and were incubated with 10 mM DTT for 30 minutes at 37 °C and with 55 mM iodoacetamide for 20 minutes at 37 °C in the dark. Samples were desalted with StageTips.

Peptides were separated using reversed-phase liquid chromatography (EASY nLC II 1200, Thermo Fisher Scientific) with self-made C18 microcolumns (20 cm long) packed with ReproSil-Pur C18-AQ 1.9 μm resin (Dr. Maisch, cat# r119.aq.0001). The chromatography system was coupled online to the electrospray ion source (Proxeon) of an Orbitrap Exploris 480 mass spectrometer (Thermo Fisher Scientific). The mobile phase consisted of buffer A (0.1% formic acid and 5% acetonitrile) and buffer B (0.1% formic acid and 80% acetonitrile). Peptides were eluted using a gradient with increasing concentrations of buffer B over 110 minutes, at a flow rate of 250 nl/min. Mass spectrometry data was acquired in data-dependent mode with settings for one full scan (resolution: 60000; *m/z* range: 350-1600; normalized AGC target: 300%; maximum injection time: 10 ms), followed by top 20 MS/MS scans using higher-energy collisional dissociation (resolution: 15000; *m/z* range: 200-2000; normalized AGC target: 100%; maximum injection time: 120 ms; isolation width: 1.3 *m/z*; normalized collision energy: 28%).

Raw files were analyzed with MaxQuant, version 2.0.3.0. The Protein Groups data table was further processed and analyzed using R. The table was first filtered for contaminants, identified only by site and identified by the reverse database. The SILAC ratios were log2 transformed and the labels were swapped. The data was visualized in scatter plots with the forward SILAC ratios on the x-axis and reverse SILAC ratios on the y-axis. Differential proteins were identified as those with ratio log2 fold change greater than one or smaller than minus one.

### Recombinant protein expression and purification

The 14-3-3ε (YWHAE) cDNA was purchased from Twist Bioscience in the pTwist Chlor High Copy vector and cloned into the pGEX6P1 plasmid (GE Healthcare) for recombinant expression as a GST-fusion protein followed by a Prescission protease cleavage site. The plasmid was freshly transformed into *Escherichia coli*C41 (DE3) cells. Cultures were grown in terrific broth supplemented with ampicillin (100 μg/ml) at 37 °C and 80 rpm until an optical density at 600 nm of 0.7 was reached. Protein expression was subsequently induced by the addition of 300 μM isopropyl β-d-1-thiogalactopyranoside (IPTG), and the cultures grown for another 18 hours at 20 °C. Cells were harvested by centrifugation at 4,000 g and frozen at −20°C.

Following resuspension in lysis buffer (50 mM HEPES/NaOH pH 7.5, 500 mM NaCl and 3 mM Dithiothreitol (DTT)) supplemented with 1 mg deoxyribonuclease I (DNaseI, Roche) and protease inhibitor 4-(2-Aminoethyl) benzenesulfonyl fluoride hydrochloride (AEBSF), cells were disrupted using a microfluidizer (Microfluidics). The cell extract was centrifuged at 55,000 g for 45 min at 4 °C to remove insoluble parts. The cleared supernatant was applied onto a prepacked Glutathione Sepharose 4B column (Cytiva) equilibrated in lysis buffer. The column was extensively washed with lysis buffer. To cleave off the GST tag, 1 mg PreScission protease was diluted in 5 ml OCC buffer (50 mM HEPES/NaOH pH 7.5, 150 mM NaCl, and 3 mM DTT), applied to the column material, and incubated overnight at 4 °C. Cleaved 14-3-3ε protein was eluted from the column using OCC buffer, concentrated and loaded onto an 16/60 S200 column (Cytiva) equilibrated in SEC buffer (20 mM HEPES/NaOH pH 7.5, 150 mM NaCl and 2 mM DTT). Pure and homogenous 14-3-3ε was concentrated to 13 mg/ml, flash-frozen in liquid nitrogen, and stored at −80°C until further use.

### Isothermal titration calorimetry (ITC)

GATAD1 peptides and the positive control used in these experiments were obtained from JPT peptide technologies with an N-terminus in amine and a C-terminus in amide form with 95% purity. ITC experiments were performed using a PEAQ-ITC microcalorimeter (Malvern). All titrations were performed at 18 °C with 400 µM peptide in the syringe and 25 µM 14-3-3ε in the reaction chamber. The protein and the titration components were dissolved in a buffer containing 20 mM HEPES pH 7.0, 150 mM NaCl, and 2 mM DTT. Malvern software was used for data visualization and fitting.

### Crystallization and structure determination

The protein 14-3-3ε at a concentration of 13 mg/ml in 20 mM HEPES pH 7.5. 150 mM NaCl, 2 mM DTT was combined with the GATAD1 peptide 95-LRNTKYKpSAPAAEKK-109 in 1:2 molar ratio for complex formation. Crystallization setups were performed with the sitting-drop vapor diffusion method by using a Gryphon pipetting robot (Matrix Technologies Co.) for pipetting 200 nl of protein to an equal volume of precipitant solution. The Rock Imager 1000 storage system (Formulatrix) was used for storing and imaging of the experiments. Crystals appeared within 3-10 days in 19% PEG 3350, 0.35 M NaBr, 0.1 M BisTris-Propane pH 6.5 at 20 °C and were flash-frozen in liquid nitrogen in the presence of 20% ethylene glycol. Diffraction data were collected on BL14.1 at the BESSY II electron storage ring operated by the Helmholtz-Zentrum Berlin (Mueller *et al*, 2015), processed and scaled using XDSapp (Krug *et al*, 2012). The 3.2 Å structure was solved by molecular replacement with Phaser (McCoy *et al*, 2007) using the 14-3-3 protein epsilon structure (PDB: 2BR9) as a search model. The structure was built using COOT (Emsley *et al*, 2010) and iteratively refined with Refmac (Murshudov *et al*, 1997). Data statistics are summarized in Table S5. Crystals of 14-3-3 epsilon with the GATAD1 phospho-peptide belong to space group R32 containing 2 complex molecules per asymmetric unit connected by a two-fold rotational symmetry. Residues 2-235 of 14-3-3 are explained in the electron density, whereas residues 1 and 236-255 are disordered and therefore not visible in the electron density. The GATAD1 peptide is covered by residues 98-109 in chain C and residues 98-107 in chain P. 97.0% of the residues in the complex structure were in the favored regions and no outlier was observed in the Ramachandran map. The Ramachandran statistics was analysed using Molprobity (Chen *et al*, 2010). Figures were generated with PyMol (http://www.pymol.org). The atomic coordinates of 14-3-3ε with the GATAD1 95-LRNTKYKpSAPAAEKK-109 peptide have been deposited in the Protein Data Bank (PDB ID code XXX).

### Alanine Scanning

To investigate the contribution of individual amino acids in the GATAD1 peptide for binding to 14-3-3 proteins, an alanine scanning experiment was designed. In this experiment, each amino acid from the -5 to +5 positions surrounding the phosphorylated residue was mutated to alanine, except for original alanine residues which were mutated to glycine. The experiment also included the wild-type phosphorylated, wild-type non-phosphorylated, disease-causing mutant, aspartic acid mutant peptides, and a positive control. These 16 peptides were synthesized on a cellulose membrane in triplicates to facilitate the pull-down of proteins from HEK-293T cell lysate. The synthesis of peptides, pull-downs, and sample preparation for mass spectrometry followed the procedures described earlier for PRISMA screen. Subsequently, samples were eluted from StageTips using a solution containing 50% acetonitrile and 0.1% formic acid, dried, and resuspended in a solution containing 3% acetonitrile and 0.1% formic acid. LC-MS measurements were performed similarly as in the BioID experiment except for the gradient, which this time was 45 minutes, and the maximum injection time for MS2 scans was 22 ms.

The acquired mass spectra were subjected to further analysis using MaxQuant (version 1.6.3.4). Subsequent analysis was performed using Perseus (version 1.6.7.0). The ProteinGroups data table was filtered to exclude potential contaminants, proteins identified only by site, and proteins identified by the reverse database. LFQ values of the remaining proteins were log2 transformed, and only proteins with at least two valid values in three replicates were considered for analysis. An ANOVA significance test with an FDR of 0.05 was conducted. The LFQ values of significant proteins were z-scored by row and hierarchical clustering was performed (Figure 6A). To better visualize the 14-3-3 proteins, their LFQ values were extracted from the whole table and were z scored and clustered individually (Figure 5C).

### Immunofluorescence studies on Flp-In^TM^-293 T-REx stable cell lines expressing GATAD1 variant

Flp-In^TM^-293 cells stably expressing WT-GATAD1 were cultured and maintained. Cells were seeded on glass coverslips at a density of 50,000 cells per 24-well plate. The following day, tetracycline was added to a final concentration of 1 μg/mL, and the cells were incubated for an additional 24 hours. After the incubation period, the cells were fixed with 4% PFA for 15 minutes. Subsequently, the cells were washed three times for 5 minutes each with PBS. Permeabilization was achieved by incubating the cells with 0.5% Triton X-100 in PBS for 10 minutes, followed by two washes with PBST (PBS with 0.1% Triton X-100). To reduce background signal, the cells were incubated with blocking solution (1.5% BSA in PBST) for 1 hour. Next, the cells were incubated with the primary antibody, mouse monoclonal IgG anti-GATAD1 (sc-81092, 1:100, Santa Cruz), in blocking solution for 1 hour. After washing the cells three times for 5 minutes with PBST, the cells were incubated with the secondary antibody, goat anti-mouse IgG (H+L) Alexa Fluor 488 (1:500, Invitrogen), and phalloidin - Alexa 594 (1:500, Invitrogen) for 1 hour. Following another round of washing with PBST, the cells were incubated for 3 minutes with 0.1 μg/mL DAPI (Sigma Aldrich) in PBS. A final wash with PBS was performed, and the coverslips were then washed in MilliQ water and mounted on slides using ProLong Gold Antifade Mountant (Life Technologies).

### Immunofluorescence studies on HEK-293T cells transiently expressing GFP-GATAD1 Peptides

The pTwist_CMV_GFP_SPACER_KQSKQEIHRRSARLRNTKYKSAPAAEKKVSTKGKGRR constructs was ordered in the pTwist_CMV expression vector from TwistBioscience. HEK-293T cells were seeded on glass coverslips at a density of 0.1 x 10^6 cells per 12-well plate. The following day, cells were transfected with 1 µg of the pTwist_CMV plasmid using 3 µg of PEI (Polysciences) as a transfection reagent or with with 1 µg of the pDEST_GFP plasmid (Control). After 24 hours, the cells were fixed with 4% PFA for 15 minutes. Following fixation, cells were washed three times for 5 minutes each with PBS. Permeabilization was performed by incubating the cells with 0.5% Triton X-100 in PBS for 10 minutes, followed by two washes with PBST (PBS with 0.1% Triton X-100). Subsequently, the cells were incubated with 0.1 μg/mL DAPI (Sigma Aldrich) in PBS for 3 minutes. After another wash with PBS, the coverslips were rinsed in MilliQ water and mounted on slides using ProLong Gold Antifade Mountant (Life Technologies).

All images were acquired using a Leica DM5000B microscope with an HC PL APD 63x/1.4-0.6 objective. The acquired images were processed using Fiji ImageJ software (Schindelin *et al*, 2012).

For manual quantification, at least 70 cells were counted in each group to determine the subcellular localization of GFP-GATAD1 peptides (nuclear, cytosolic, or widespread). The DAPI channel was utilized to create a nuclei mask, which was used to separate the images into nuclear and cytoplasmic segments. The intensities of these segments were measured, and the mean intensities were utilized for further calculations. Statistical analysis was performed using a two-sample t-test, and the corresponding p-values are indicated in the plot.

## Acknowledgement

We would like to thank Martha Hergeselle and Christian Sommer for technical support, and Tobias Bock-Bierbaum and Carola Bernert for the expression and purification of the 14-3-3 protein. We further thank the advanced light microscopy facility at the MDC, especially Anca Margineanu for her help with microscopy and image analysis. T.R. was partially funded by Berlin School of Integrative Oncology (BSIO) and K.M. conducted the experiments as a JSPS International Research Fellows at Kyoto University, Japan as part of JSPS Summer Program (ID Number: SP1831).

## Author contributions

T.RR., K.M., and M.S. contributed to the design of the experiments and implementation of the project. T.RR., K.M., and Y.R. conducted experiments. T.RR., Y.R., and B.U. processed and analyzed the experimental data. A.A., K.I., Y.I., O.D., and M.S. supervised the work and provided resources. T.RR. visualized the data and produced most of the figures. T.RR. and M.S., wrote the manuscript with contributions from all co-authors.

