## Supplementary Figures for "Pathogenic mutations of human phosphorylation sites affect protein-protein interactions"

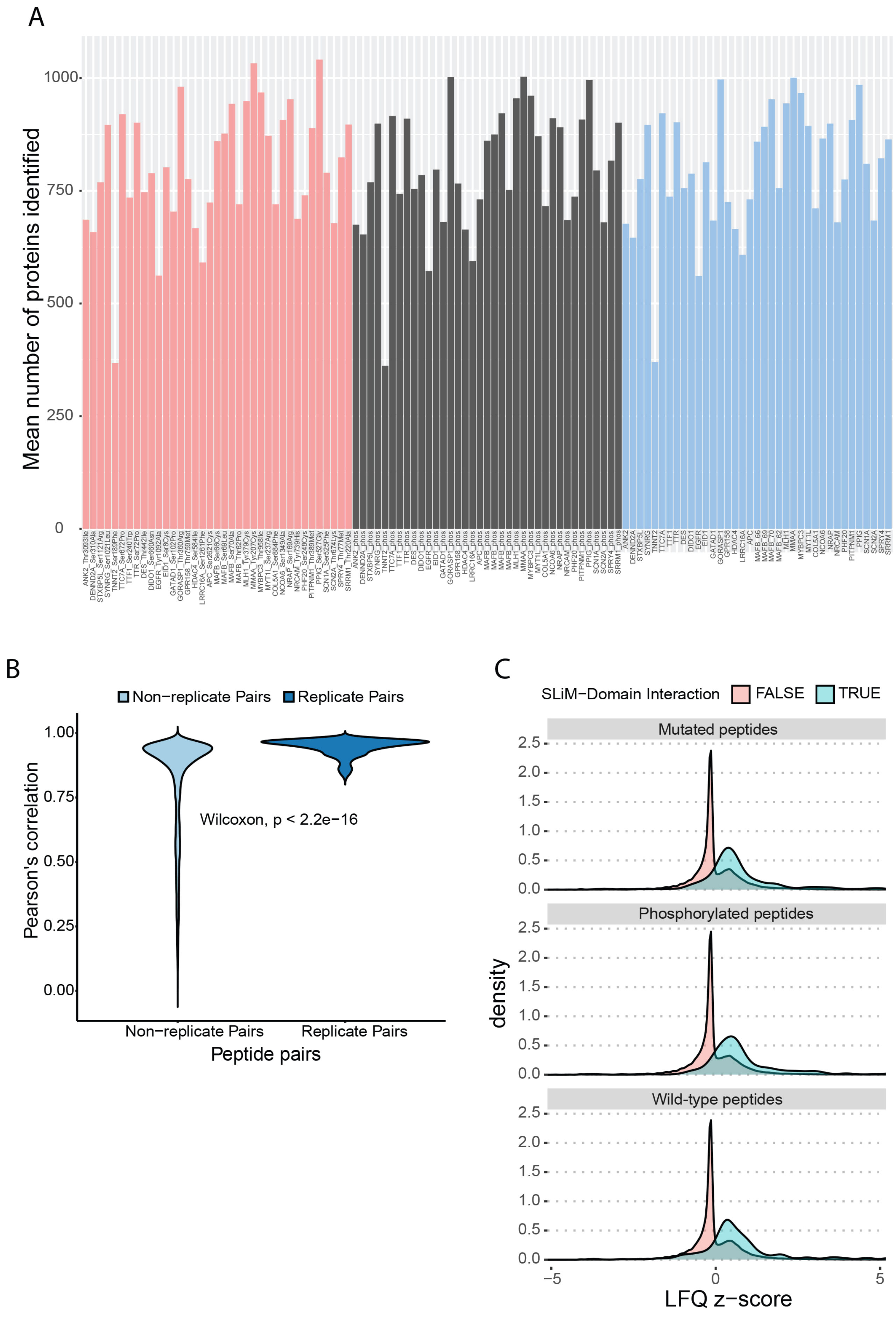


**Figure S1.** A) Bar plot showing the total number of proteins identified in each pull-down. B) Replicate pairs show a high correlation that changes between 0.75 to 1 while non replicate pairs correlate less and with a higher variability. C) Analysis of the identified interactors, after categorizing them based on their association with Short Linear Motif (SLiM)-Domain pairs, reveals that interactions explained by SLiM-Domain pairs show higher LFQ values. LFQ values were standardized (z-scored) for each protein across all peptides

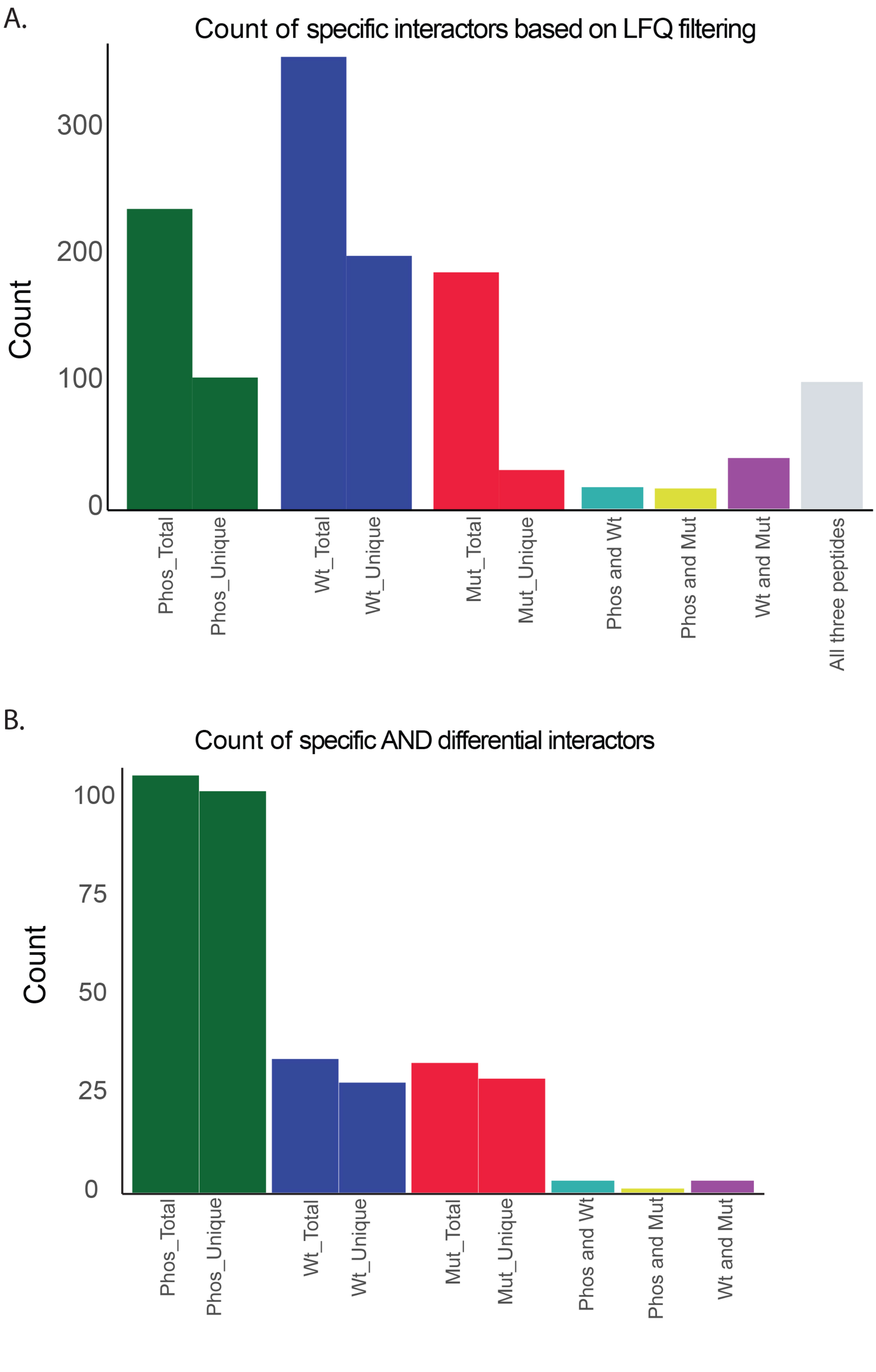


**Figure S2**. A) Bar plot illustrating the total number of proteins identified as significant based on LFQ filtering (Fold change >2 and p values < 0.005) for each pull-down. The y-axis represents the number of proteins, while the x-axis displays the type of interaction. The green bar indicates the number of specific interactors for phosphorylated peptides, with Phos_total including proteins specific to phosphorylated and other peptide forms, and Phos_unique representing proteins exclusively interacting only with the phosphorylated peptides. The same applies to Wt (wild-type) and Mut (mutated) peptides. B) Similar to the bar charts in A, but after implementing the SILAC filtering. These proteins have passed the LFQ significance cut-off and also exhibit a significant SILAC ratio (log2FC >1 or < -1).

### ANK2: RTPTEEGpTPTSEQNP

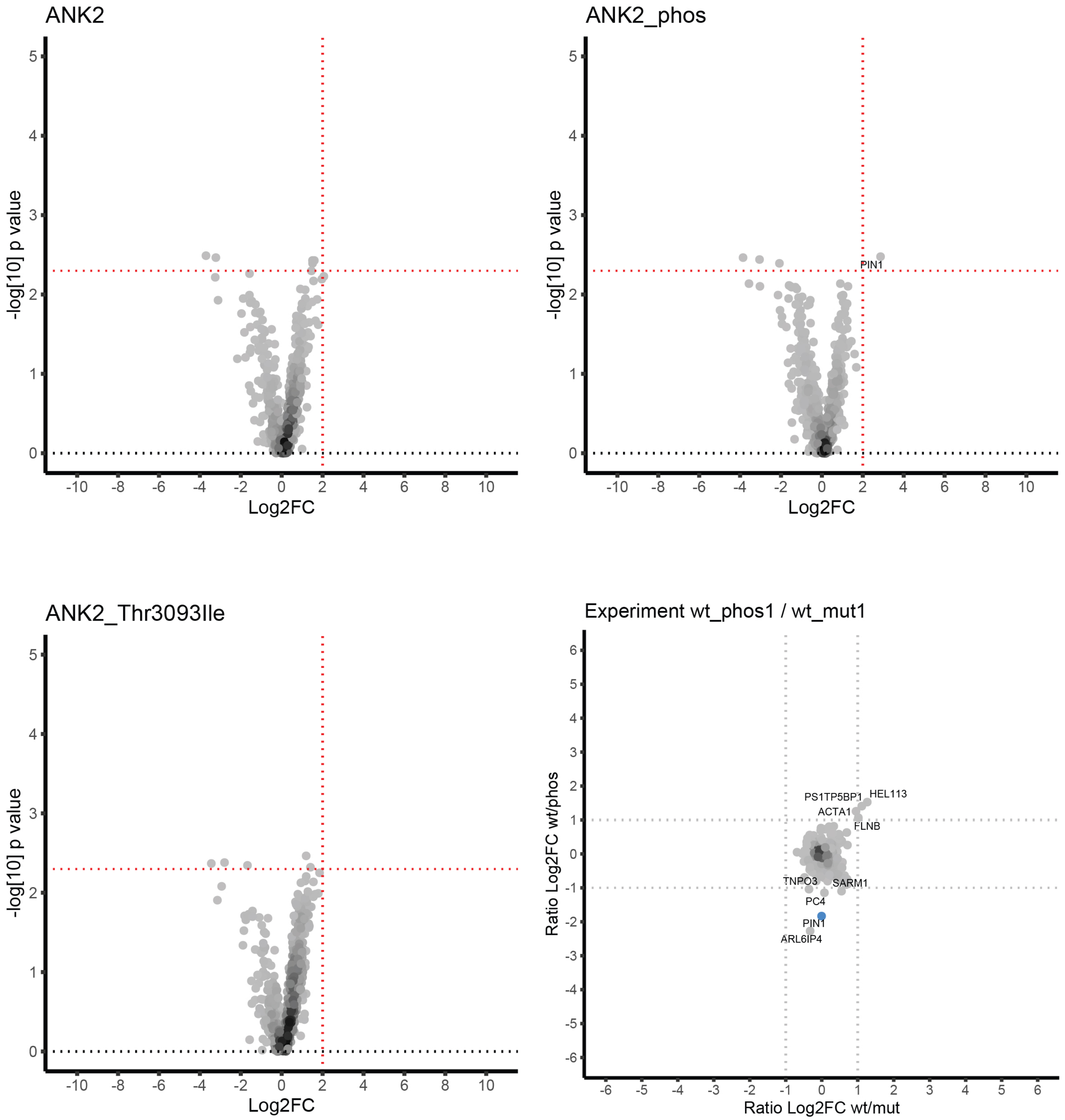

### APC: KIKENEFpSPTNSTSQ

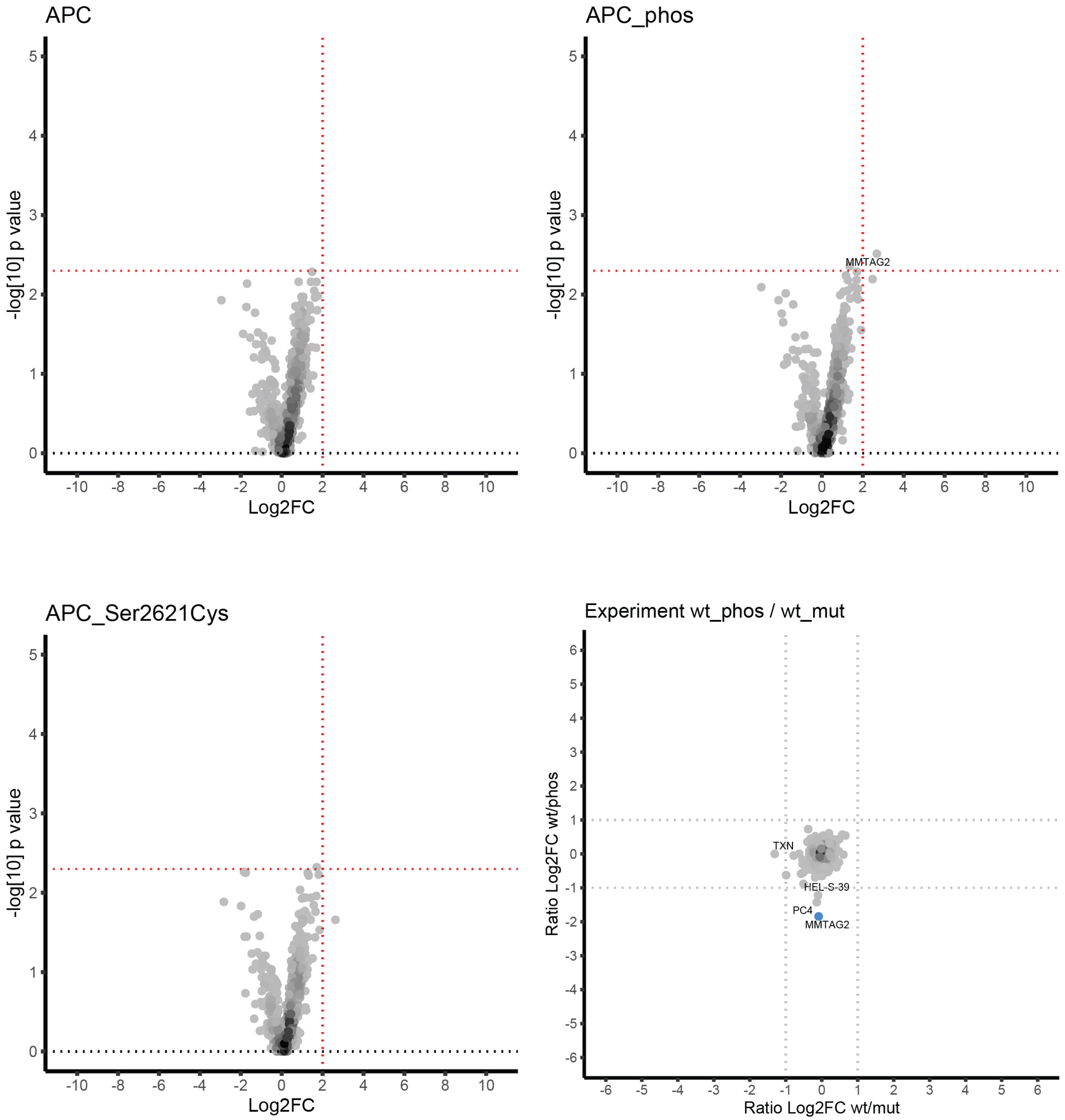

### COL5A1: GRQGPKGpSIGFPGFP

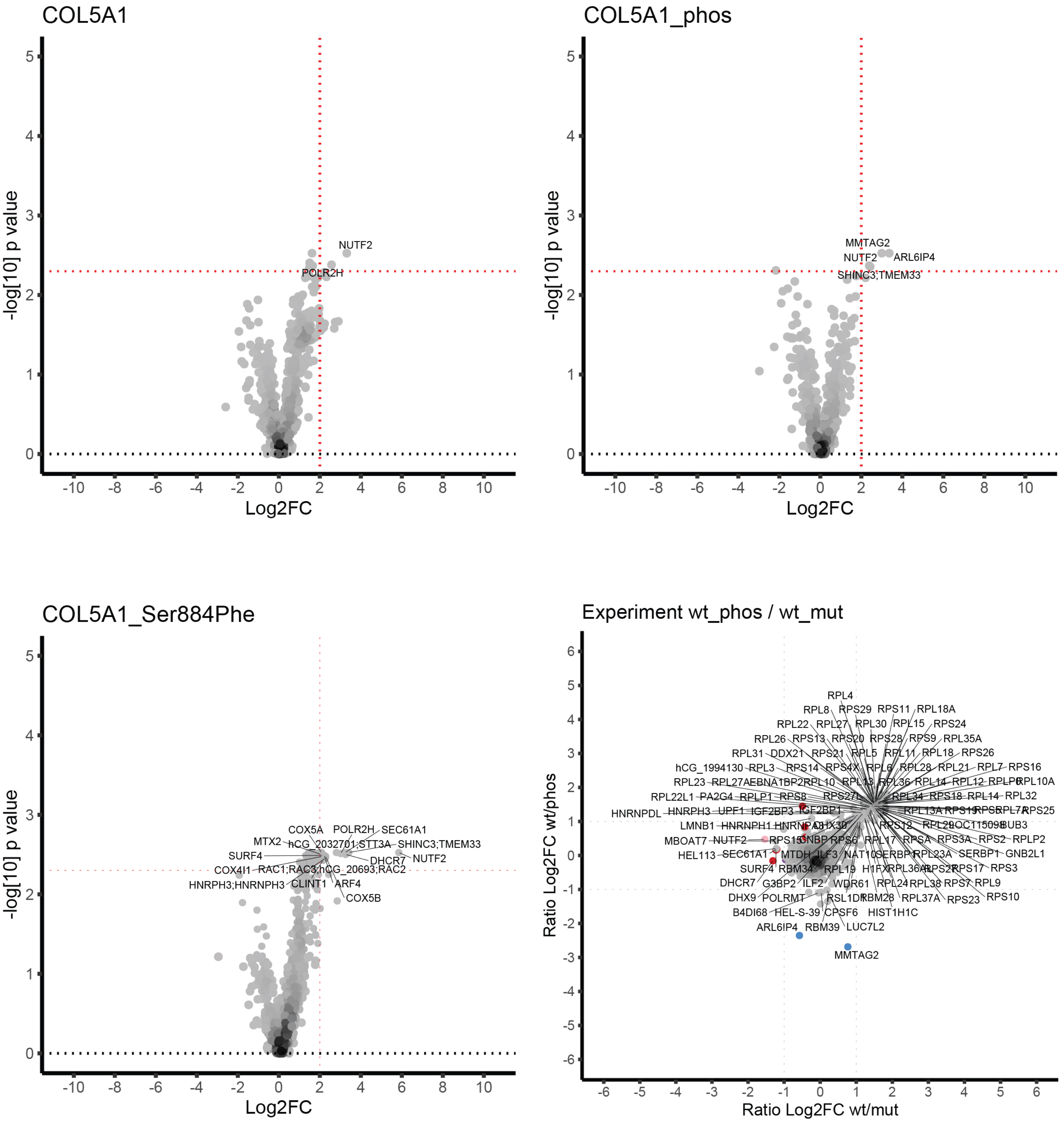

### DENND2A: PPPPLPSpSPPPSSVN

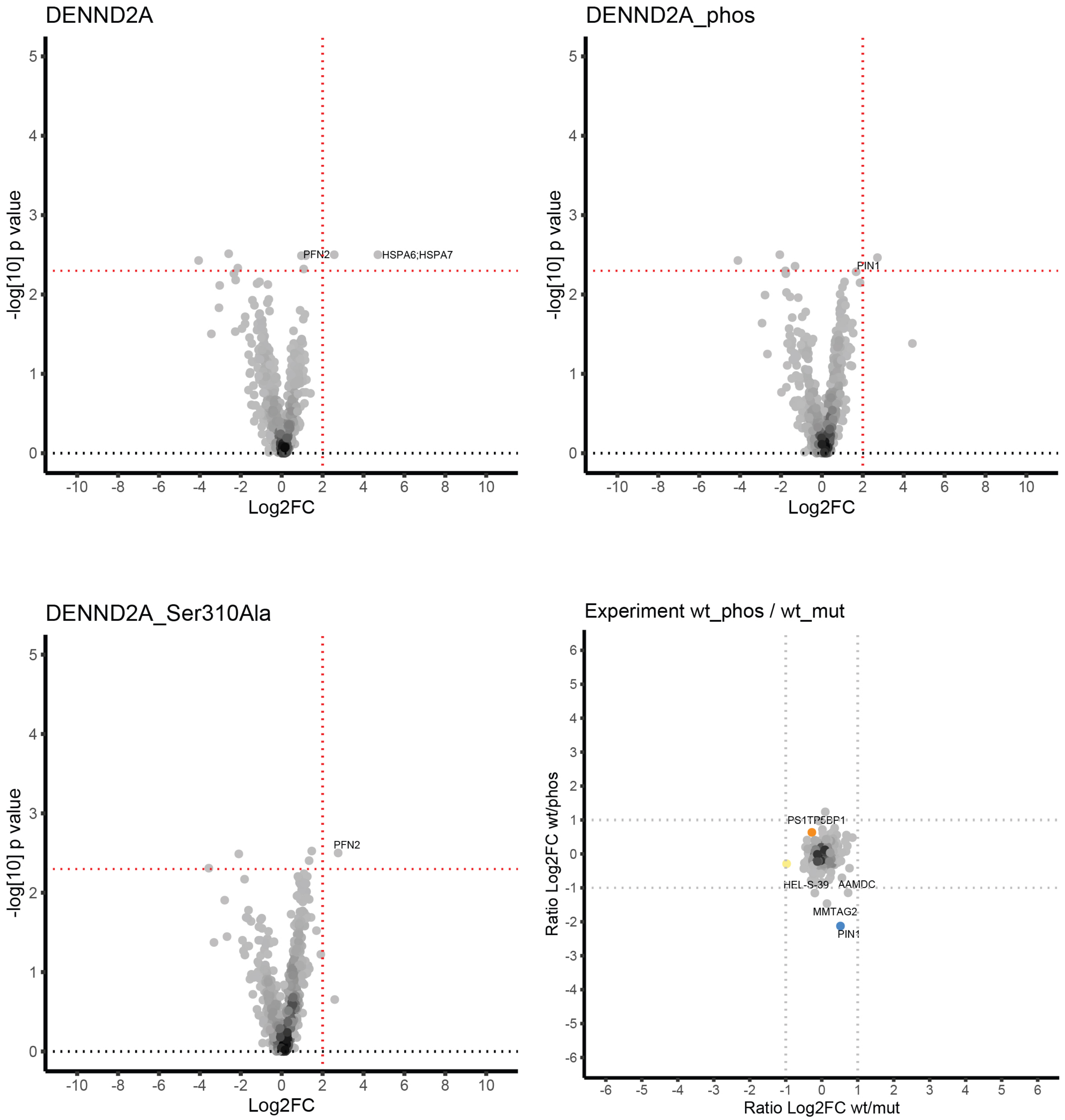

### DES: QRGSEVHpTKKTVMIK

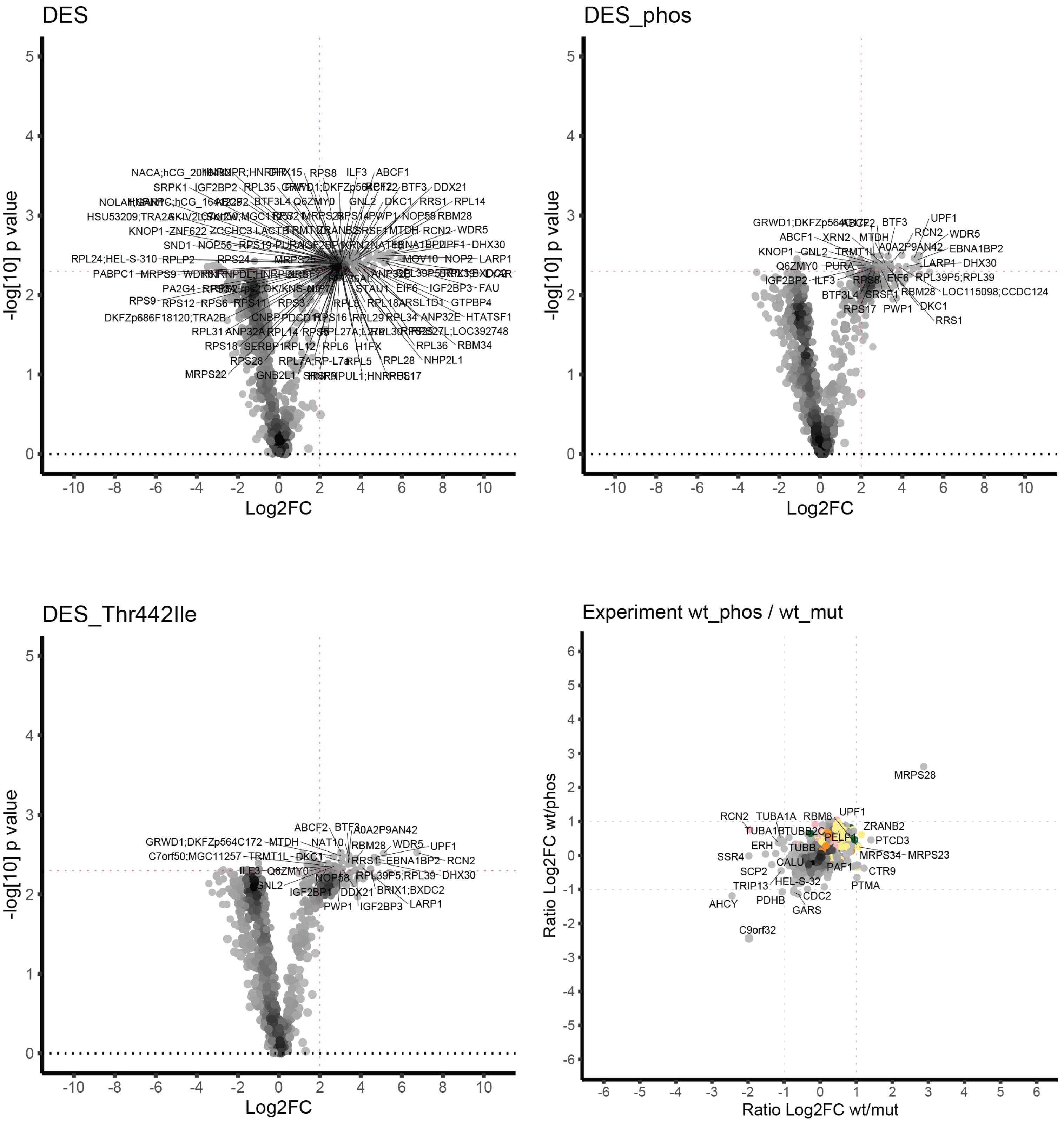

### DIDO1: PGRLGAMpSAAPSQPN

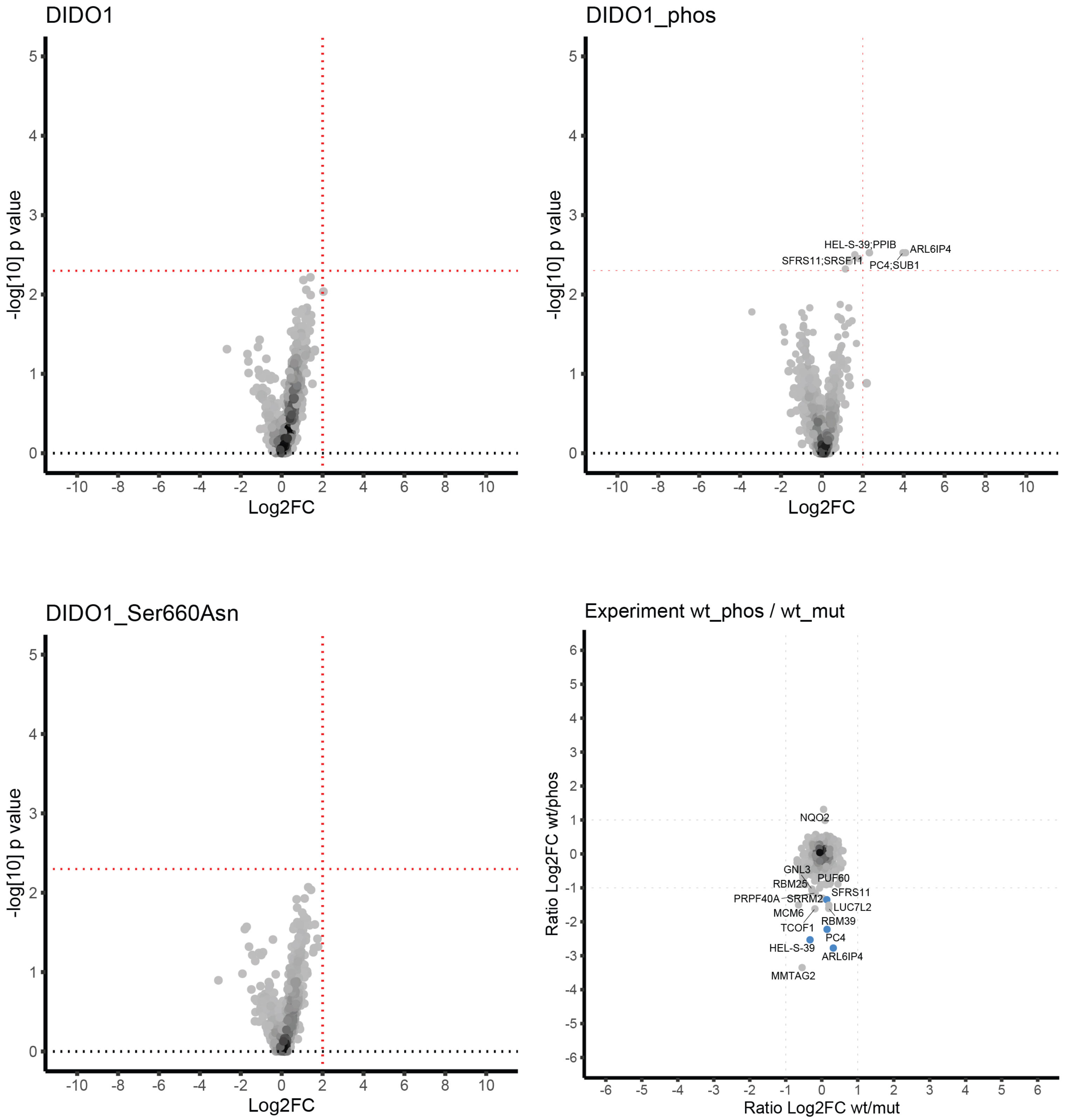

### EGFR: TFLPVPEpYINQSVPK

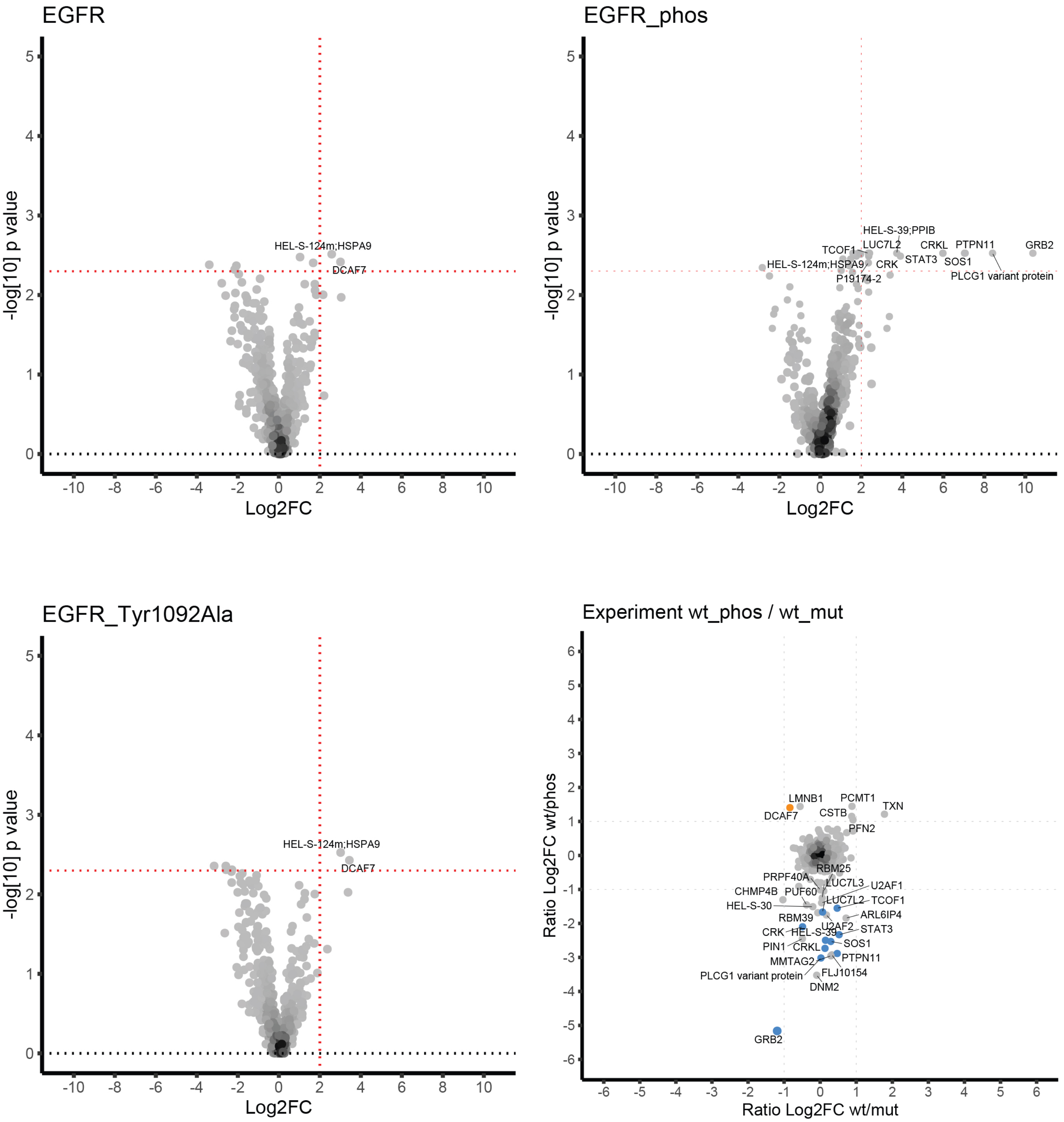

### EID1: MSEMAELpSELYEESS

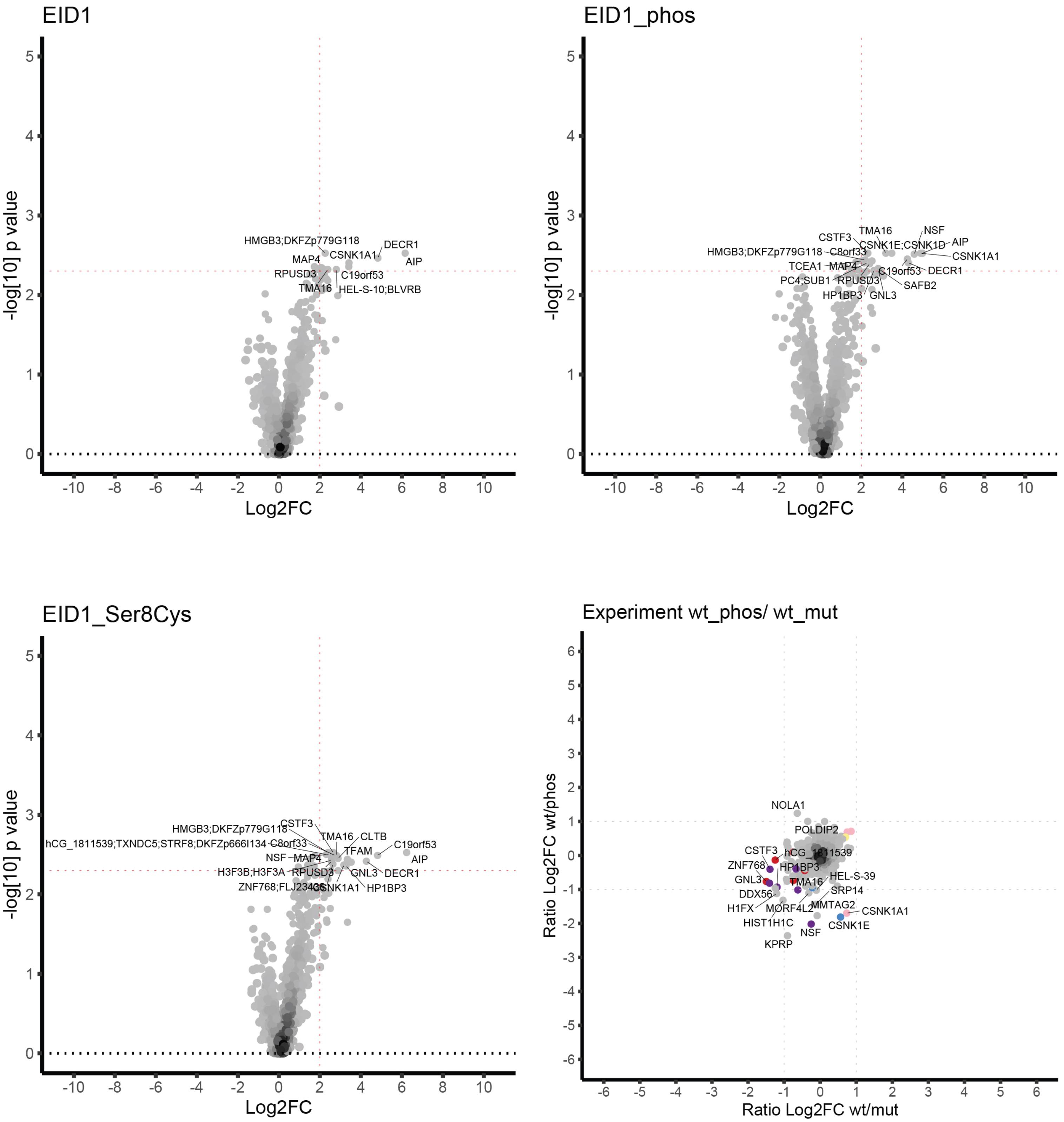

### GATAD1: LRNTKYKpSAPAAEKK

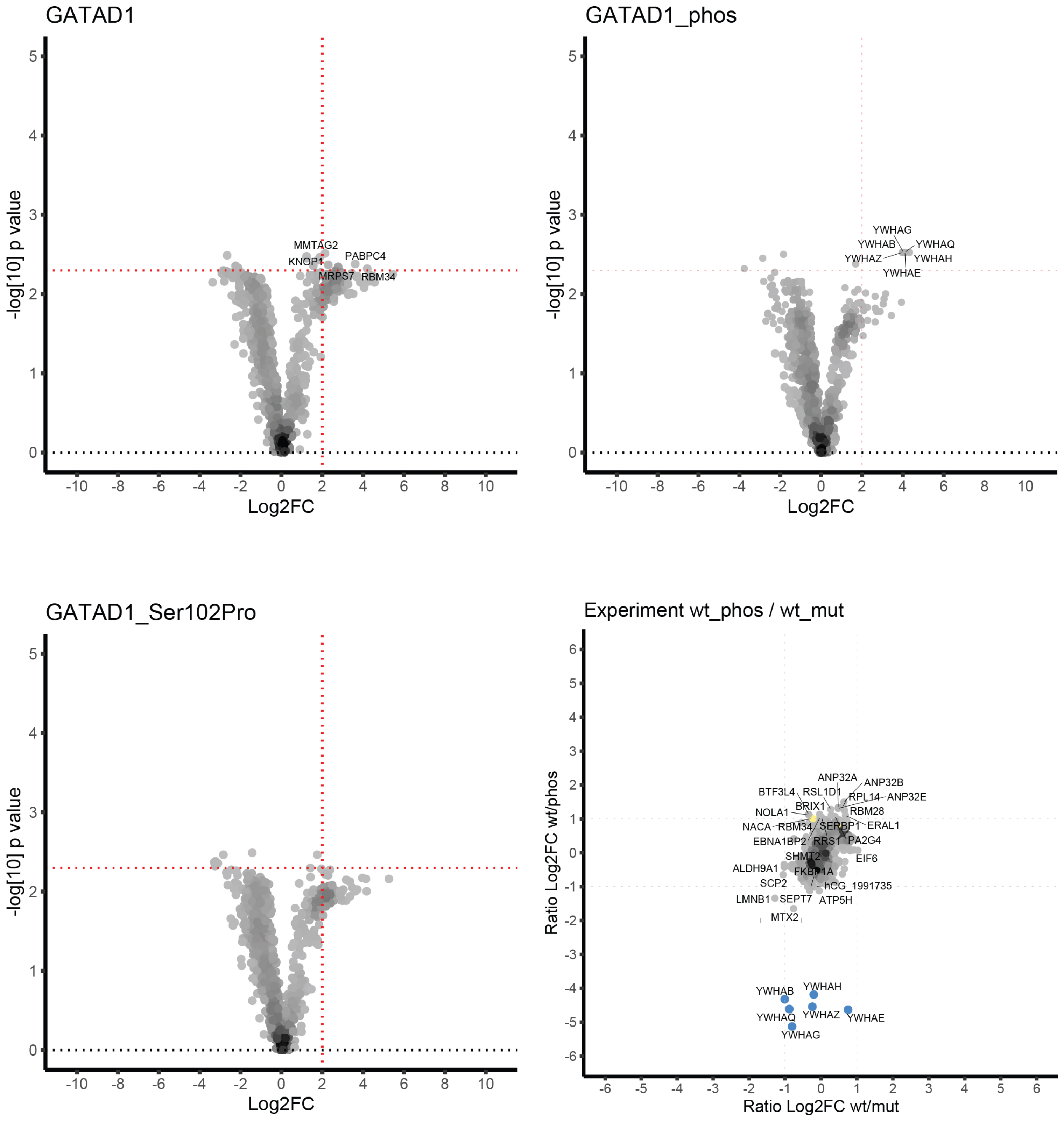

### GORASP1: HERGGEApTWSGSEFE

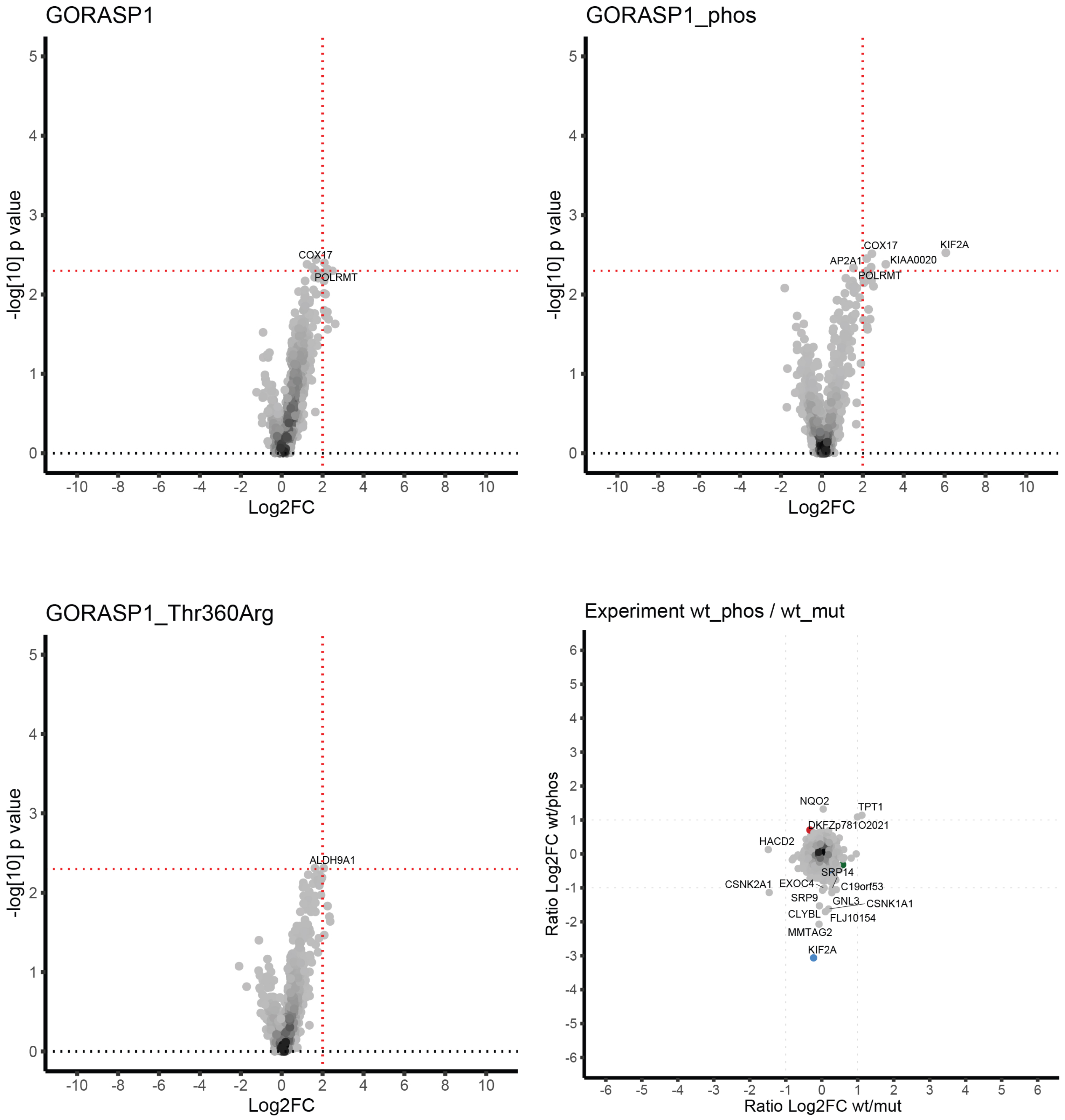

### GPR158: RSIMRRIpTEIPETVS

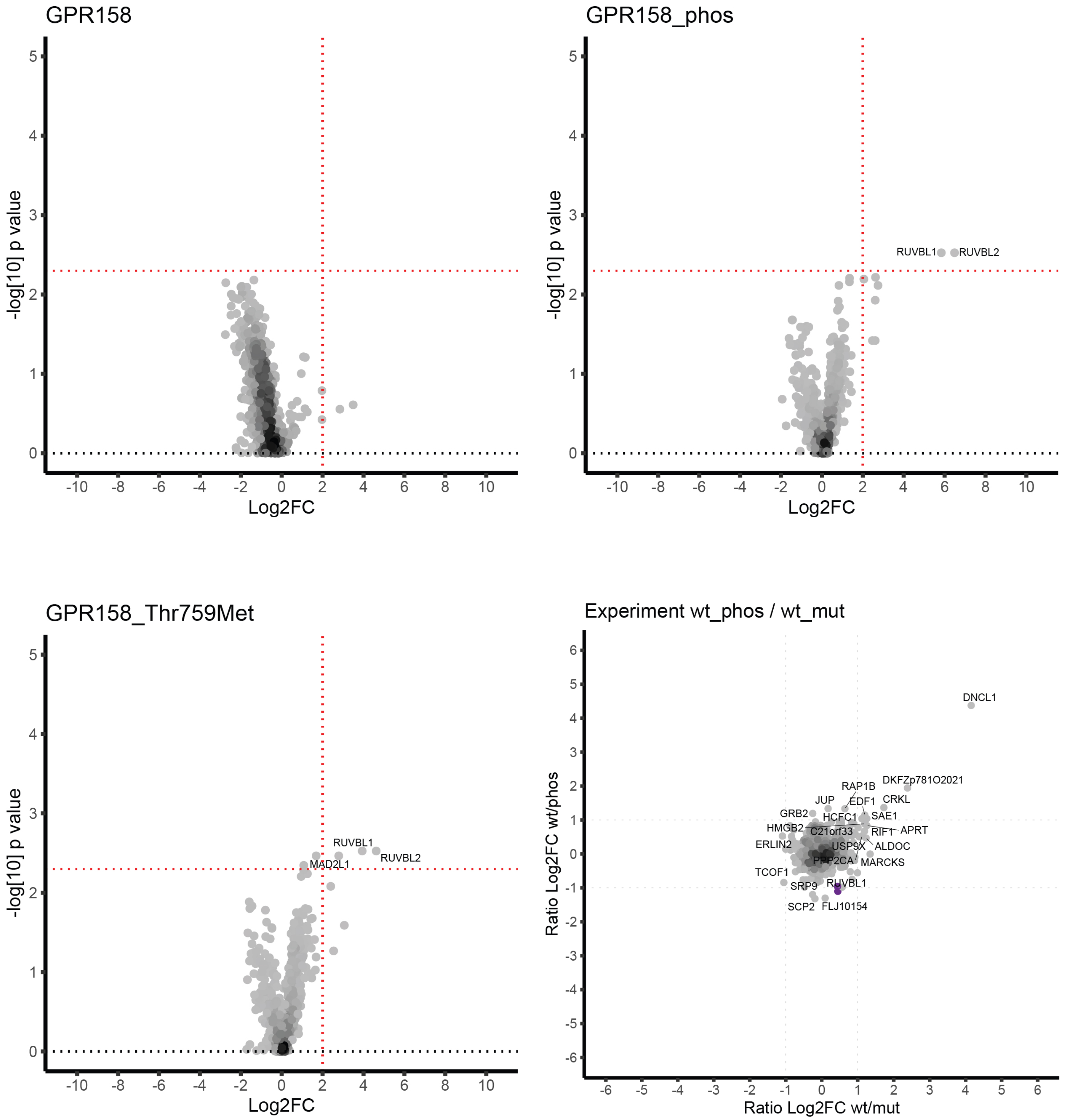

### HDAC4: EPGQRQPpSEQELLFR

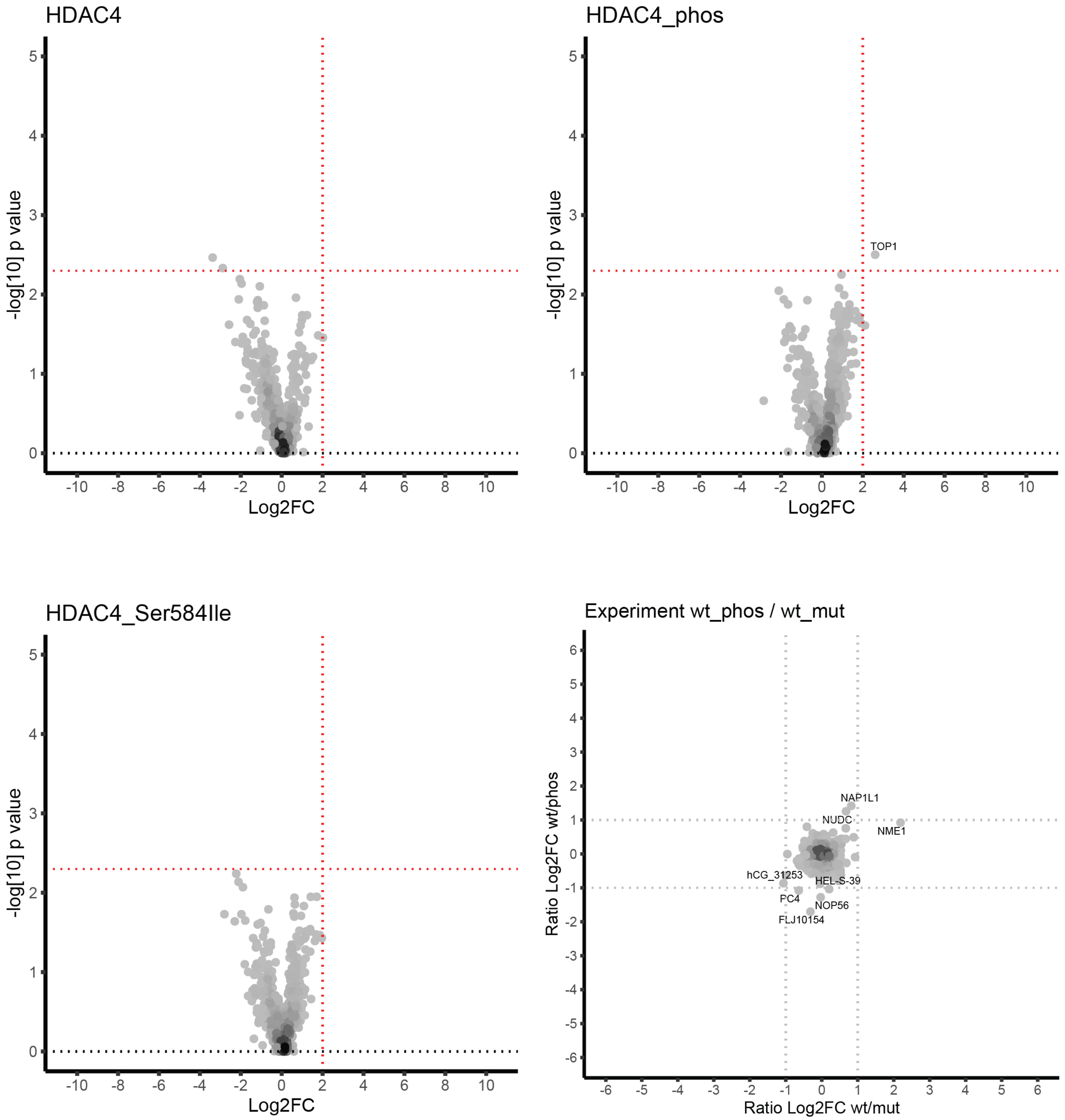

### LRRC16A: SPKPLLQpSPKPSLAA

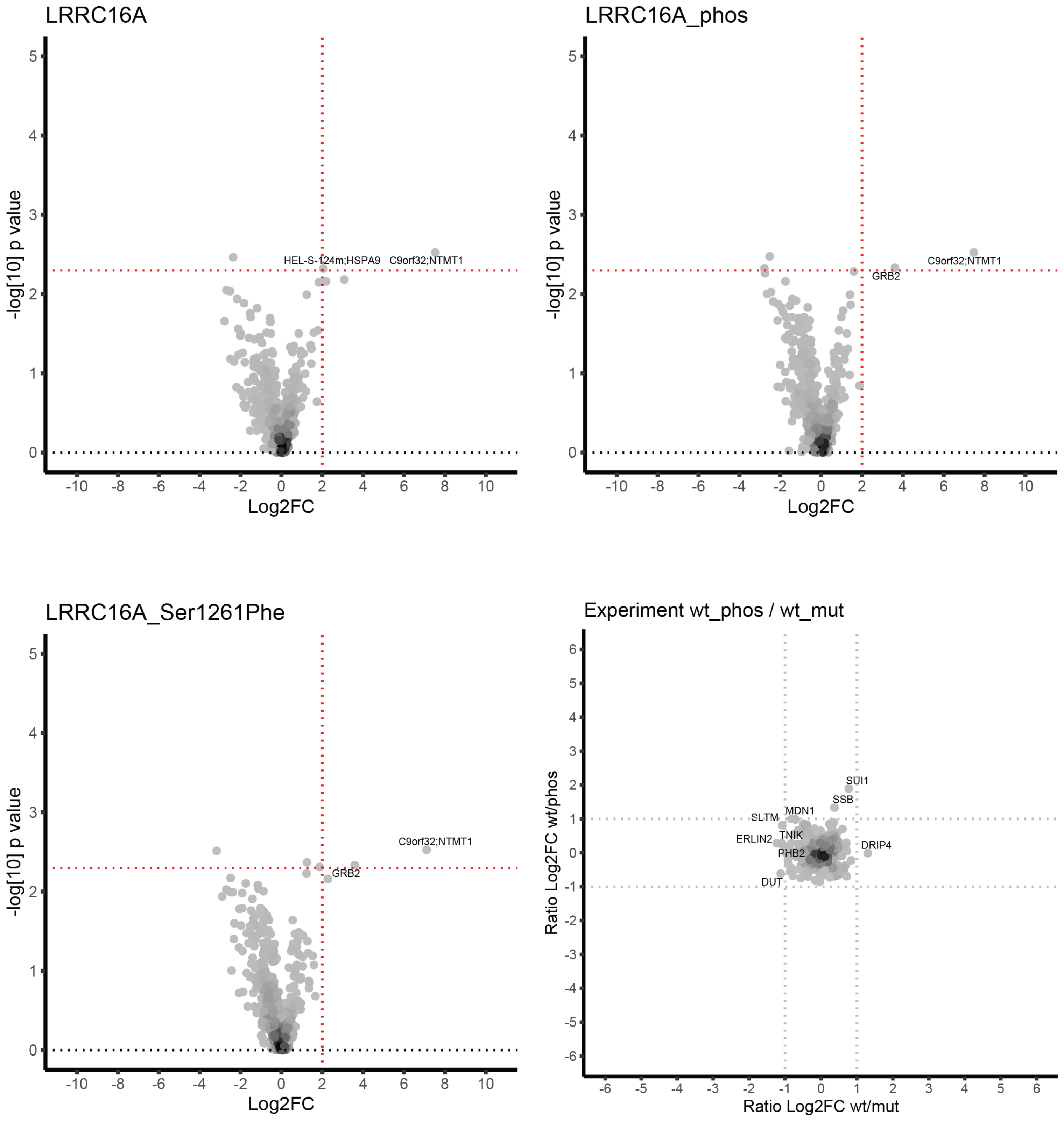

### MAFB: PLSTPCSpSVPSSPSF

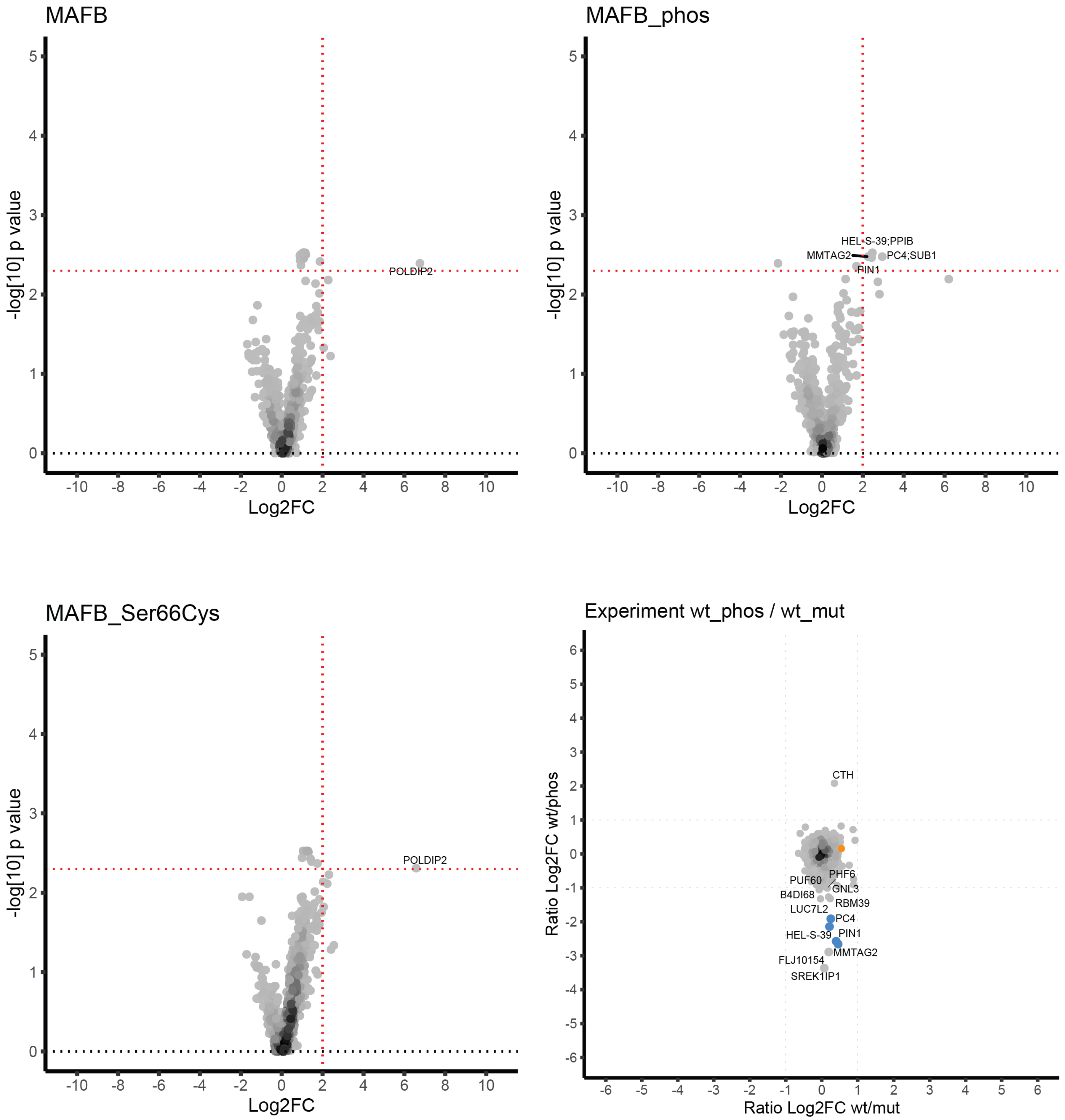

### MAFB: TPCSSVPpSSPSFSPT

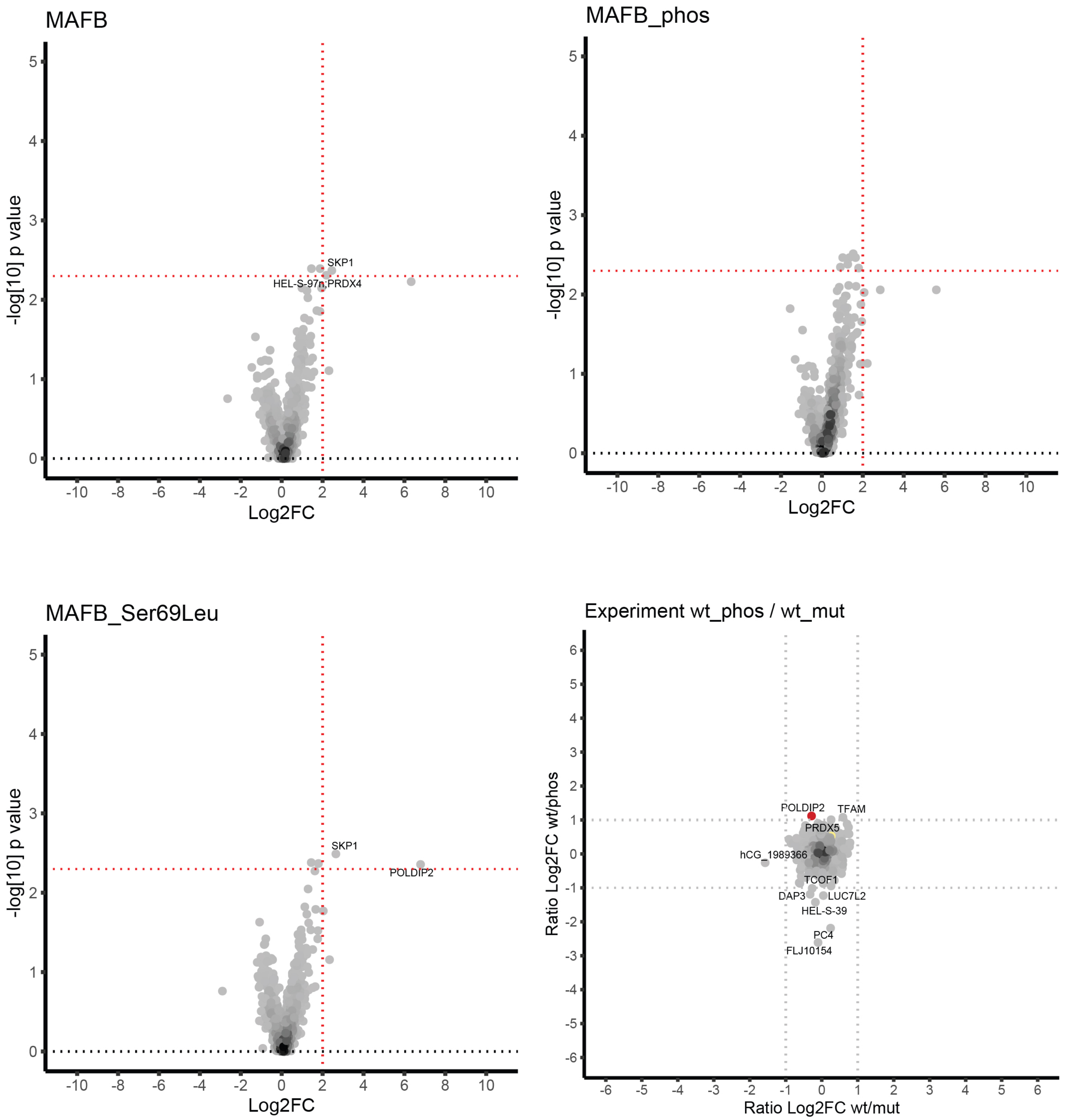

### MAFB: PCSSVPSpSPSFSPTE

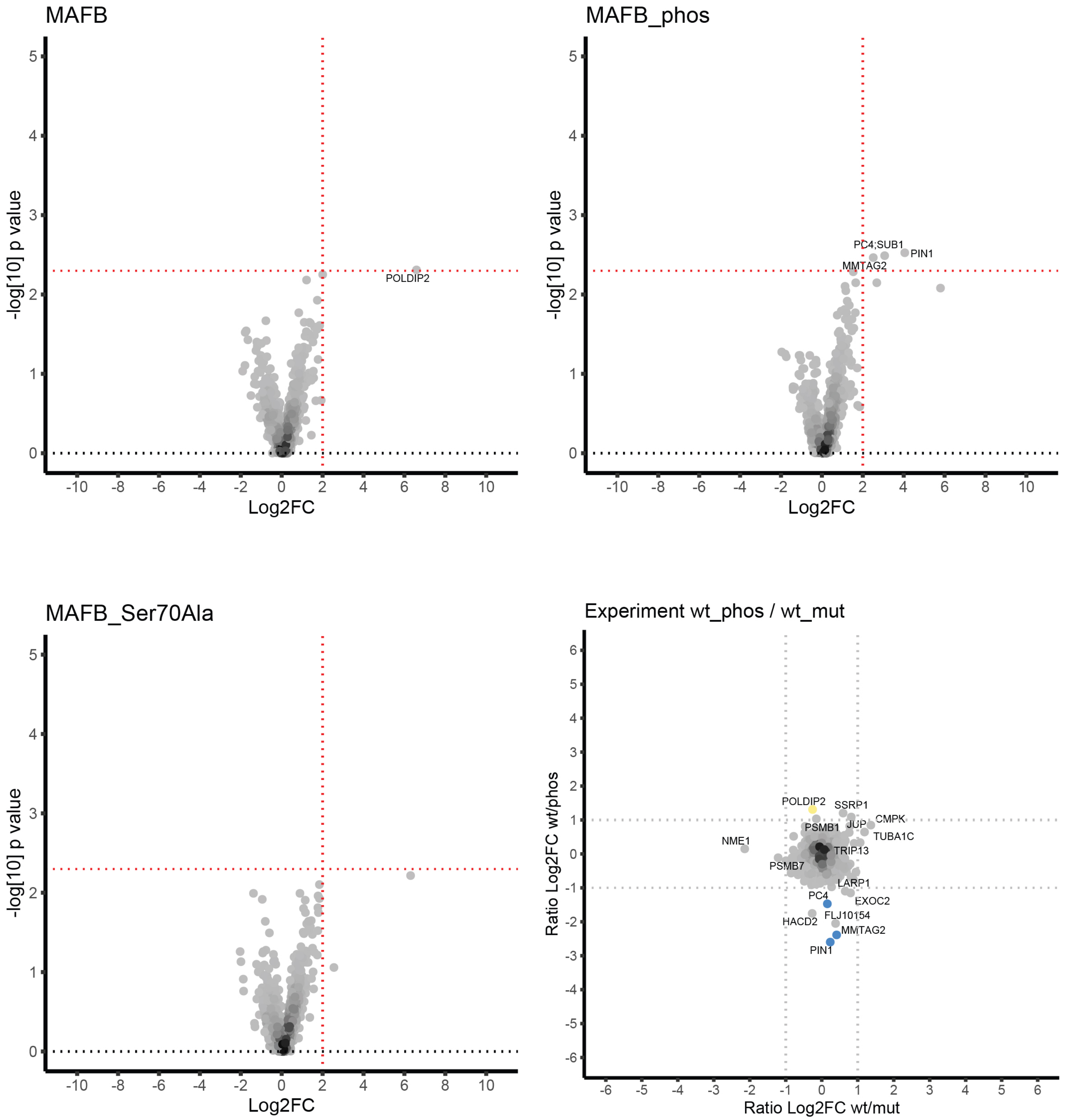

### MAFB: VSSTPLSpTPCSSVPS

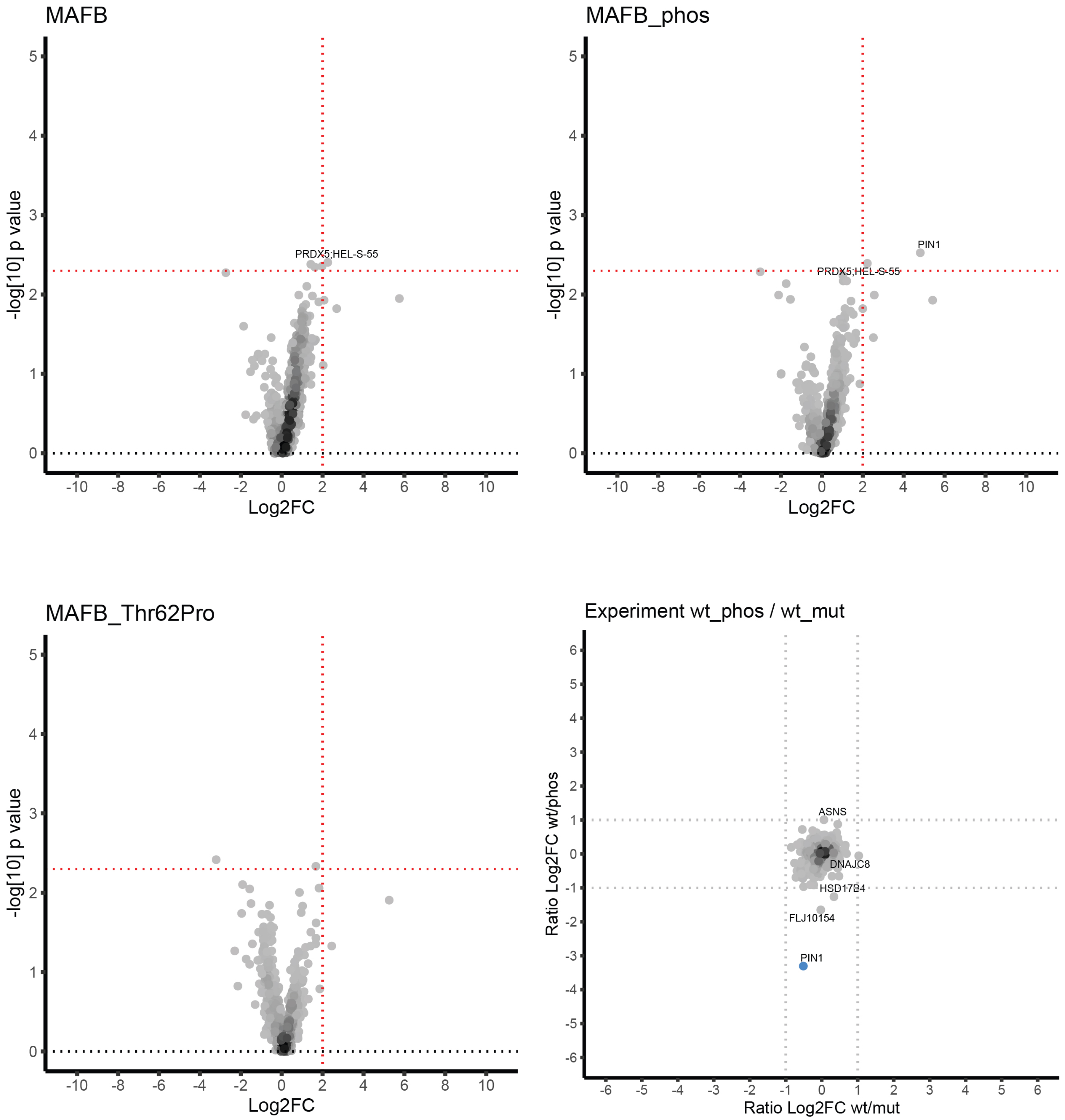

### MLH1: SGSSDKVpYAHQMVRT

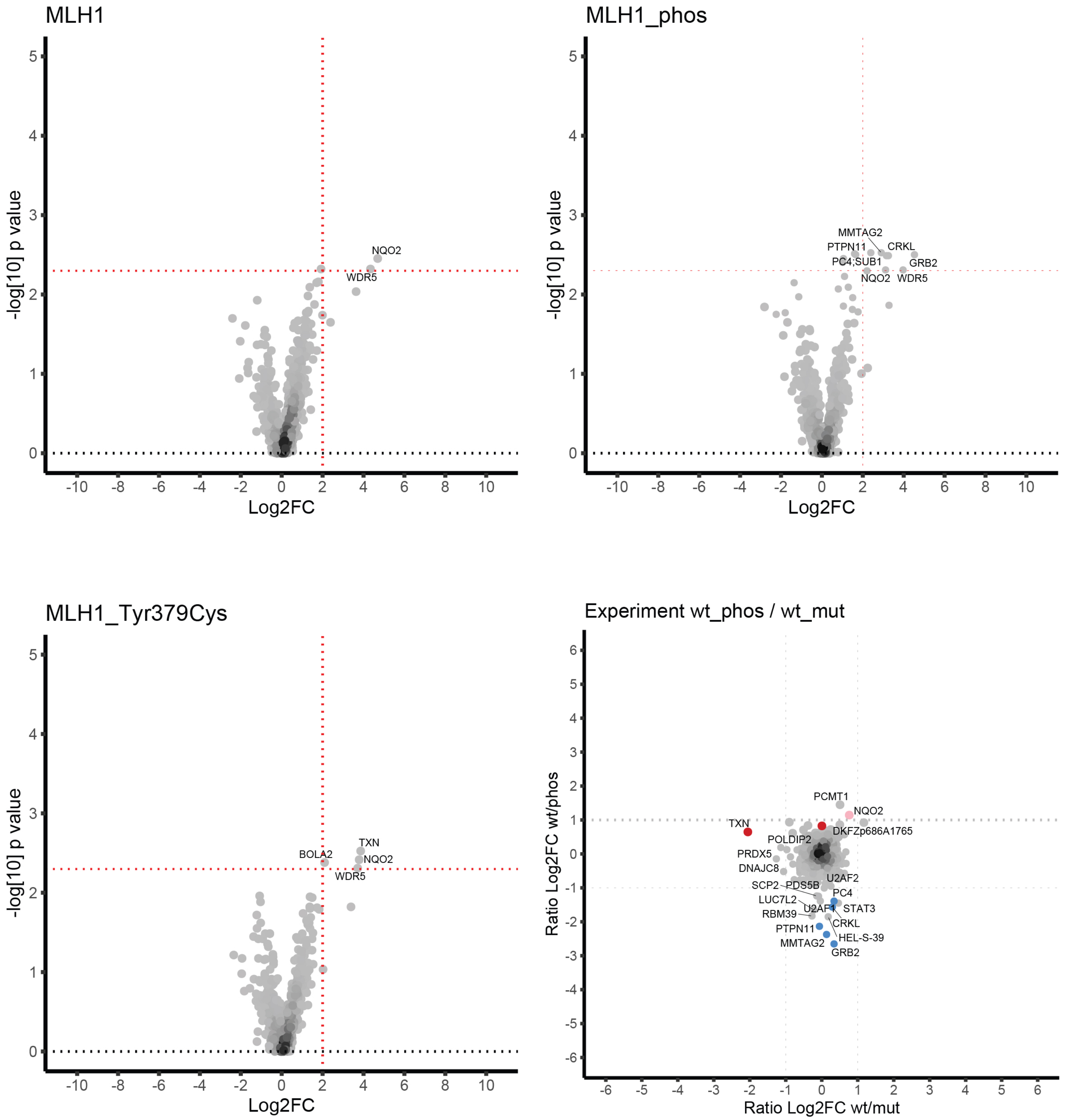

### MMAA: LSRDMNApYIRPSPTR

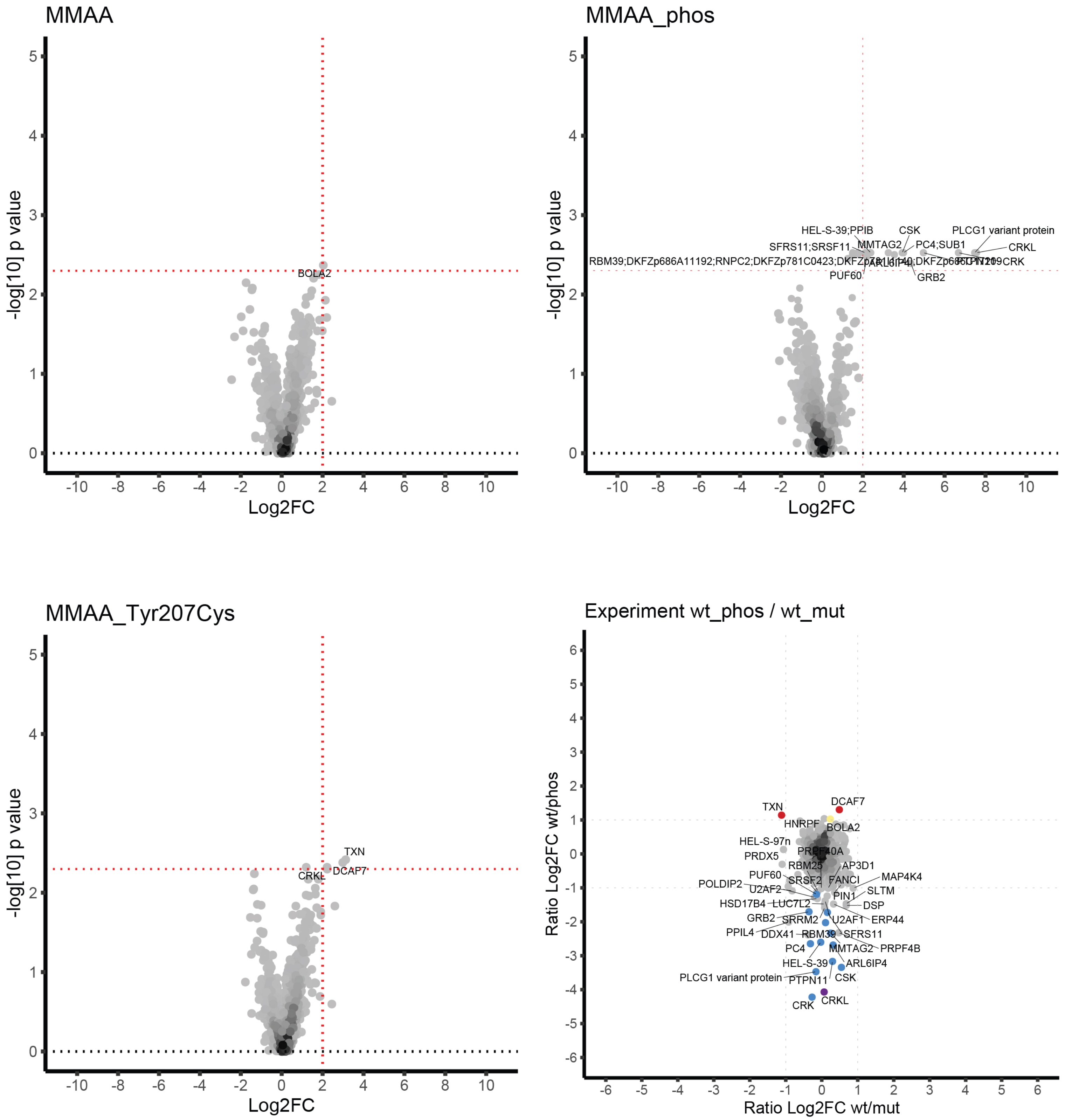

### MYBPC3: GPGAPVTpTTEPVTVQ

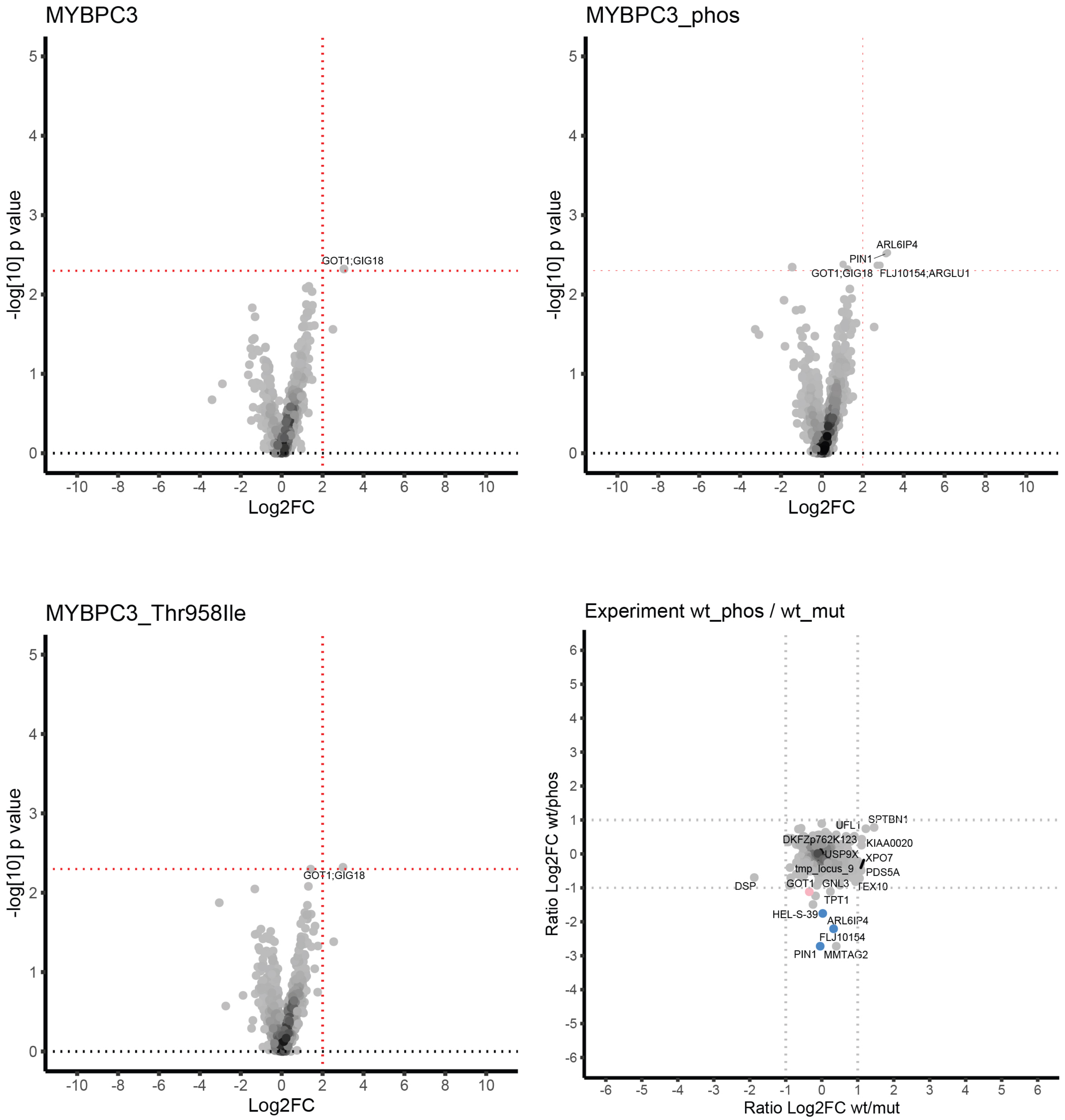

### MYT1L: SNSLEDDpSDKNENLG

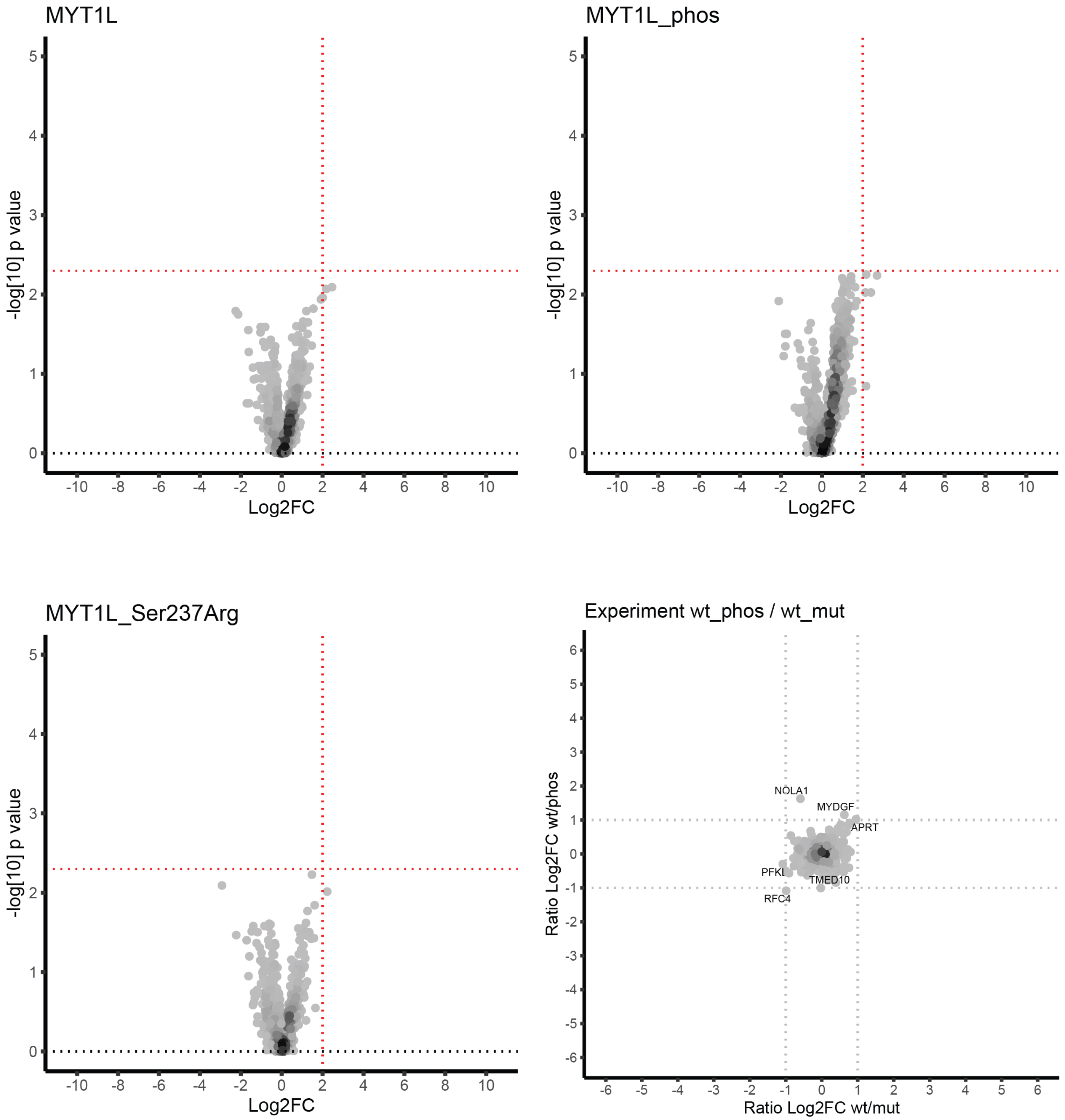

### NCOA6: PSPGRQNpSKAPKLTL

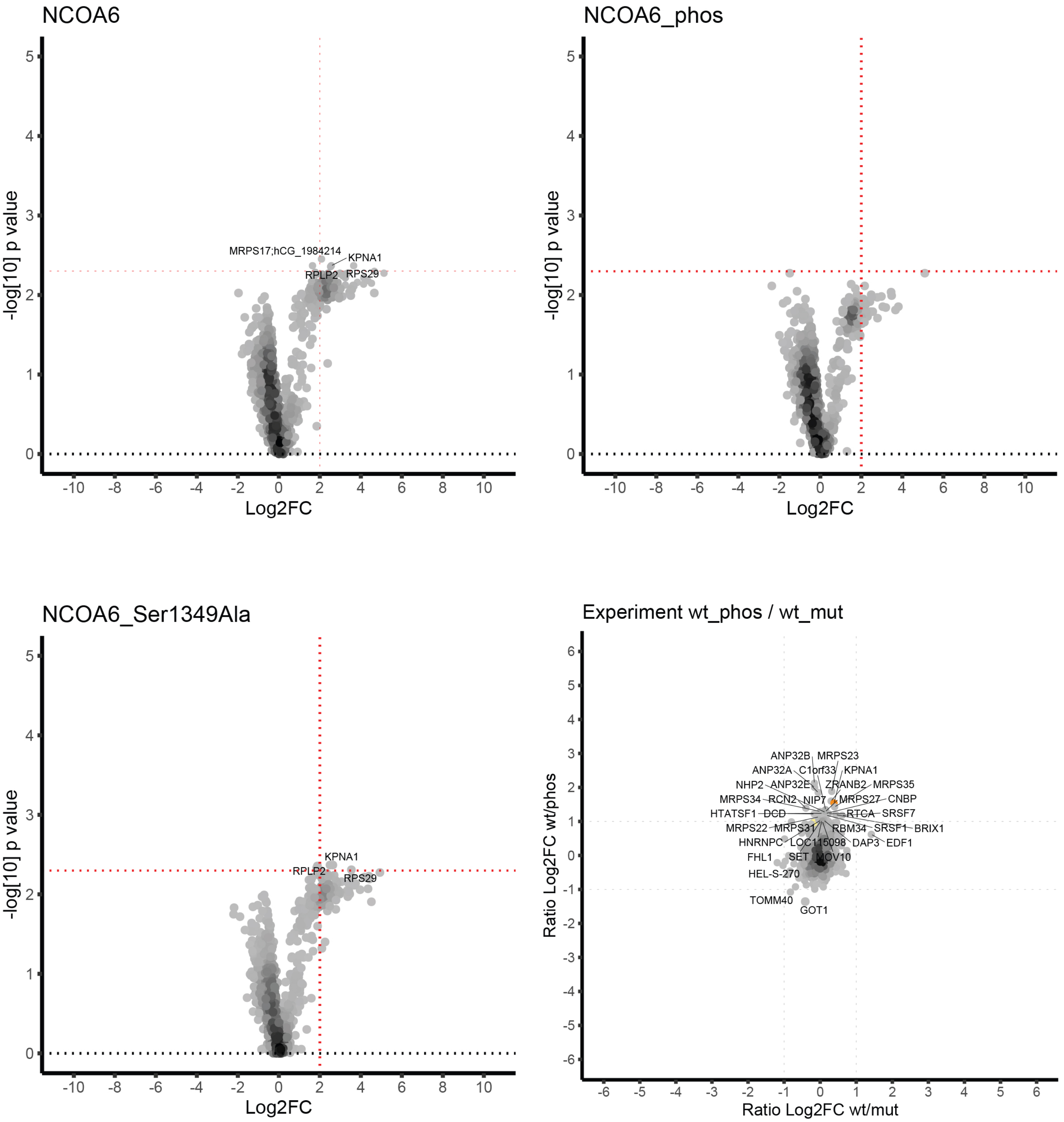

### NRAP: EQPRGKGpSFPAMITP

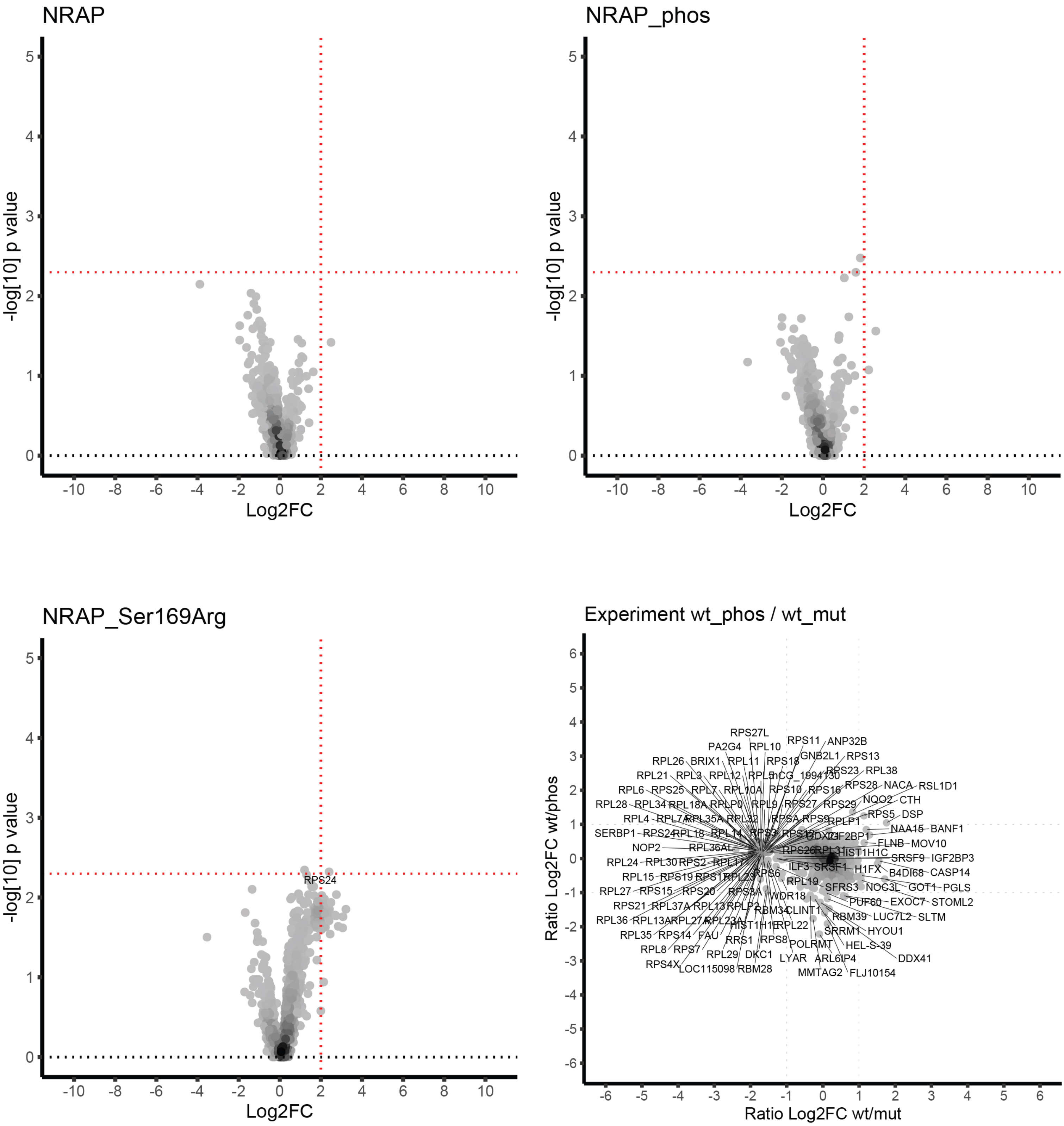

### NRCAM: PSEASEQpYLTKASEP

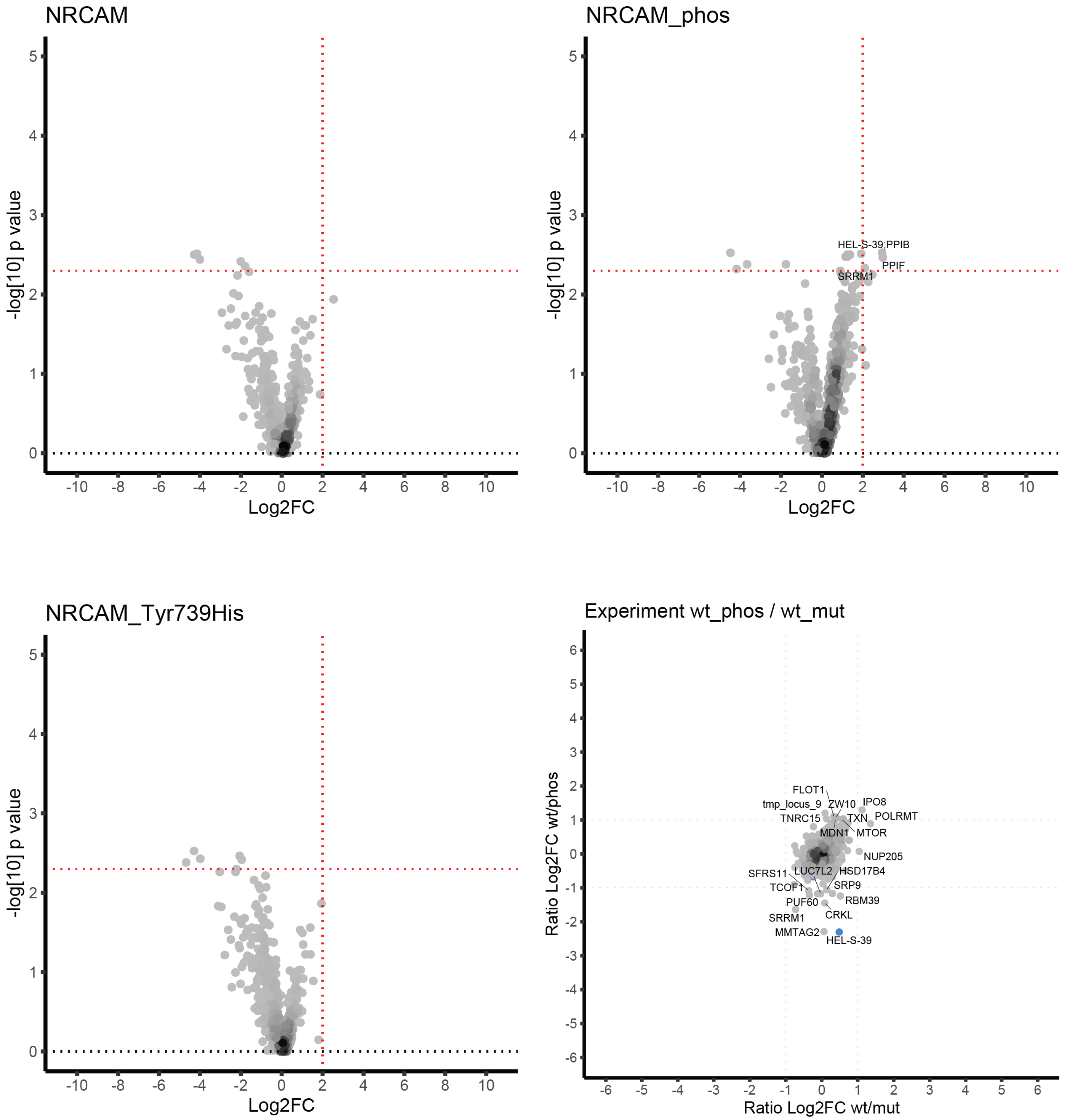

### PHF20: PENDIVKpSPQENLRE

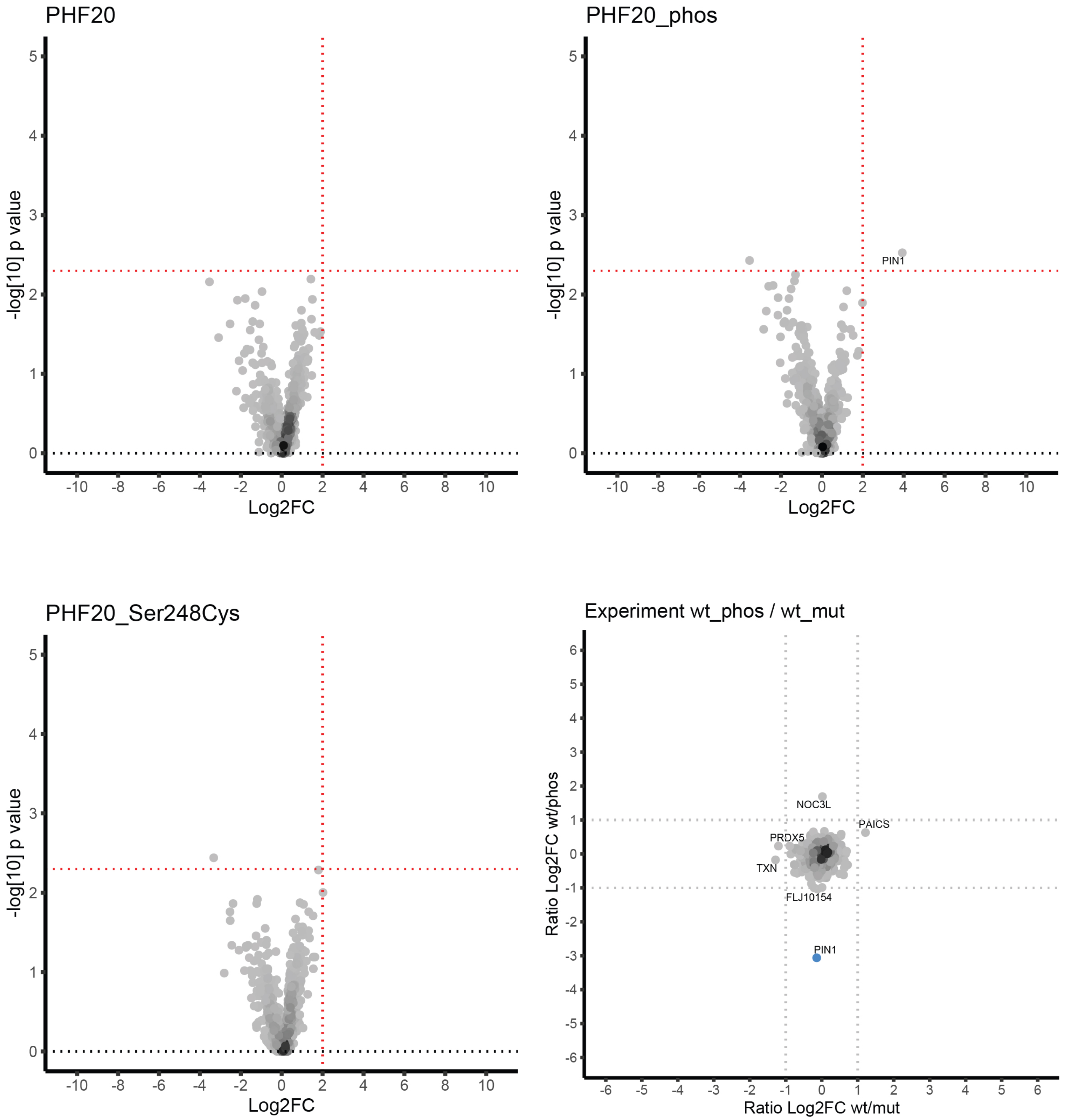

### PITPNM1: SPVEAEGpTPEPGAEA

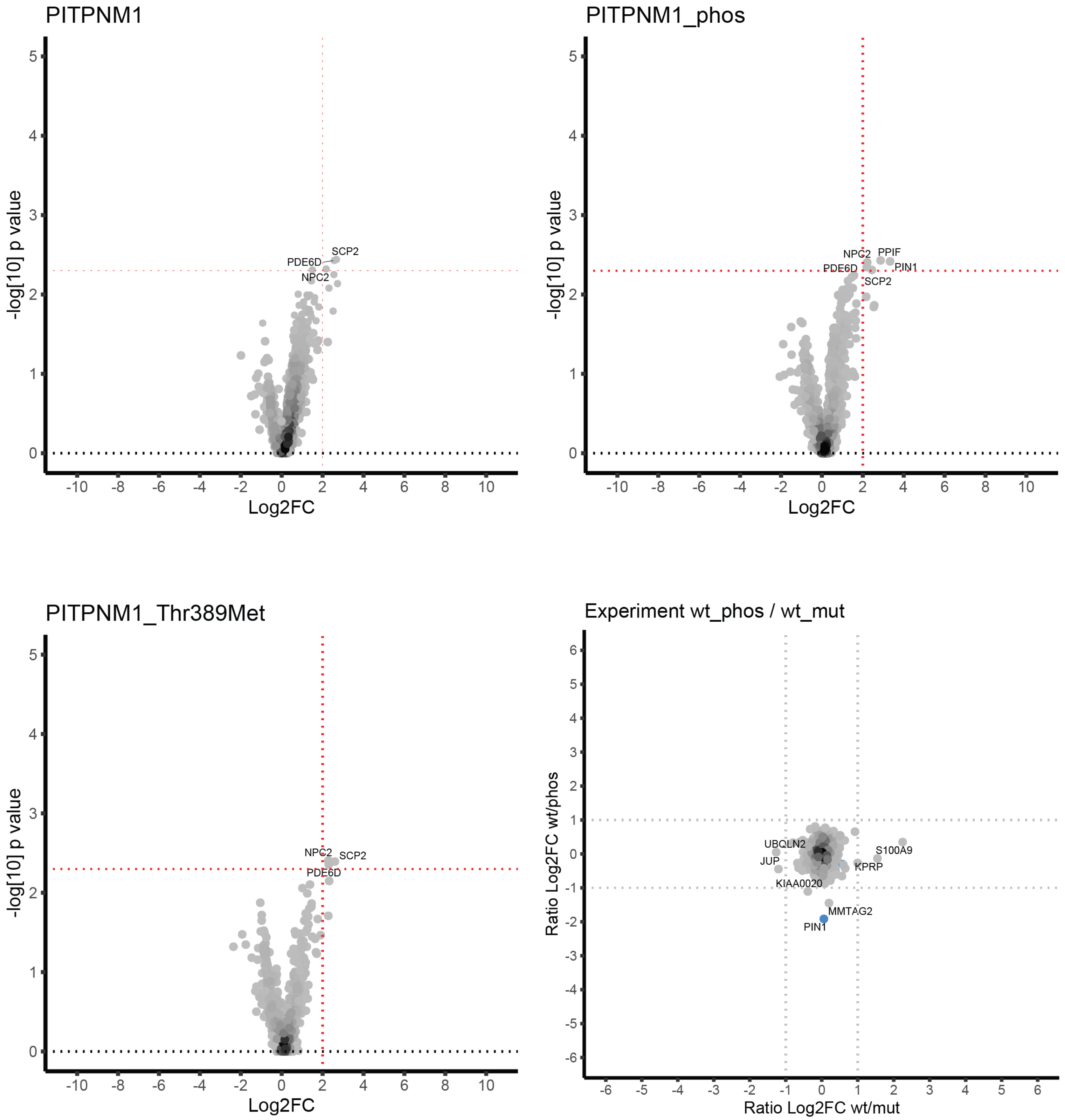

### PPIG: SKQLESKpSNEHDHSK

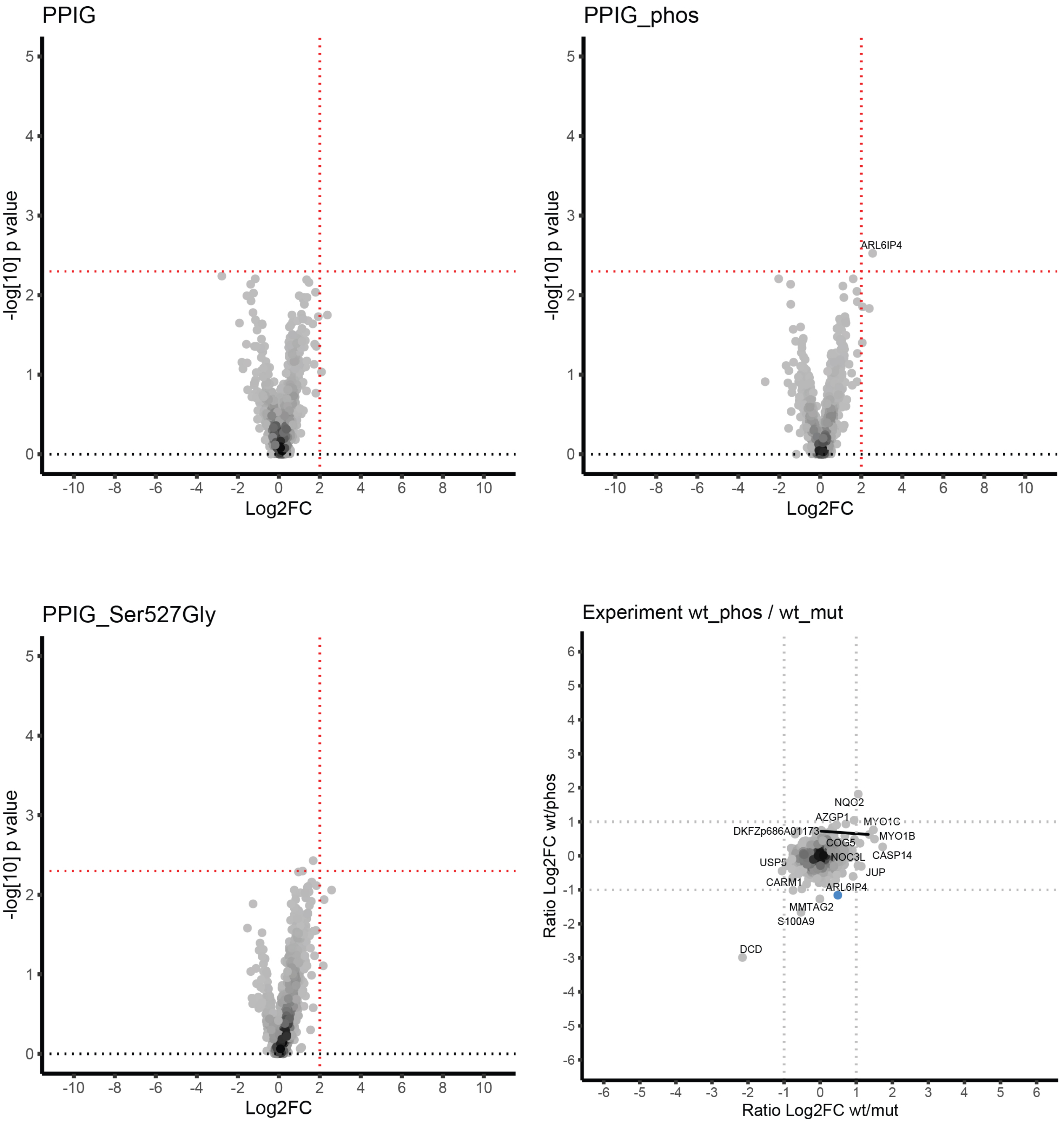

### SCN1A: DEFQKSEpSEDSIRRK

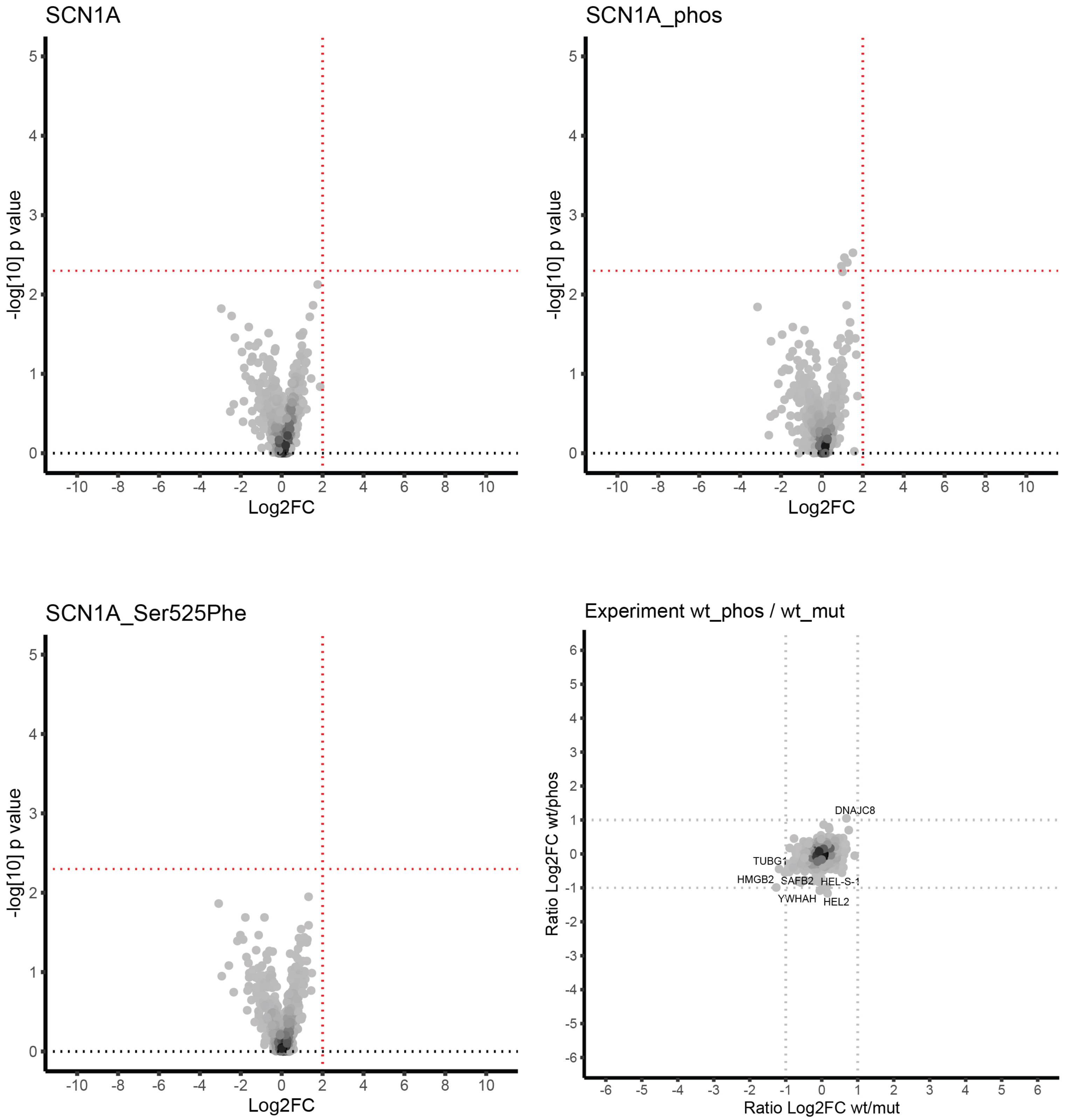

### SCN2A: GQLLPEGpTTTETEIR

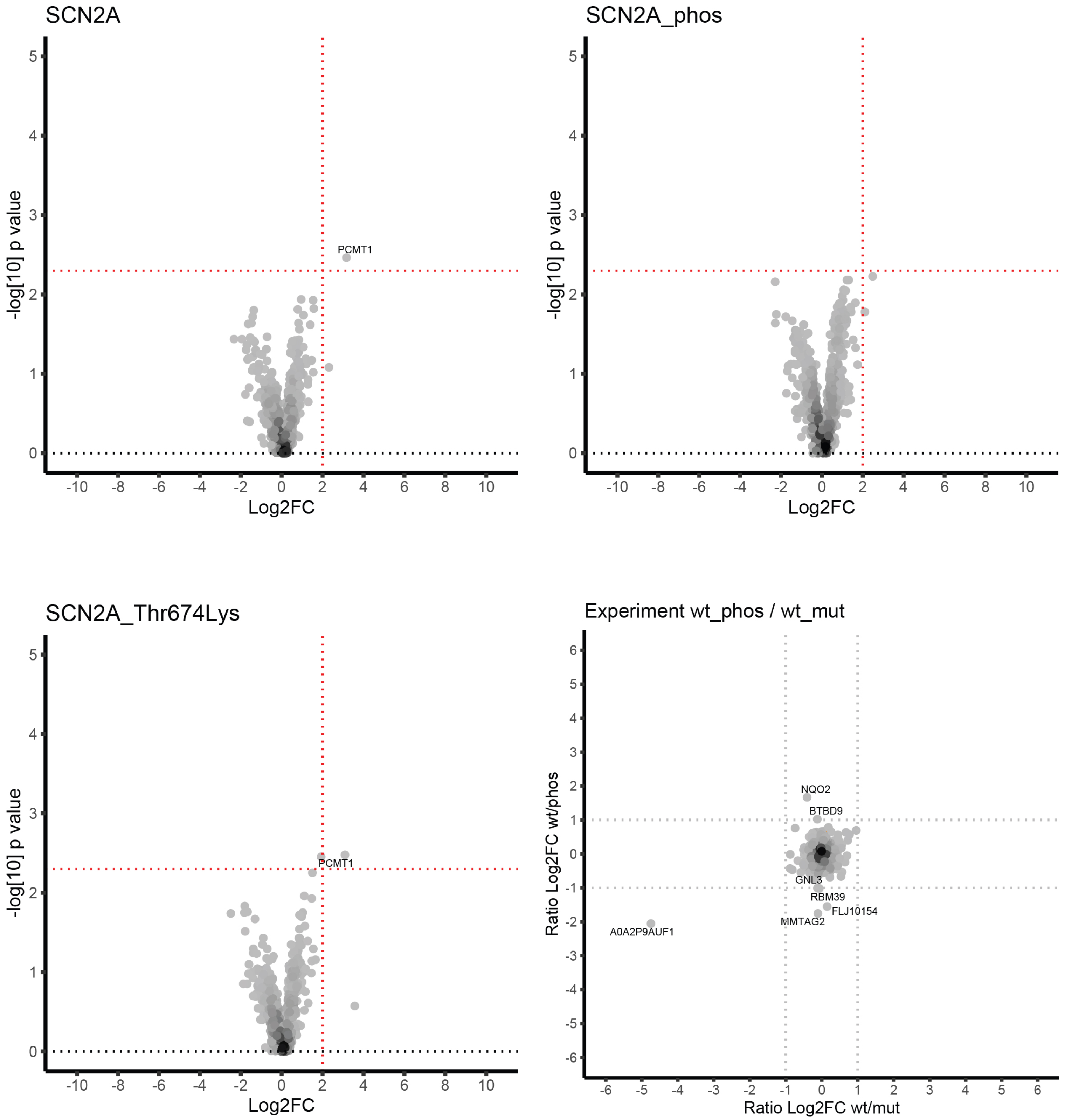

### SPRY4: GAPELAPpTPARCDQD

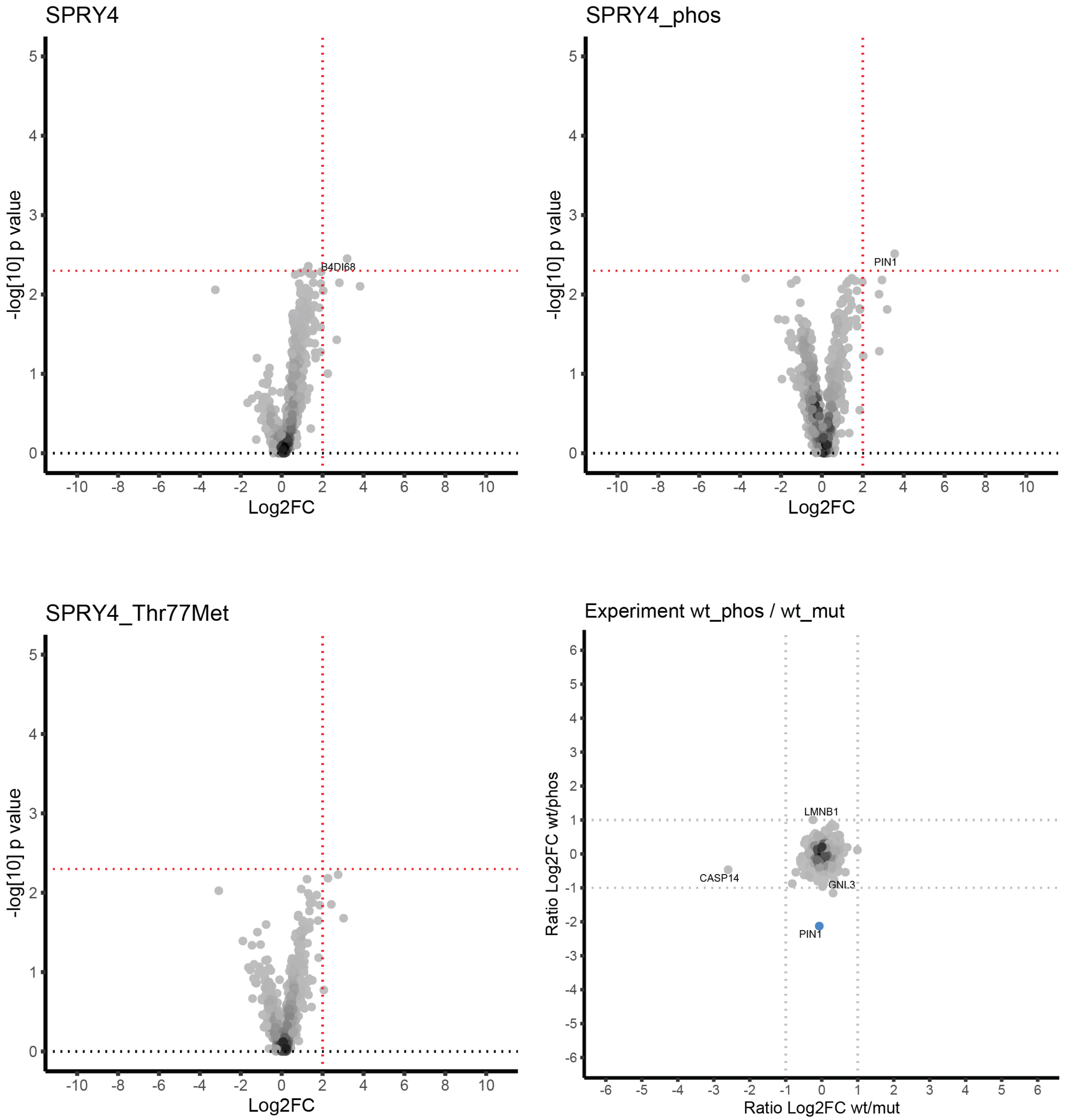

### SRRM1: APEKKEKpTPELPEPS

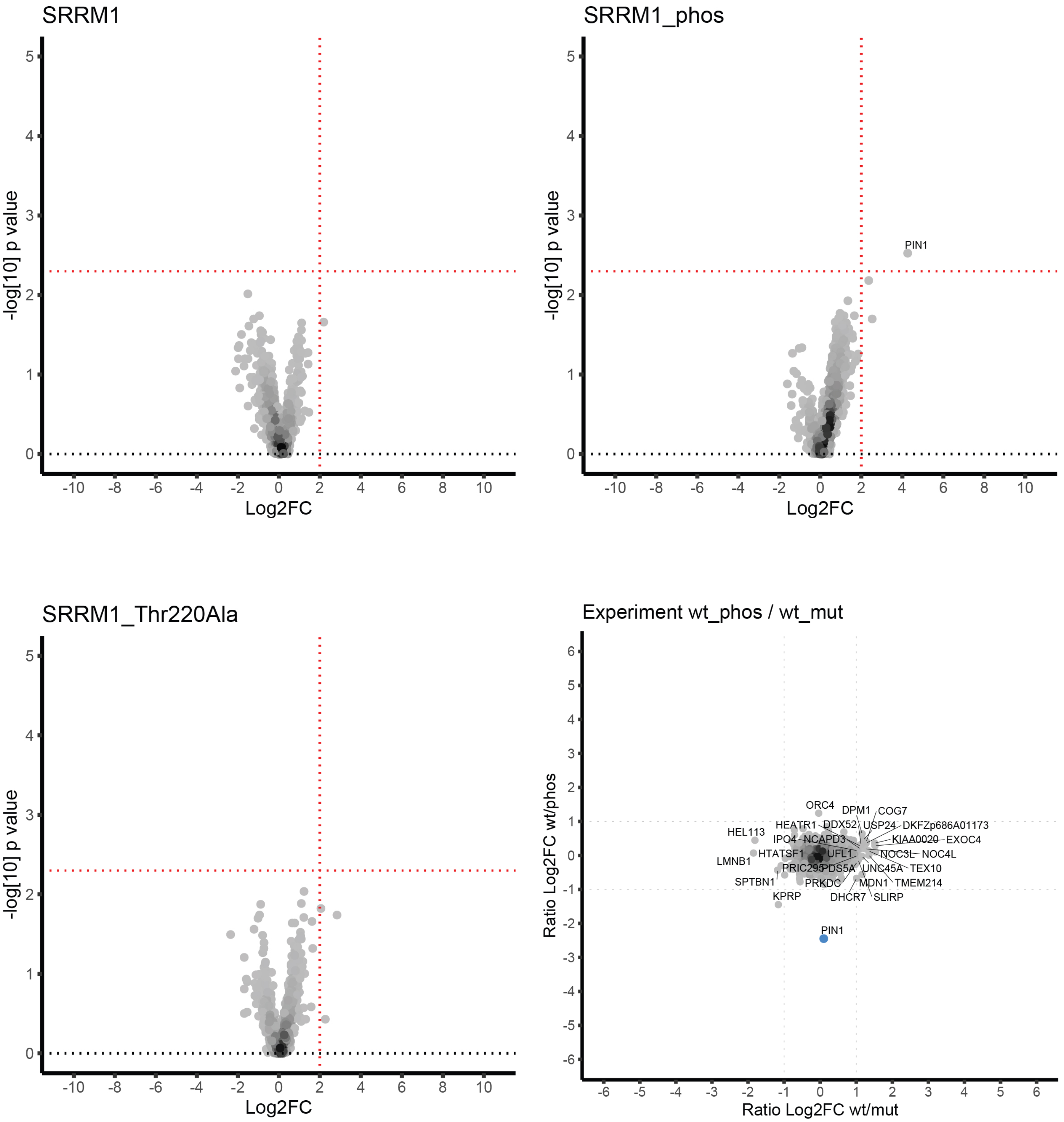

### STXBP5L: QHIPGPGpSIEGMKGA

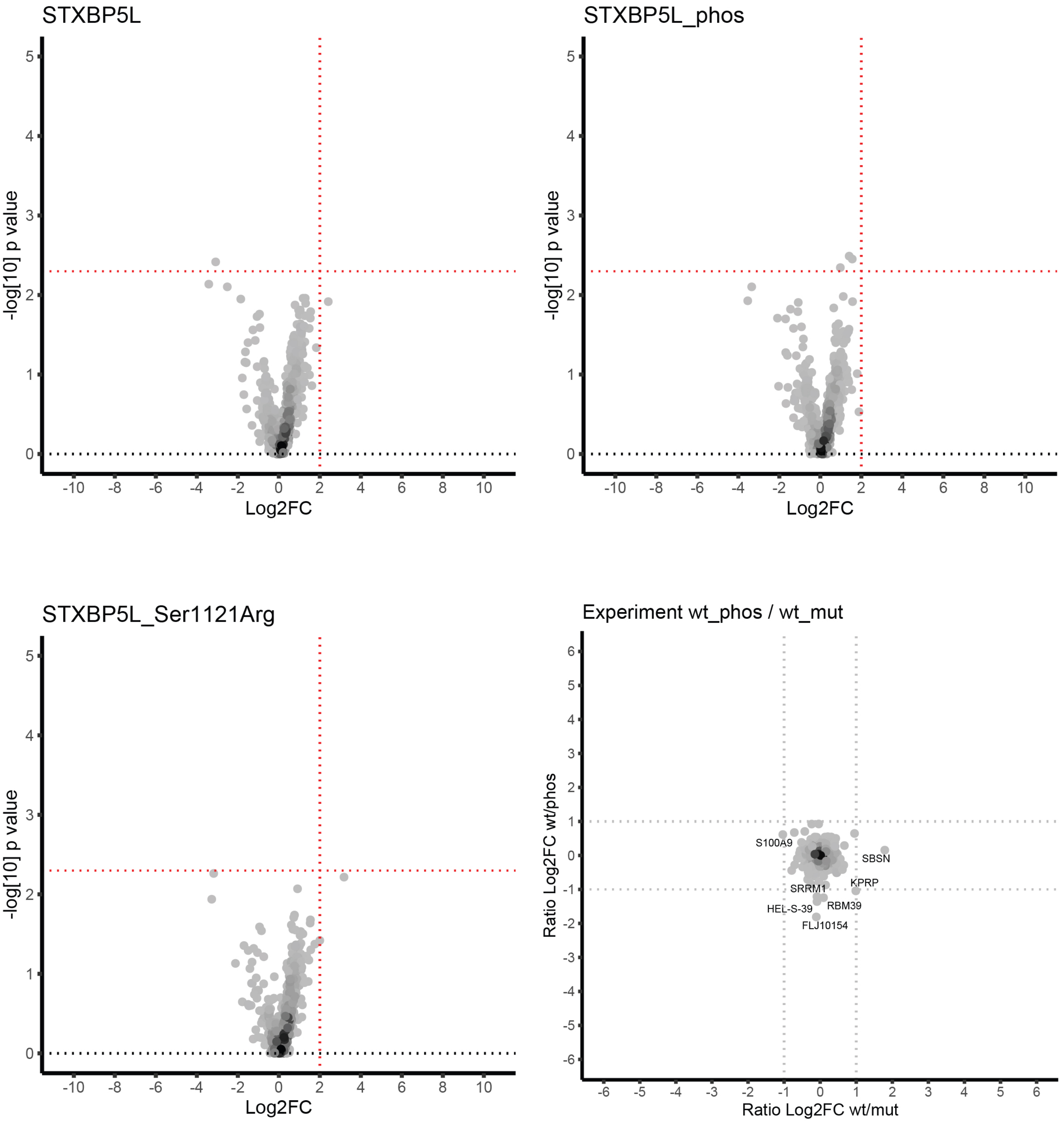

### SYNRG: QETPNECpSDDFGEFQ

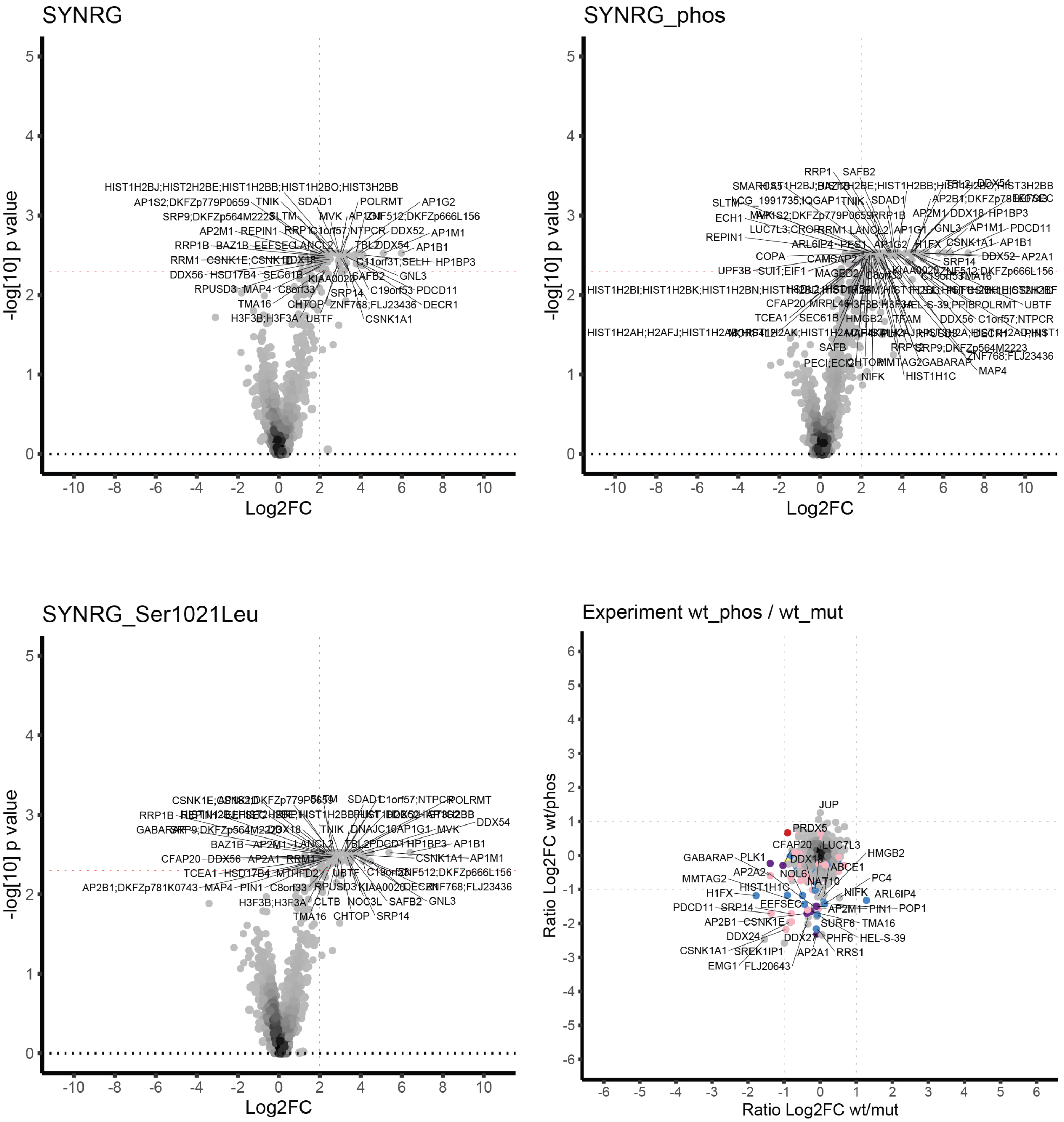

### TNNT2: ARKKKALpSNMMHFGG

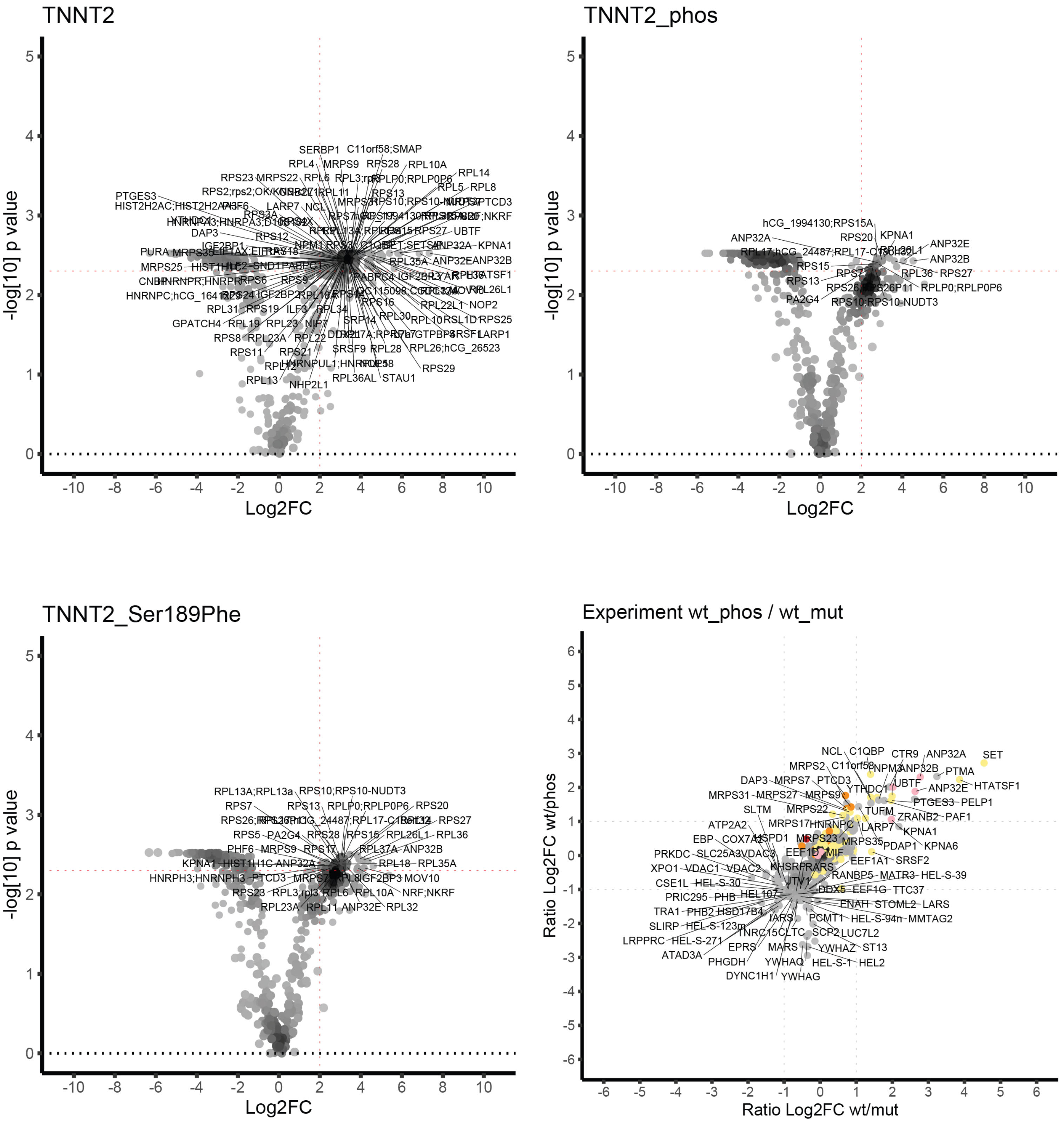

### TTC7A: PDAHDADpSGSRRASS

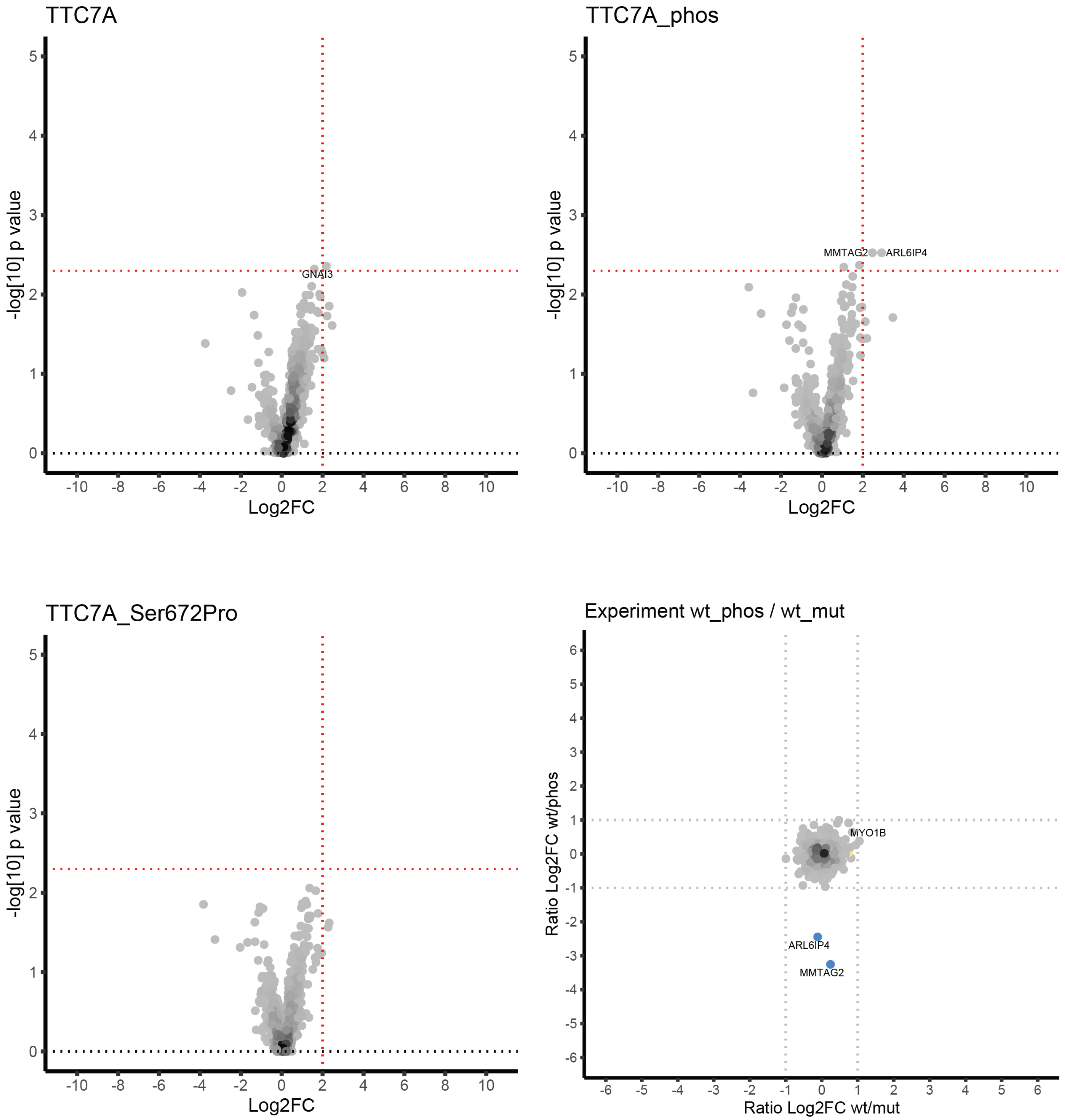

### TTF1: TLAMPEGpSQAGREAG

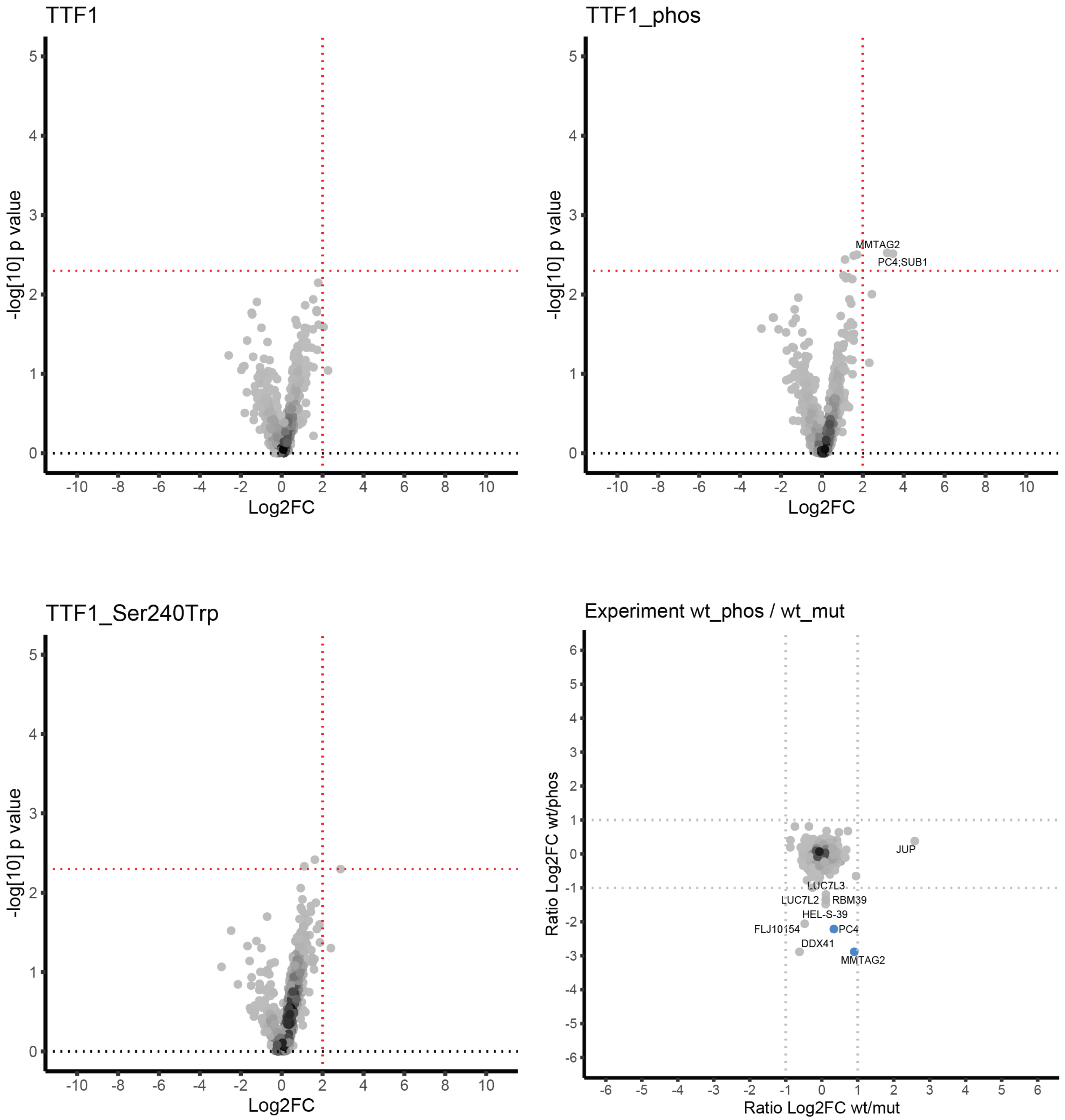

### TTR: ASGKTSEpSGELHGLT

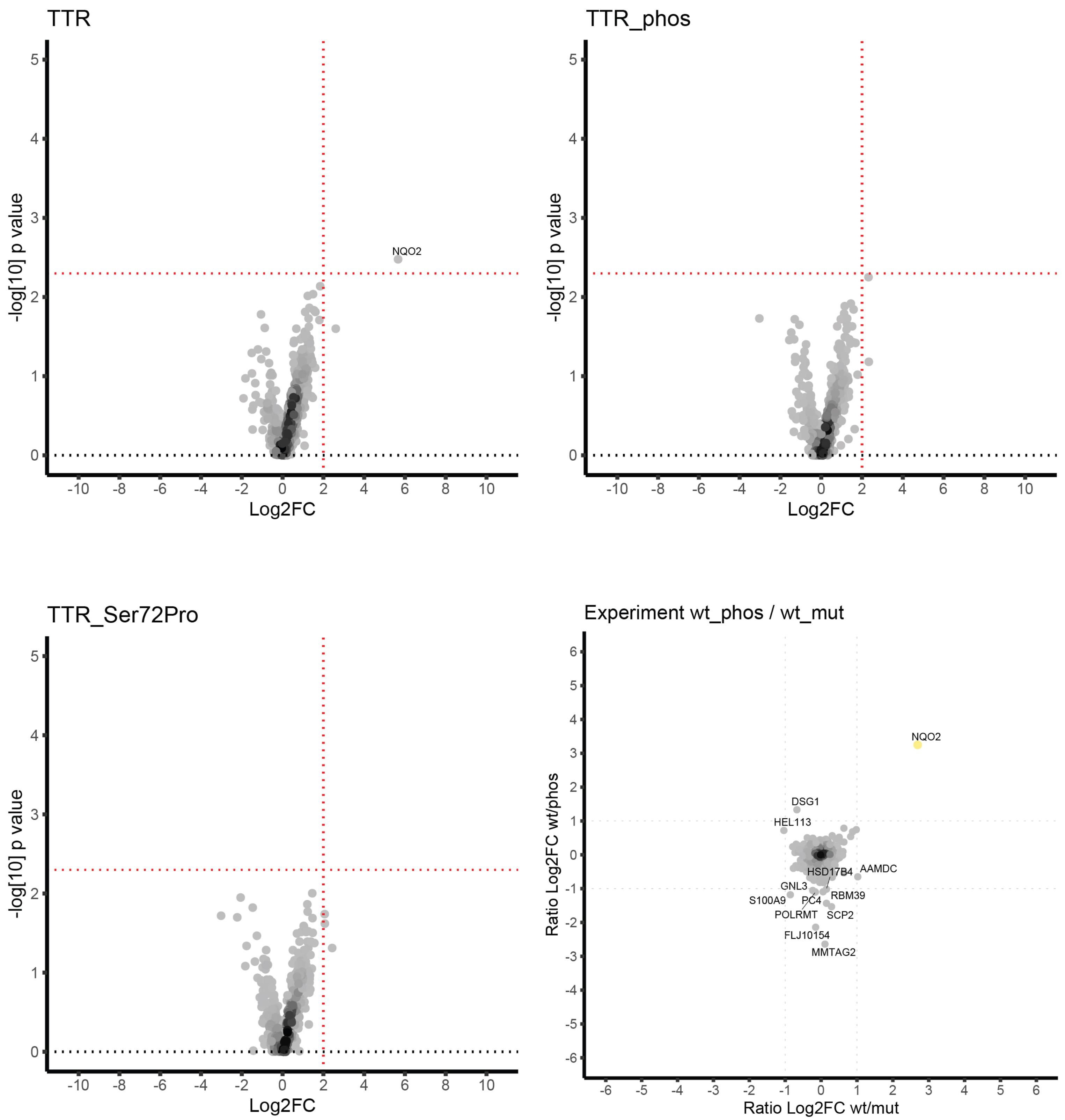


**Figure S3.** Volcano and SILAC scatter plots for all the peptide candidates: LFQ values were used to perform a Wilcoxon test where we compared proteins identified in one peptide form with all the proteins identified with all other WT and Mutant peptide forms. We treated each peptide form as a separate pull down to identify specific interactors. The calculated p values and fold changes(Fold change >2 and p values < 0.005) were plotted as Volcano plots (Top and bottom right). The log2 transformed SILAC ratios were plotted on scatter plots with wt/mut ratios on the x-axis and wt/phos ratios on the y-axis (bottom left) whereas the phos/mut ratios are not shown. SILAC ratios reveal differential interactors. To point out LFQ specific interactors in the scatter plots we used colored nods as follows: Blue; phospho-peptide specific, yellow; wt-peptide specific, red; mut-peptide specific, orange; wt and mut-peptide specific, green; wt and phospho-peptide specific, purple; mut and phospho-peptide specific, and pink; wt, mut and phosphor-peptide specific.

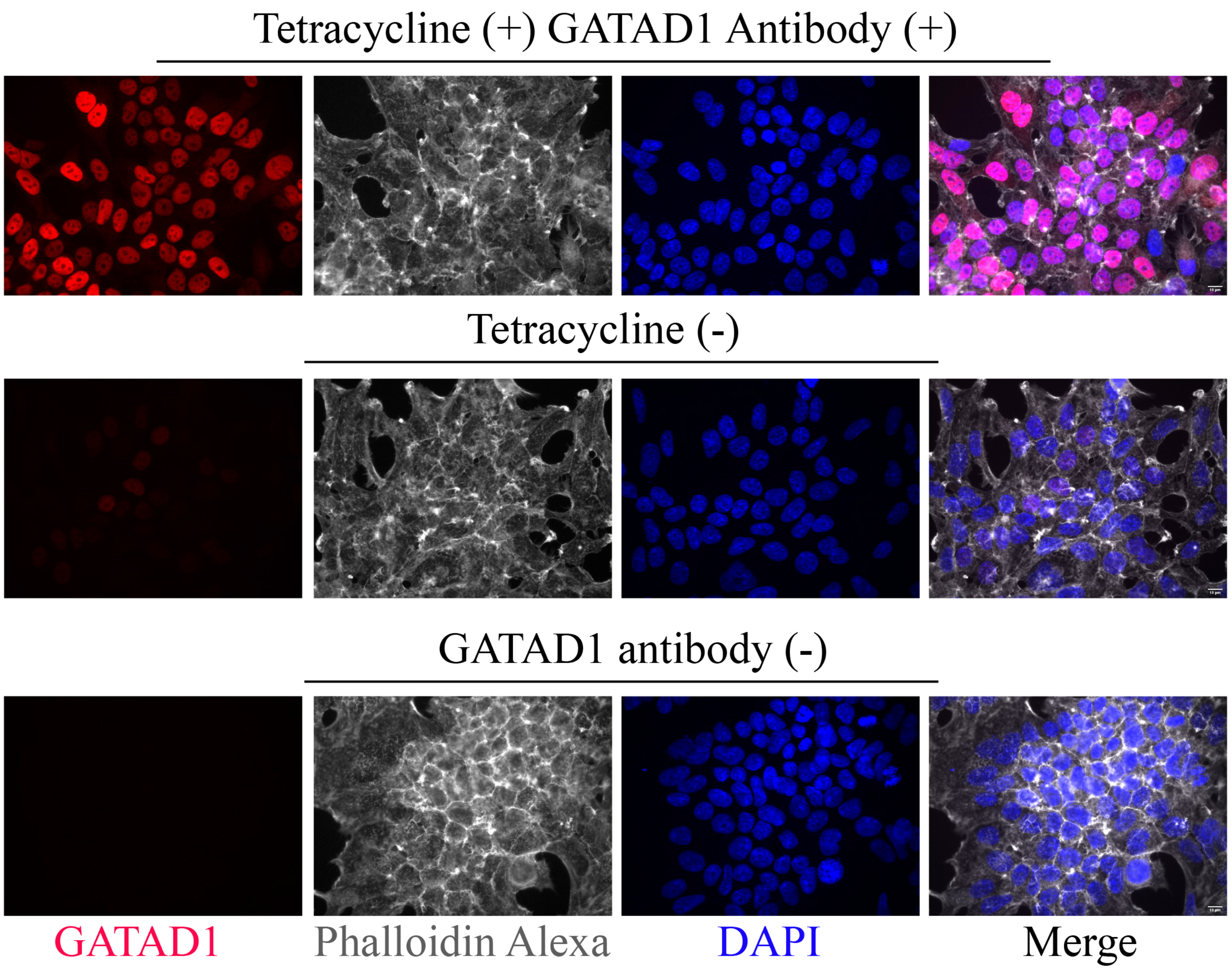


**Figure S4**. Immunofluorescence images of FlpIn-293 cells that express BirA*-FLAG-GATAD1 in a tetracycline inducible manner. The images display a nuclear localization of GATAD1 protein as expected, indicating that the BirA* and FLAG does not alter protein localization (Top lane). We observed little to no leakage of expression in the absence of 1 µg/µl of tetracycline (Middle lane) and no signal in the absence of primary GATAD1 antibody (Bottom lane).
